# Phasic pupil responses to melanopsin- and cone-directed stimulation remain stable across circadian time

**DOI:** 10.64898/2026.10.13.758787

**Authors:** Hannah Sophie Heinrichs, Diana Glebowicz, Manuel Spitschan

## Abstract

Human pupil responses vary with time of day, but whether this reflects circadian modulation of specific photoreceptor pathways or other time-dependent processes remains unclear. We investigated circadian variation in photoreceptor-directed pupil responses using silent substitution combined with a 40-hour ultra-short sleep–wake protocol designed to limit the progressive accumulation of homeostatic sleep pressure. Twelve healthy adults completed the protocol, of whom eight met the quality-control criteria for analysis. Repeated pupillometry measured responses to 0.4-Hz modulations targeting broadband flux, melanopsin, luminance, L–M cone-opponent, and S-cone pathways. Salivary melatonin provided participant-specific period estimates. We separated slow tonic constriction from phasic pupil tracking and compared hierarchical circadian cosinor models with constant-level models. Across all five stimulus conditions, constant-level models were favoured for phasic amplitude (log_10_ BF_10_ = −3.36 to −1.10). Constant-level models were also favoured for tonic minimum in four conditions, whereas L–M favoured the circadian model (log_10_ BF_10_ = 0.67), although the evidence was numerically imprecise. These findings suggest that phasic pupil tracking of photoreceptor-directed modulation remains relatively stable across the sampled protocol, despite previously reported time-of-day variation in other pupil metrics. They indicate that circadian influences on the pupillary light response may depend on the temporal component and stimulus conditions examined, rather than reflecting a uniform change in photoreceptor-driven responsiveness. Smaller, non-sinusoidal, or adaptation-dependent effects remain possible.

**Impact statement:** Measuring pupil responses under a protocol designed to separate circadian phase from accumulating sleep pressure constrains retinal explanations for reported time-of-day variation in the pupillary light response under the tested conditions.

## Introduction

The pupillary light response provides an accessible readout of retinal input to a non-image-forming visual pathway. Pupil diameter is controlled by signals originating in rods, cones, and melanopsin-expressing intrinsically photosensitive retinal ganglion cells (ipRGCs) (***Spitschan, 2019***). The ipRGCs combine their intrinsic melanopsin response with extrinsic rod and cone input, a property demonstrated directly in primate retina (***Dacey et al., 2005***). They send signals to subcortical targets that include the olivary pretectal nucleus, which mediates pupil constriction, and the suprachiasmatic nucleus, which coordinates circadian timing (***Hattar et al., 2006***). The functional contribution of melanopsin-expressing retinal ganglion cells to sustained pupil constriction has also been demonstrated in macaques and humans (***Gamlin et al., 2007***). This shared retinal origin has made pupillometry attractive as a non-invasive assay of photoreceptor contributions to non-image-forming vision. However, the pupillomotor and retino-hypothalamic pathways are not identical, and pupil measurements should not be treated as a direct assay of circadian light responses. The measured contribution of each photoreceptor pathway also depends on the stimulus and response component. Cone signals contribute prominently to rapid constriction and phasic tracking of temporal modulation, whereas melanopsin has slower temporal dynamics and contributes strongly to sustained constriction and the post-illumination pupil response (***Adhikari et al., 2015; Joyce et al., 2015; Spitschan et al., 2014***). Pupillometric outcomes therefore provide pathway-weighted measurements rather than interchangeable measures of a single underlying sensitivity.

Daily variation in pupil size and light-reflex dynamics was investigated decades before melanopsin was identified, but the early literature did not converge on a consistent circadian pattern. ***Döring and Schaefers (1950***) reported a diurnal rhythm in resting pupil diameter, and ***Tiedt (1963***) later described a biphasic 24-hour pattern in the kinetics of the pupillary light reflex, with generally faster responses during the day and slower responses at night. Subsequent findings were mixed. In a 24-hour study conducted under controlled illumination, Loving and colleagues found that the estimated peak phases of pupil diameter were distributed across the day rather than clustering at a common circadian phase (***Loving et al., 1996***). Dark-adapted pupillography during 30 hours of wakefulness likewise revealed non-monotonic daytime changes in spontaneous pupil instability and diameter (***Wilhelm et al., 2001***). More recently, steady-state pupil size varied during a 29-hour constant-routine protocol, whereas pupil-size variability did not; neither measure was associated with subjective sleepiness or electroencephalographic alertness (***Van Egroo et al., 2019***). Together, these studies establish a long-standing basis for investigating daily pupil variation, while showing that its direction, timing, and physiological interpretation depend strongly on the measured outcome and the experimental protocol.

Contemporary studies have extended this question to light-evoked responses associated with melanopsin and cone function, but the source of the reported variation remains unresolved. In a 24-hour protocol, ***Zele et al. (2011***) reported variation in the post-illumination pupil response and interpreted it as modulation of ipRGC-mediated function. ***Münch et al. (2012***) similarly found that a post-stimulus response following blue light covaried with melatonin, whereas the response following red light was associated with subjective sleepiness. During a constant-routine protocol, ***Daguet et al. (2019***) identified both a rhythmic component and a wake-dependent trend in baseline pupil diameter, and later work reported circadian variation in responses to short- and long-wavelength stimuli (***Daguet et al., 2024***). These studies differ in stimulus duration, wavelength, adaptation state, and outcome definition, including baseline diameter, maximum constriction, redilation, and the post-illumination response. Moreover, monochromatic stimuli change the excitation of several photoreceptor classes simultaneously. The findings therefore support the possibility that retinal or downstream pupillomotor responsiveness changes across circadian phase, but they do not identify whether that variation originates in melanopsin, cone, or post-receptoral pathways, or in mechanisms that influence pupil control more generally.

A central interpretive problem is that circadian phase and homeostatic sleep pressure usually covary in protocols that sample an entire biological day. The two-process model distinguishes an endogenous circadian process from a homeostatic process that increases during wakefulness and declines during sleep (***Borbély, 1982; Borbély et al., 2016***). Under ordinary sleep–wake schedules, clock time, circadian phase, prior light exposure, behaviour, and time awake also change together. Constant-routine protocols reduce masking by controlling factors such as posture, illumination, activity, and food intake, but they typically require sustained wakefulness. Sleep pressure therefore accumulates across the same interval in which circadian phase is measured. Statistical separation of a rhythmic component from a wake-dependent trend can be informative, but it depends on assumptions about the form and additivity of those processes (***Daguet et al., 2019, 2024***). Consequently, an observed time-of-day difference or fitted circadian pattern in pupil diameter, constriction, or redilation cannot by itself identify a circadian change in photoreceptor-driven responsiveness. A protocol that distributes sleep opportunities across circadian phase can reduce, although not eliminate, this source of confounding.

A second interpretive problem is that conventional chromatic stimuli do not isolate the photoreceptor pathway responsible for a pupil response. Narrow-band blue and red lights differ in their relative activation of melanopsin and cones, but each stimulates several photoreceptor classes because their spectral sensitivities overlap. Responses to such stimuli are therefore melanopsin-weighted or cone-weighted rather than photoreceptor-specific (***Spitschan and Woelders, 2018***). Silent substitution addresses this limitation by changing the excitation of a targeted photoreceptor class, or a defined combination of classes, while holding excitation of the remaining classes nominally constant (***Estévez and Spekreijse, 1982; Tsujimura et al., 2010; Spitschan, 2019; Martin et al., 2023; Nugent et al., 2023; Huchzermeyer and Kremers, 2026***). Applied to pupillometry, this method can distinguish melanopsin-directed responses from cone-luminance and cone-opponent responses. Sinusoidal modulation adds a complementary temporal measure because the amplitude with which the pupil tracks the modulation quantifies the phasic response at the stimulus frequency, while slower changes in pupil diameter can be analysed separately as a tonic component (***Spitschan et al., 2014; Barrionuevo et al., 2014***). Spectral isolation and temporal decomposition together therefore permit a more specific test of where any circadian variation is expressed within the pupillomotor system, without inferring photoreceptor identity from stimulus wavelength alone.

The present study combines photoreceptor-directed stimulation with a protocol designed to separate circadian timing from systematic accumulation of sleep pressure. Participants completed repeated pupillometry during a 40-hour ultra-short sleep–wake forced-desynchrony protocol in which 2.5-hour wake episodes alternated with 1.25-hour sleep opportunities (***Heinrichs and Spitschan, 2025b***). This schedule distributes sleep and wake episodes across circadian phase and limits the progressive increase in homeostatic pressure that occurs during continuous wakefulness. Salivary melatonin was sampled throughout the protocol to estimate each participant’s circadian period and set the participant-specific period of the pupil model. Pupillary responses were measured using 0.4-Hz sinusoidal silent-substitution stimuli in five conditions: broadband flux modulation, achromatic L+M+S cone modulation, L–M cone-opponent modulation, S-cone-directed modulation, and melanopsin-directed modulation. The common frequency was selected as a practical compromise at which each targeted pathway was expected to produce a measurable response, while allowing all conditions to be presented repeatedly within each experimental block (following prior data on pupillary temporal transfer functions measured in ***Spitschan et al., 2014***).

We tested whether circadian phase modulates photoreceptor-directed pupil responses under these conditions. The main outcome reported here was the fundamental amplitude of the phasic pupil response at 0.4 Hz, which quantified how strongly the pupil tracked each modulation; the minimum of the tonic component provided a complementary outcome for slower response dynamics. For each stimulus condition, hierarchical Bayesian cosinor models incorporating participant-specific melatonin-derived periods were compared with constant-level models. The published protocol (***Heinrichs and Spitschan, 2025b***) instead specified maximum pupil-constriction amplitude for each condition normalised to the corresponding broadband FLUX response; the condition-specific, unnormalised phasic and tonic outcomes reported here are therefore revised rather than confirmatory analyses. The phasic amplitude is matched to the sinusoidal stimulus design, whereas maximum constriction is the natural summary of a light-pulse paradigm (***McAdams et al., 2018***); analysing phasic and tonic outcomes separately also keeps two separable response components apart (Appendix 1). The central question was whether the measured phasic and tonic responses were better represented by an approximately circadian sinusoid or by a constant level across the protocol.

## Results

Of the 453 participant-by-condition-by-block pupil traces that passed trial- and block-level quality control (see Materials and methods), 55 (spanning 19 participant-by-condition series) fell below the six-block coverage criterion and were excluded, leaving *N*_fits_ = 398 traces from eight participants for parameter extraction.

### Salivary melatonin rhythms provided participant-specific period estimates

Salivary melatonin concentrations varied across the 40-hour protocol, rising in the biological evening, peaking overnight, and returning to low daytime values, as illustrated for one participant in Figure 2A. The bi-harmonic cosinor described this time course with a mean *R*^2^ of 0.92 (*SD* = 0.04, range 0.84–0.97; Figure 2B). Estimated periods ranged from 23.90 to 24.65 h (*M* = 24.23 h, *SD* = 0.30 h). The fundamental amplitude averaged 10.17 ± 2.79 pg/mL.

**Figure 1.**
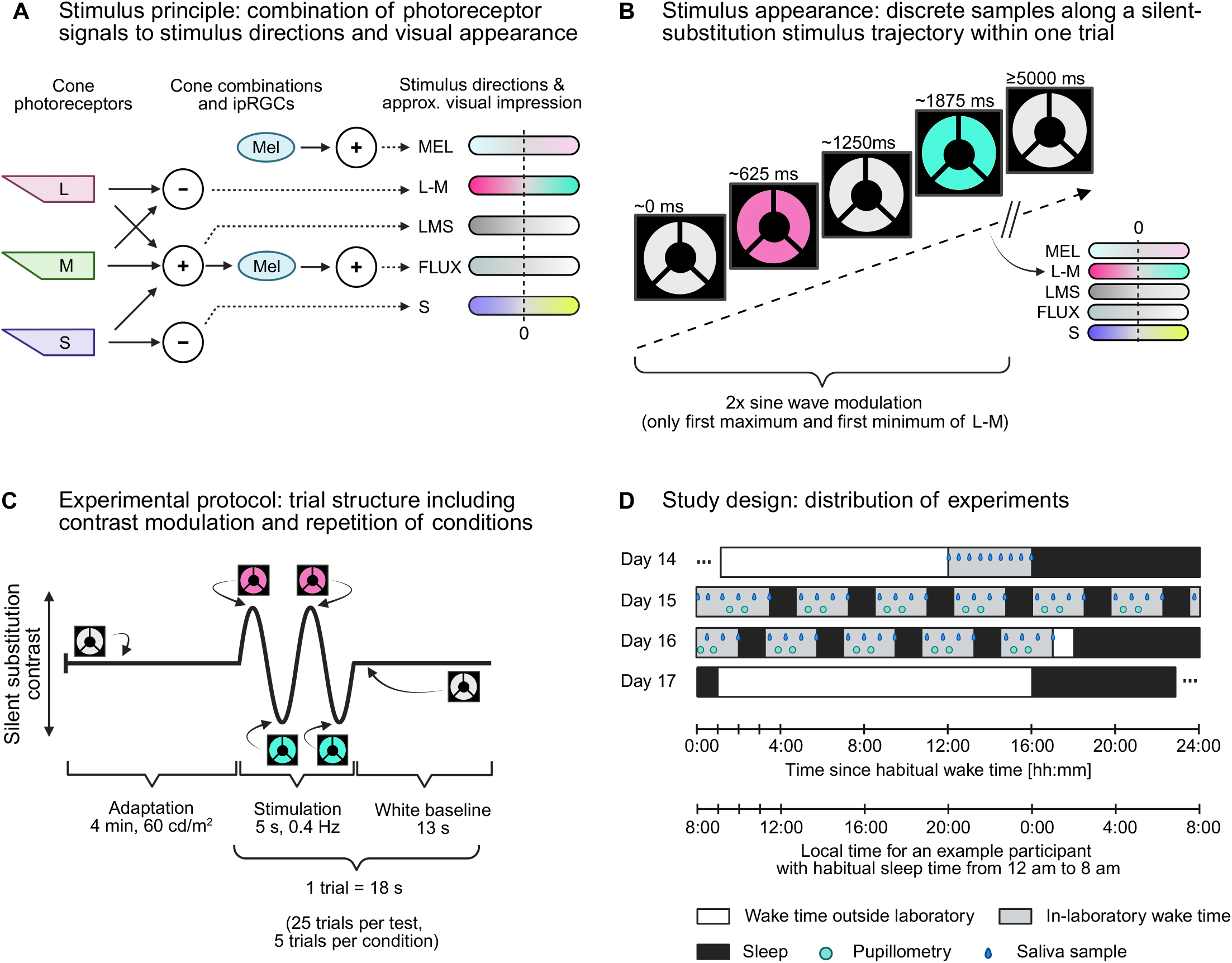
Study design and photoreceptor-directed stimulation. **A**, Retinal and post-receptoral mechanisms targeted by the stimulus directions. L, M and S denote cone signals and Mel melanopsin signals of intrinsically photosensitive retinal ganglion cells; + and ™ mark summing and opponent combinations. **B**, Approximate visual appearance of the L–M example. **C**, Trial sequence comprising adaptation, 0.4-Hz contrast modulation, and baseline periods. **D**, Dim-light evening and 40-h ultra-short sleep–wake protocol, with selected saliva and pupillometry measurements during scheduled wake episodes; the complete protocol is described by ***Heinrichs and Spitschan (2025b)***.

**Figure 2.**
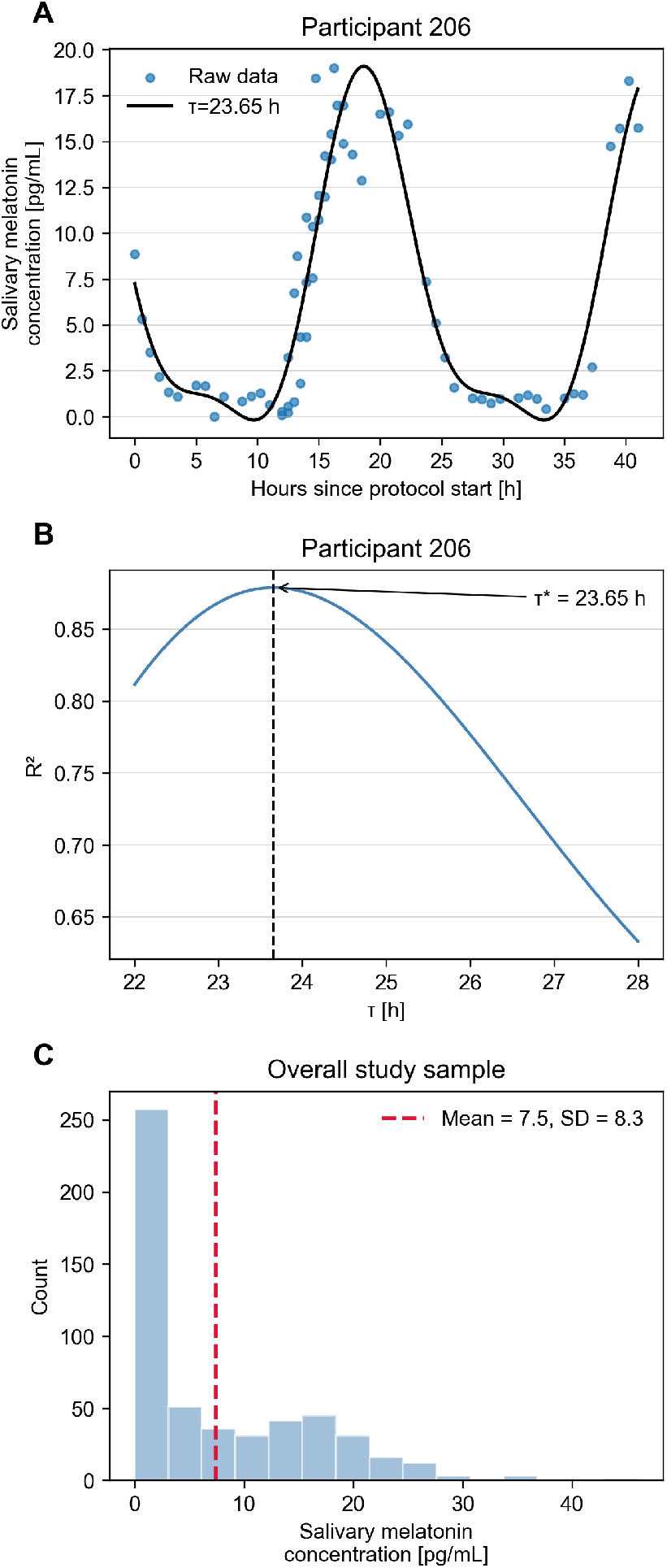
Participant-specific period estimation from salivary melatonin. **A**, Raw concentrations and the selected bi-harmonic fit for participant 206. **B**, *R*2 across candidate periods from 22 to 28 h for the same participant; the dashed line marks the selected period. **C**, Distribution of raw concentrations across the eight analysed participants; the dashed line marks the mean. Per-participant observation counts and fitted parameters are reported in Table 1.

Raw melatonin concentrations were right-skewed and included values below the measurement threshold during the subjective day. Across the 368 concentrations from the eight analysed participants, the mean was 6.41 pg/mL and the median was 2.09 pg/mL (Figure 2C).

**Table 1.**
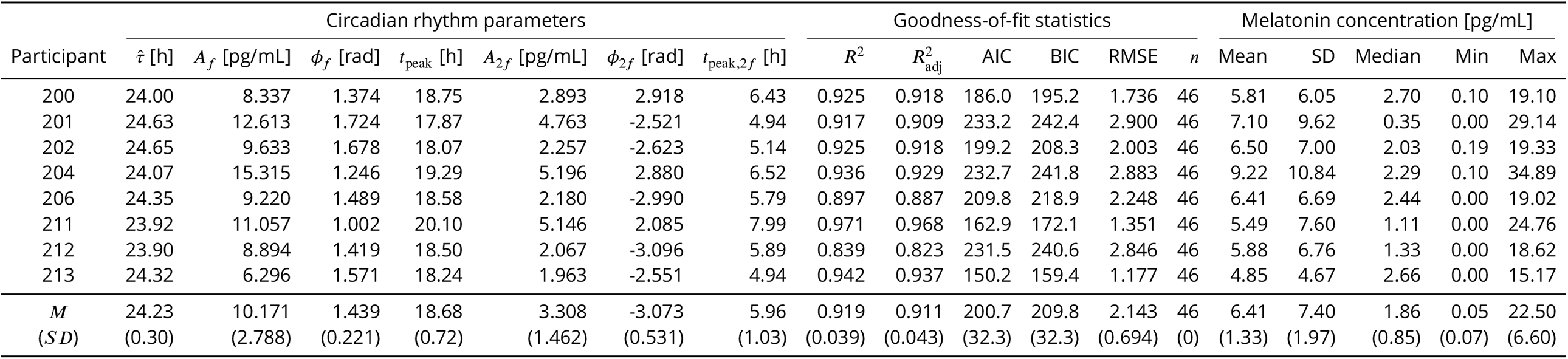
Best-fit period, circadian rhythm parameters, and goodness-of-fit statistics per participant. 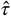 is the period selected by AIC from the 22–28 h scan; *A* and ϕ are the amplitude and phase of the fundamental (*f*) and second harmonic (2*f*) of the bi-harmonic cosinor, and *t*_peak_ is the corresponding peak time in hours from protocol start. *n* is the number of saliva samples, and the concentration columns summarise the raw values entering the fit. The final rows give the mean and SD across participants (circular mean and circular SD for ϕ).

| Participant | Circadian rhythm parameters |  |  |  |  |  |  | Goodness-of-fit statistics |  |  |  | Melatonin concentration [pg/mL] |  |  |  |  |  |  |
| --- | --- | --- | --- | --- | --- | --- | --- | --- | --- | --- | --- | --- | --- | --- | --- | --- | --- | --- |
| | $\hat{\tau}$ [h] | $A_f$ [pg/mL] | $\phi_f$ [rad] | $t_{\text{peak}}$ [h] | $A_{2f}$ [pg/mL] | $\phi_{2f}$ [rad] | $t_{\text{peak},2f}$ [h] | $R^2$ | $R^2_{\text{adj}}$ | AIC | BIC | RMSE | $n$ | Mean | SD | Median | Min | Max |
| 200 | 24.00 | 8.337 | 1.374 | 18.75 | 2.893 | 2.918 | 6.43 | 0.925 | 0.918 | 186.0 | 195.2 | 1.736 | 46 | 5.81 | 6.05 | 2.70 | 0.10 | 19.10 |
| 201 | 24.63 | 12.613 | 1.724 | 17.87 | 4.763 | -2.521 | 4.94 | 0.917 | 0.909 | 233.2 | 242.4 | 2.900 | 46 | 7.10 | 9.62 | 0.35 | 0.00 | 29.14 |
| 202 | 24.65 | 9.633 | 1.678 | 18.07 | 2.257 | -2.623 | 5.14 | 0.925 | 0.918 | 199.2 | 208.3 | 2.003 | 46 | 6.50 | 7.00 | 2.03 | 0.19 | 19.33 |
| 204 | 24.07 | 15.315 | 1.246 | 19.29 | 5.196 | 2.880 | 6.52 | 0.936 | 0.929 | 232.7 | 241.8 | 2.883 | 46 | 9.22 | 10.84 | 2.29 | 0.10 | 34.89 |
| 206 | 24.35 | 9.220 | 1.489 | 18.58 | 2.180 | -2.990 | 5.79 | 0.897 | 0.887 | 209.8 | 218.9 | 2.248 | 46 | 6.41 | 6.69 | 2.44 | 0.00 | 19.02 |
| 211 | 23.92 | 11.057 | 1.002 | 20.10 | 5.146 | 2.085 | 7.99 | 0.971 | 0.968 | 162.9 | 172.1 | 1.351 | 46 | 5.49 | 7.60 | 1.11 | 0.00 | 24.76 |
| 212 | 23.90 | 8.894 | 1.419 | 18.50 | 2.067 | -3.096 | 5.89 | 0.839 | 0.823 | 231.5 | 240.6 | 2.846 | 46 | 5.88 | 6.76 | 1.33 | 0.00 | 18.62 |
| 213 | 24.32 | 6.296 | 1.571 | 18.24 | 1.963 | -2.551 | 4.94 | 0.942 | 0.937 | 150.2 | 159.4 | 1.177 | 46 | 4.85 | 4.67 | 2.66 | 0.00 | 15.17 |
| $M$ | 24.23 | 10.171 | 1.439 | 18.68 | 3.308 | -3.073 | 5.96 | 0.919 | 0.911 | 200.7 | 209.8 | 2.143 | 46 | 6.41 | 7.40 | 1.86 | 0.05 | 22.50 |
| (SD) | (0.30) | (2.788) | (0.221) | (0.72) | (1.462) | (0.531) | (1.03) | (0.039) | (0.043) | (32.3) | (32.3) | (0.694) | (0) | (1.33) | (1.97) | (0.85) | (0.07) | (6.60) |

**Table 2.**
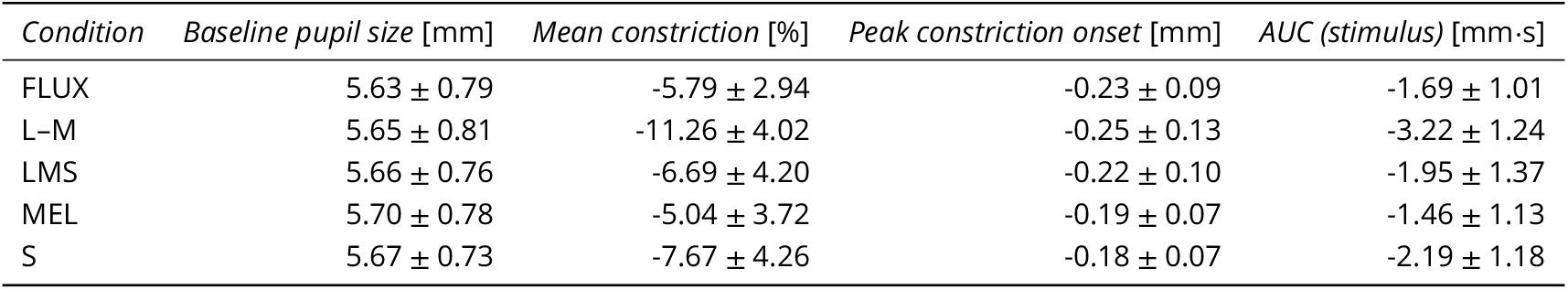
Descriptive pupil response metrics per chromatic condition. Values are mean ± SD across participants (averaged over blocks within participant). Baseline pupil size: mean absolute pupil diameter during 1 s pre-stimulus window. Mean constriction: mean percentage change from baseline during stimulus. Peak constriction onset: minimum baseline-corrected pupil size within 1 s of stimulus onset. AUC (stimulus): area under the baseline-corrected pupil trace over the stimulus window (trapezoidal rule).

| Condition | Baseline pupil size [mm] | Mean constriction [%] | Peak constriction onset [mm] | AUC (stimulus) [mm-s] |
| --- | --- | --- | --- | --- |
| FLUX | 5.63 ± 0.79 | -5.79 ± 2.94 | -0.23 ± 0.09 | -1.69 ± 1.01 |
| L-M | 5.65 ± 0.81 | -11.26 ± 4.02 | -0.25 ± 0.13 | -3.22 ± 1.24 |
| LMS | 5.66 ± 0.76 | -6.69 ± 4.20 | -0.22 ± 0.10 | -1.95 ± 1.37 |
| MEL | 5.70 ± 0.78 | -5.04 ± 3.72 | -0.19 ± 0.07 | -1.46 ± 1.13 |
| S | 5.67 ± 0.73 | -7.67 ± 4.26 | -0.18 ± 0.07 | -2.19 ± 1.18 |

Individual best-fit salivary melatonin periods and rhythm parameters, together with summary statistics of the raw melatonin values, are reported in Table 1. Melatonin results for the full study sample of 12 participants, of whom eight contributed to the pupillometry analysis here, are reported in Figure 2 of ***Heinrichs and Spitschan (2025a***).

### Photoreceptor-directed stimuli evoked tonic and phasic pupil responses

Pupil diameter changed during all five stimulus conditions (FLUX, L–M, LMS, MEL, and S). The averaged response profiles differed descriptively across conditions, whereas FLUX and LMS produced similar profiles (Figure 3).

**Figure 3.**
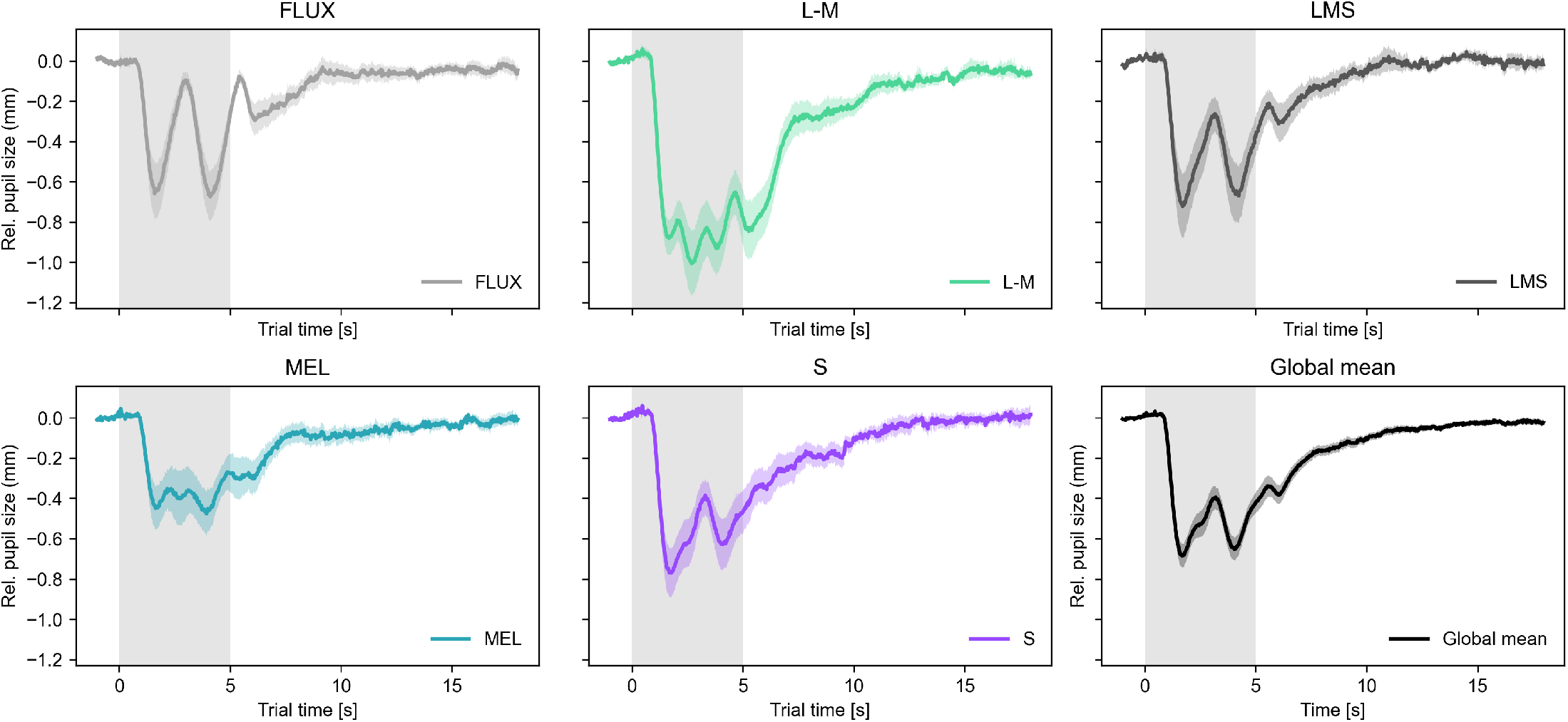
Average pupil responses differed across stimulus conditions. Lines and ribbons show the mean ± SEM across participants of baseline-corrected pupil diameter, after averaging each participant’s retained blocks at every time point. The final panel shows the mean ± SEM across all participant-by-condition averages; because each participant contributes up to five curves, its SEM is descriptive. The grey interval marks the 5-s stimulus. Retained trace counts were 83 FLUX (eight participants), 74 L–M (seven), 80 LMS (eight), 80 MEL (eight),

The two-stage decomposition described in Materials and methods produced tonic and phasic parameters for each of the *N*_fits_ = 398 retained traces (Table 3 and Table 4). Figure 4 illustrates the decomposition using per-condition averages across blocks and participants. The tonic component captured the slow constriction, and the cosinor captured oscillatory responses superimposed on it; participant-level examples and block-level traces are shown in ***Figure 4—figure Supplement 1*** and ***Figure 4—figure Supplement 2***.

**Table 3.** Tonic pupil component per chromatic condition. Values are mean ± SD across participants, with blocks averaged within participant; the *M* ± *SD* row averages these participant values across conditions. The tonic minimum is the most negative deflection of the low-pass filtered component within the 5-s stimulus window, and its latency is measured from stimulus onset.

| Condition | Tonic mean [mm] | Tonic minimum [mm] | Tonic minimum latency [s] |
| --- | --- | --- | --- |
| FLUX | -0.32 ± 0.20 | -0.45 ± 0.23 | 3.46 ± 0.40 |
| L-M | -0.66 ± 0.25 | -0.92 ± 0.35 | 3.36 ± 0.32 |
| LMS | -0.38 ± 0.27 | -0.53 ± 0.33 | 3.27 ± 0.33 |
| MEL | -0.29 ± 0.23 | -0.44 ± 0.29 | 3.23 ± 0.48 |
| S | -0.44 ± 0.23 | -0.60 ± 0.30 | 3.04 ± 0.48 |
| $M \pm SD$ | -0.41 ± 0.22 | -0.58 ± 0.28 | 3.26 ± 0.29 |

**Table 4.** Phasic pupil component per chromatic condition. Values are mean ± SD across participants, with blocks averaged within participant; the *M* ± *SD* row averages these participant values across conditions. *A*_*f*_ /*A*_2*f*_ is computed for each block and then averaged in the same way, so it is not the ratio of the displayed *A*_*f*_ and *A*_2*f*_ means. The fundamental (*f*) and second-harmonic (2*f*) components correspond to 0.4 and 0.8 Hz. Phases ϕ are summarised by the circular mean and circular SD at every averaging step. α and β are the sine and cosine slopes of the cosinor fit. Adjusted *R*^2^ refers to the bi-harmonic cosinor fitted to the residual after subtraction of the tonic component.

| Condition | Fundamental |  |  |  | 2nd harmonic |  |  |  | Ratio | Fit |
| --- | --- | --- | --- | --- | --- | --- | --- | --- | --- | --- |
| | $A_f$ [mm] | $\phi_f$ [rad] | $\alpha_f$ | $\beta_f$ | $A_{2f}$ [mm] | $\phi_{2f}$ [rad] | $\alpha_{2f}$ | $\beta_{2f}$ | $A_f/A_{2f}$ | adj. $R^2$ |
| FLUX | 0.25 ± 0.12 | -1.19 ± 0.20 | 0.23 ± 0.11 | 0.09 ± 0.07 | 0.05 ± 0.03 | 1.60 ± 0.51 | -0.04 ± 0.03 | -0.01 ± 0.02 | 7.52 ± 3.11 | 0.75 ± 0.16 |
| L-M | 0.12 ± 0.04 | -1.11 ± 1.00 | 0.09 ± 0.07 | 0.03 ± 0.03 | 0.10 ± 0.04 | 2.09 ± 0.34 | -0.08 ± 0.02 | -0.06 ± 0.05 | 1.49 ± 0.90 | 0.43 ± 0.09 |
| LMS | 0.22 ± 0.10 | -1.34 ± 0.24 | 0.21 ± 0.10 | 0.05 ± 0.05 | 0.06 ± 0.04 | 1.59 ± 0.56 | -0.05 ± 0.04 | -0.01 ± 0.02 | 4.53 ± 1.22 | 0.70 ± 0.18 |
| MEL | 0.10 ± 0.03 | -1.13 ± 0.37 | 0.07 ± 0.03 | 0.03 ± 0.03 | 0.06 ± 0.04 | 1.55 ± 0.47 | -0.05 ± 0.04 | -0.01 ± 0.02 | 3.18 ± 2.07 | 0.45 ± 0.15 |
| S | 0.19 ± 0.08 | -1.52 ± 0.23 | 0.18 ± 0.08 | 0.01 ± 0.04 | 0.06 ± 0.03 | 1.68 ± 0.31 | -0.06 ± 0.03 | -0.01 ± 0.02 | 4.79 ± 3.71 | 0.62 ± 0.14 |
| $M \pm SD$ | 0.18 ± 0.06 | -1.25 ± 0.24 | 0.16 ± 0.06 | 0.04 ± 0.04 | 0.07 ± 0.03 | 1.70 ± 0.36 | -0.05 ± 0.03 | -0.02 ± 0.02 | 4.36 ± 1.22 | 0.60 ± 0.11 |

**Figure 4.**
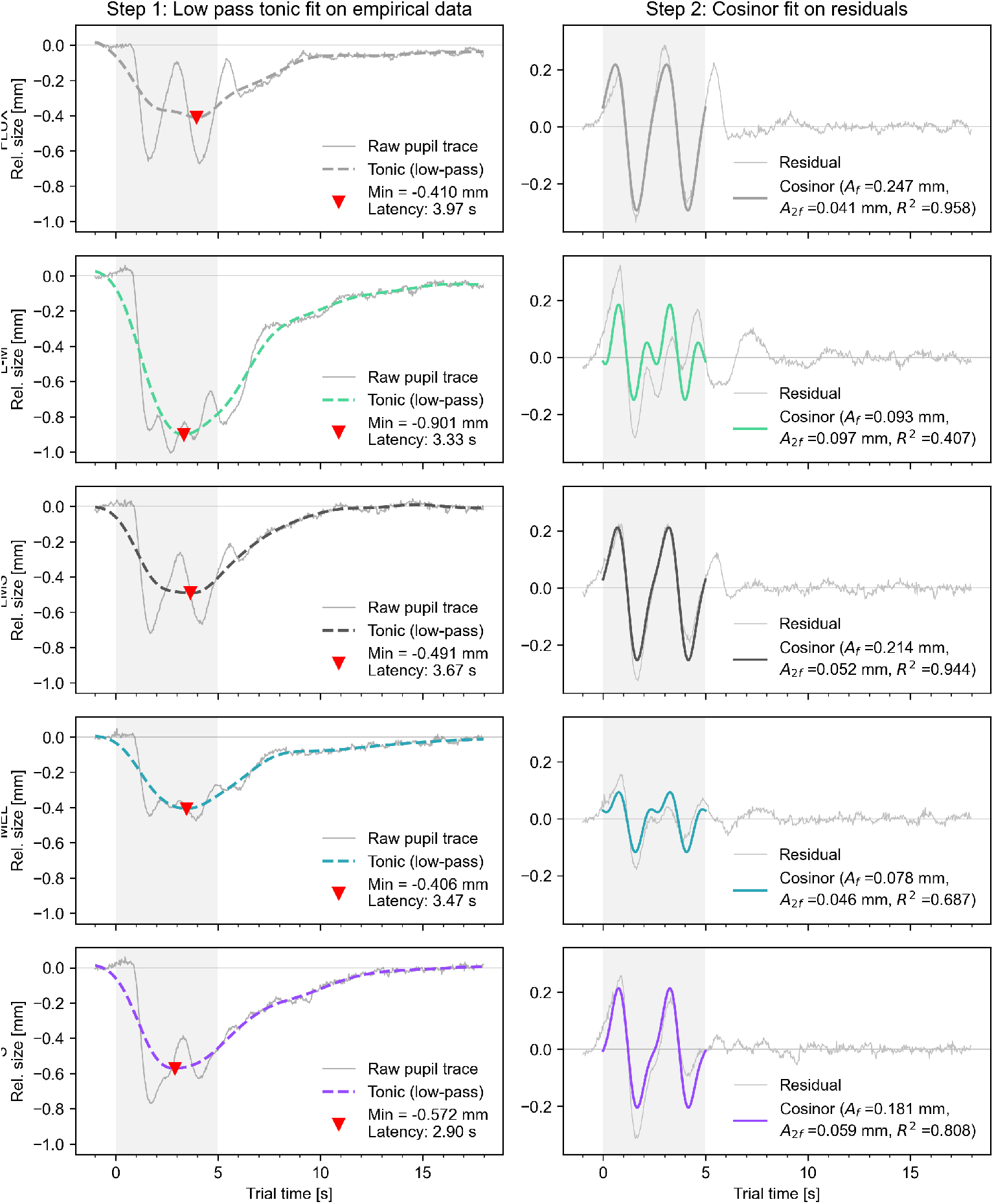
Two-step decomposition separated tonic and phasic response components. For each stimulus condition, left panels show the averaged baseline-corrected pupil trace and zero-phase low-pass component; right panels show the residual and fitted 0.4- and 0.8-Hz cosinor. The grey interval marks the 5-s stimulus. The visualisation uses retained traces averaged across trials, blocks, and participants. Figure supplements 1–5 provide participant examples and extended descriptive checks. **Figure 4—figure supplement 1**. Two-step decomposition for participant 202. **Figure 4—figure supplement 2**. Block-level response traces for participant 201. **Figure 4—figure supplement 3**. Diagnostics for the two-step decomposition: tonic mean-minimum correlation and phasic model fit. **Figure 4—figure supplement 4**. Extracted pupil parameters across protocol blocks. **Figure 4—figure supplement 5**. Extracted pupil parameters for participant 202.

The phasic fundamental amplitude was the main outcome reported in the subsequent circadian analysis, and tonic minimum was the complementary outcome. Group- and participant-level summaries of the additional derived parameters are shown in ***Figure 4—figure Supplement 4*** and ***Figure 4—figure Supplement 5***.

#### Tonic constriction varied across stimulus conditions

The low-pass filtered signal captured the slow, sustained pupil constriction evoked by the condition-specific contrast modulation. When averaging across all conditions and blocks, the tonic minimum was −0.58 mm relative to the mean pre-stimulus baseline pupil diameter, occurring at a mean latency of 3.26 s after formal stimulus onset. Across the retained block-level traces, tonic minimum and tonic mean were descriptively strongly correlated within every condition (*r* = 0.98 to *r* = 0.99); these pooled correlations do not account for observations nested within participants (***Figure 4— figure Supplement 3***). The minimum of the low-pass-filtered, trial-averaged pupil trace during the stimulus window was used as the tonic minimum for the respective block.

Tonic constriction depth differed descriptively across chromatic conditions (Table 3). The L–M cone-opponent modulation produced the deepest mean sustained constriction (-0.92 mm), whereas MEL and FLUX produced shallower responses (-0.44 and -0.45 mm, respectively). The ordering could differ between participants.

Latency varied little across conditions, from 3.04 s (S) to 3.46 s (FLUX) after onset (Table 3). For every stimulus direction, the mean latency placed the tonic minimum in the second half of the 5-s stimulus window and before stimulus offset.

#### Phasic pupil tracking was strongest for broadband flux and luminance modulation

The phasic residual (raw trace minus tonic component) was fitted over the 5-s stimulus window with a bi-harmonic cosinor at the 0.4-Hz stimulus frequency and its second harmonic. Fit quality differed between participants and conditions (***Figure 4—figure Supplement 3***). Each bi-harmonic fit estimates only a few regression coefficients from the pupil size samples within each 5 s stimulus window, so the correction for model complexity is negligible and the adjusted *R*^2^ is almost identical to the unadjusted value. We therefore report only the adjusted value. Adjusted *R*^2^ was low for L–M and MEL (mean 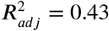 and 0.45, respectively), and highest for FLUX, near 0.8 (mean 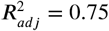)On average, the fundamental plus second-harmonic cosinor accounted for the majority of variance in the residual trace of the FLUX, LMS, and S conditions (mean 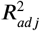 ranging from 0.62 to 0.75), but a considerable amount of unexplained variance remained for MEL and L–M (mean 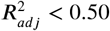). This mirrors the differences in fundamental amplitude *A*_*f*_.

The fundamental amplitude *A*_*f*_ quantifies the approximate magnitude of the sinusoidal pupil tracking response. Mean amplitudes per condition were: FLUX 0.25 mm, L–M 0.12 mm, LMS 0.22 mm, MEL 0.10 mm, and S 0.19 mm. The broadband FLUX and LMS modulations produced the largest average tracking amplitudes, whereas MEL and L–M produced lower amplitudes; S was intermediate (Figure 4). Response patterns nevertheless varied between participants.

The reported mean second-harmonic amplitude (*A*_2*f*_) was smaller than *A*_*f*_ in every condition, although the relative contribution varied by stimulus (Table 4). The FLUX response was dominated by the 0.4-Hz fundamental (*A*_*f*_ /*A*_2*f*_ = 7.52), whereas the L–M response contained a comparatively large 0.8-Hz component (*A*_*f*_ /*A*_2*f*_ = 1.49). This doubled-frequency component is visible in the grand-average response (Figure 4).

Across the 40-h protocol, empirical parameters differed in variability and time course by parameter and condition. Visual inspection did not reveal one common temporal pattern across outcomes or conditions.

The oscillatory amplitude at 0.4 Hz appeared broadly stable across blocks within each condition, with condition differences mainly in level rather than temporal profile. Tonic minimum varied across blocks; its model-based temporal pattern is reported below. The second-harmonic amplitude *A*_2*f*_ showed no systematic trend. Across blocks, only the L–M condition consistently had a higher second-harmonic amplitude than the other conditions, which clustered at lower values. Across blocks and conditions, baseline pupil size varied moderately, with condition means of 5.63–5.70 mm (Table 2). Phase parameters at *f* and 2*f* showed condition-specific offsets and appeared largely stable across blocks (Figure 4).

### Model comparison favoured constant phasic pupil tracking across circadian phase

We fitted a hierarchical cosinor model (M1) to the derived phasic amplitudes and compared it with a constant-level model containing participant-specific intercepts (M0). This comparison was applied separately to all five chromatic conditions using Sequential Monte Carlo sampling.

Observation counts ranged from 74 (L–M, 7 participants) to 83 (FLUX, 8 participants); per-condition values appear in Table 5. Positive log_10_ BF_10_ values favour M1, and negative values favour M0.

**Table 5.** Model comparison and population-level circadian estimates for the phasic fundamental amplitude, per chromatic condition. *N*_obs_, retained block-level observations; *N*_part_, participants; log_10_ BF_10_, log Bayes factor (base 10) comparing the circadian model M1 with the constant-level model M0. *A*_circ_ [mm] and ϕ_circ_ [rad] are the amplitude and phase of the fitted circadian modulation of the phasic fundamental amplitude, derived from the population-level posterior sine and cosine slopes and reported as posterior mean (SD); for the phase, the circular mean and circular SD of the posterior draws. This SD is group-level estimation uncertainty only; across-participant variability is given by the random-effect SDs in Table 1.

| Condition | $N_{\text{obs}}$ | $N_{\text{part}}$ | $\log_{10} \text{BF}_{10}$ | $A_{\text{circ}}$ [mm] | $\phi_{\text{circ}}$ [rad] |
| --- | --- | --- | --- | --- | --- |
| FLUX | 83 | 8 | -3.356 | 0.021 (0.010) | -2.209 (0.841) |
| L-M | 74 | 7 | -1.815 | 0.017 (0.007) | -3.047 (0.690) |
| LMS | 80 | 8 | -1.561 | 0.033 (0.012) | 3.093 (0.442) |
| MEL | 80 | 8 | -2.206 | 0.011 (0.005) | 2.493 (1.056) |
| S | 81 | 8 | -1.099 | 0.026 (0.008) | 2.488 (0.363) |

Across all five conditions, Bayes factors favoured M0 over M1 (Table 5): FLUX, -3.36; L–M, -1.81; LMS, -1.56; MEL, -2.21; and S, -1.10. These values are evidence for the specified constant-level model relative to the specified cosinor hierarchical model under the stated priors; they do not establish that every form or size of temporal variation is absent.

The numerical precision ratio exceeded the pragmatic threshold of 0.10 for LMS (0.11) and S (0.12), so the magnitudes of these two Bayes factors are imprecise (Appendix 2—table 2).

#### Magnitude of circadian fixed effects

Fixed-effect sine (*α*) and cosine (*β*) slopes had 95% credible intervals including zero in every condition except LMS (*β* = −0.031, CI [−0.058; −0.005]) and S (*β* = −0.020, CI [−0.035; −0.004]), where the cosine slope was narrowly negative. The Bayes factors nevertheless favoured M0 in both conditions. Because the derived circadian amplitude is constrained to be non-negative and phase is poorly identified when sine and cosine slopes approach zero, we treat the derived amplitude and phase summaries as descriptive rather than independent evidence of rhythmicity (Table 5). Full posterior parameter estimates are reported in Appendix 2—table 1.

#### Explained variance by the circadian model

Conditional Bayesian *R*^2^ (fixed plus random effects) ranged from 0.34 for MEL to 0.79 for FLUX, whereas marginal *R*^2^ for the fixed circadian terms ranged from 0.04 to 0.17. Conditional *R*^2^ was similar between M1 and M0. Thus, most explained variance reflected stable between-participant differences in mean amplitude rather than the fixed circadian terms (Figure 5). Residual-normality checks indicated departures for the LMS and S conditions; condition-specific diagnostic plots appear in Figure 5—figure supplements 3–7, with numerical summaries in Appendix 2—table 2.

**Figure 5.**
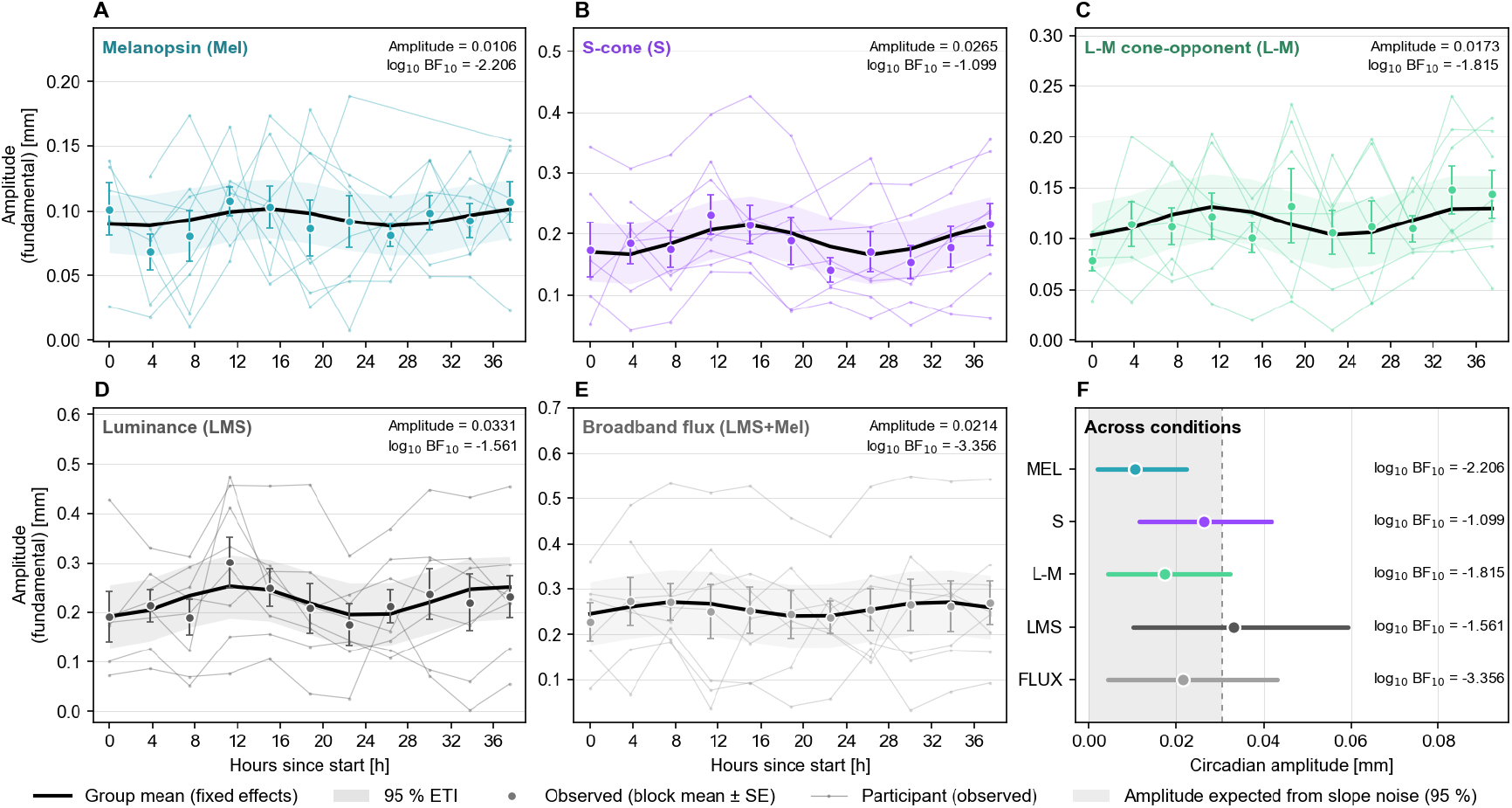
Phasic pupil tracking across nominal protocol time. **A–E**, Participant trajectories (thin lines), block means ± SEM (points and error bars), and M1 fixed-effect posterior means with 95% equal-tailed credible intervals (black lines and ribbons) for each condition. **F**, Derived fixed-effect circadian amplitude by condition, with 95% equal-tailed intervals of the posterior amplitude draws, and the corresponding log10 BF10. Negative log10 BF10 values favour the constant-level model. Because the amplitude is non-negative by construction, its interval cannot include zero; evidence for a rhythm is assessed by the Bayes factor. Figure supplements 1–2 show participant-level fits, and figure supplements 3–7 show condition-specific diagnostics. **Figure 5—figure supplement 1**. Participant-level phasic-amplitude fits for broadband flux. **Figure 5—figure supplement 2**. Participant-level phasic-amplitude fits for melanopsin-directed modulation. **Figure 5—figure supplement 3**. Phasic-amplitude diagnostics for broadband flux. **Figure 5—figure supplement 4**. Phasic-amplitude diagnostics for L–M modulation. **Figure 5—figure supplement 5**. Phasic-amplitude diagnostics for luminance modulation. **Figure 5—figure supplement 6**. Phasic-amplitude diagnostics for melanopsin-directed modulation. **Figure 5—figure supplement 7**. Phasic-amplitude diagnostics for S-cone-directed modulation.

### Evidence for circadian modulation of tonic constriction was limited and imprecise The complementary outcome was the tonic minimum–the most negative deflection of the low-pass-filtered pupil trace during the stimulus period. The same model comparison was applied to this outcome for each condition, with observation counts identical to those in the phasic analysis (Table 6)

The tonic results were more heterogeneous (Table 6). M0 was favoured for FLUX (log_10_ BF_10_ = -2.16), LMS (-3.56), MEL (-2.65), and S (-2.57). The L–M condition was the exception and favoured M1 (log_10_ BF_10_ = 0.67). Its numerical precision ratio was 0.17, above the pragmatic 0.10 threshold, so the magnitude of this Bayes factor is imprecise. This numerical imprecision is a reason for caution, but not by itself evidence against a biological effect; additional data and repeated marginal-likelihood estimation are needed.

**Table 6.** Model comparison and population-level circadian estimates for the tonic minimum, per chromatic condition. *N*_obs_, retained block-level observations; *N*_part_, participants; log_10_ BF_10_, log Bayes factor (base 10) comparing the circadian model M1 with the constant-level model M0. *A*_circ_ [mm] and ϕ_circ_ [rad] are the amplitude and phase of the fitted circadian modulation of the tonic minimum, derived from the population-level posterior sine and cosine slopes and reported as posterior mean (SD); for the phase, the circular mean and circular SD of the posterior draws. This SD is group-level estimation uncertainty only; across-participant variability is given by the random-effect SDs in Table 1.

| *Condition* | $N_{\text{obs}}$ | $N_{\text{part}}$ | $\log_{10} \text{BF}_{10}$ | $A_{\text{circ}}$ [mm] | $\phi_{\text{circ}}$ [rad] |
| --- | --- | --- | --- | --- | --- |
| FLUX | 83 | 8 | -2.163 | 0.075 (0.032) | 1.326 (0.576) |
| L-M | 74 | 7 | 0.672 | 0.178 (0.042) | -0.237 (0.283) |
| LMS | 80 | 8 | -3.564 | 0.061 (0.029) | -0.854 (0.898) |
| MEL | 80 | 8 | -2.646 | 0.066 (0.035) | -0.729 (1.317) |
| S | 81 | 8 | -2.571 | 0.081 (0.036) | -0.375 (0.747) |

#### Magnitude of circadian group effects

Fixed-effect sine (α) and cosine (β) slopes had 95% credible intervals including zero in the four M0-favouring conditions. For L–M, the cosine slope was positive (β = 0.167, 95% CI [0.086; 0.252]), whereas the sine slope remained compatible with zero (α = 0.041, 95% CI [−0.054; 0.134]). The derived amplitude was largest for L–M (0.18 mm), but it is non-negative by construction and is not treated as separate evidence of rhythmicity. Group-level fits are shown in Figure 6, participant-level examples in Figure 6—figure supplements 1 and 2, and full parameter estimates in Appendix 3— table 1.

**Figure 6.**
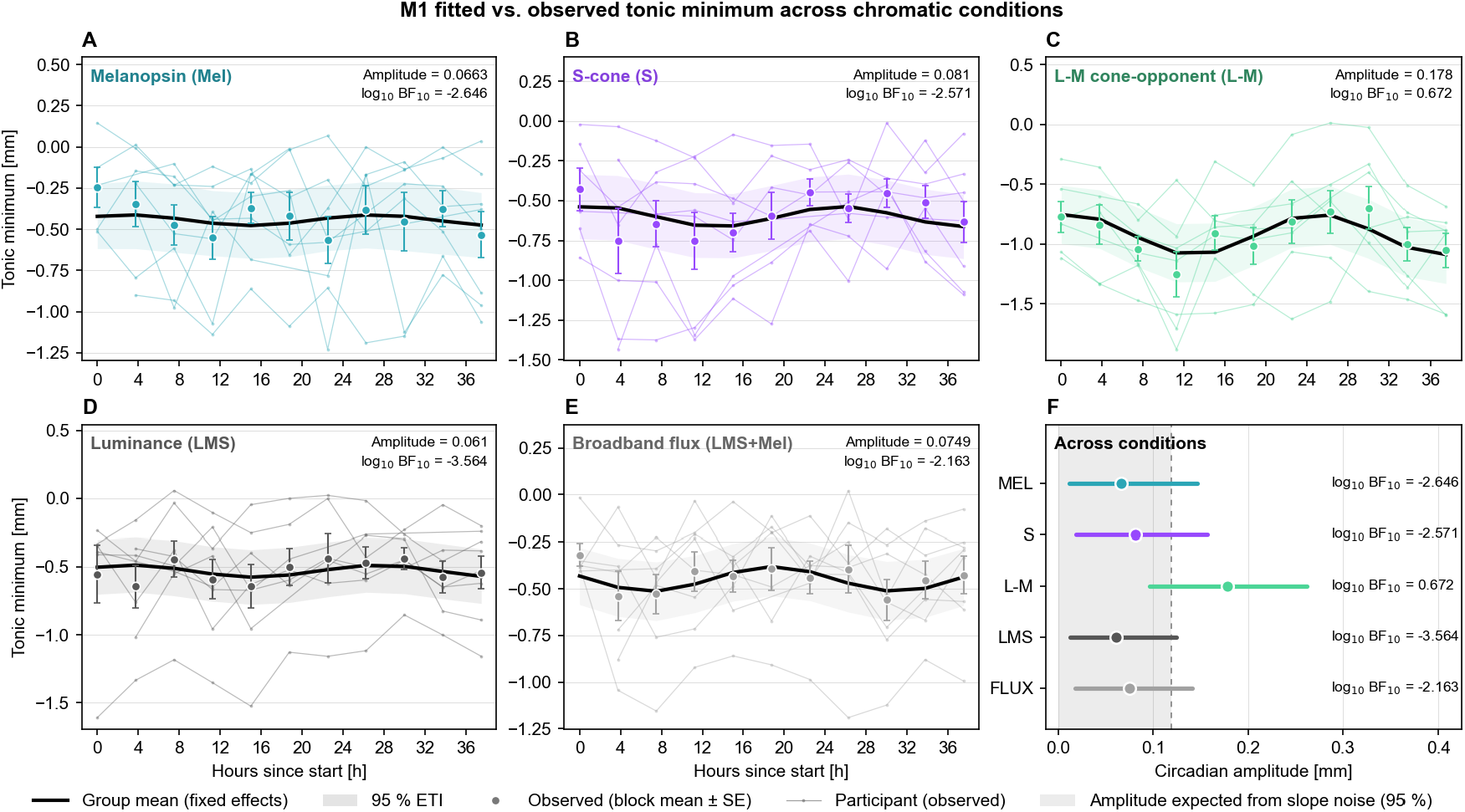
Tonic minimum across nominal protocol time. **A–E**, Participant trajectories (thin lines), block means ± SEM (points and error bars), and M1 fixed-effect posterior means with 95% equal-tailed credible intervals (black lines and ribbons) for each condition. **F**, Derived fixed-effect circadian amplitude by condition, with 95% equal-tailed intervals of the posterior amplitude draws, and the corresponding log10 BF10. The L–M condition favoured M1; the other conditions favoured M0. Because the amplitude is non-negative by construction, its interval cannot include zero; evidence for a rhythm is assessed by the Bayes factor. Figure supplements 1–2 show participant-level fits, and figure supplements 3–7 show condition-specific diagnostics. **Figure 6—figure supplement 1**. Participant-level tonic-minimum fits for L–M modulation. **Figure 6—figure supplement 2**. Participant-level tonic-minimum fits for melanopsin-directed modulation. **Figure 6—figure supplement 3**. Tonic-minimum diagnostics for broadband flux. **Figure 6—figure supplement 4**. Tonic-minimum diagnostics for L–M modulation. **Figure 6—figure supplement 5**. Tonic-minimum diagnostics for luminance modulation. **Figure 6—figure supplement 6**. Tonic-minimum diagnostics for melanopsin-directed modulation. **Figure 6—figure supplement 7**. Tonic-minimum diagnostics for S-cone-directed modulation.

#### Model diagnostics for the tonic outcome

Prior and posterior predictive checks, shrinkage, residual, and autocorrelation diagnostics were computed by condition. Numeric residual-normality checks indicated departures for MEL and S; condition-specific plots appear in Figure 6—figure supplements 3–7, with numerical summaries in Appendix 3—table 2.

## Discussion

The present study investigated whether phasic and tonic responses to photoreceptor-directed stimuli vary across circadian phase. Silent-substitution stimuli targeted broadband light flux modulation of all cone classes and melanopsin, achromatic luminance modulation, equally for all cones, nominal red-green chromatic cone-opponent modulation, melanopsin-directed modulation of ipRGCs, and approximate blue/violet-yellow chromatic, S-cone directed modulation, while an ultra-short sleep–wake protocol was designed to limit the systematic accumulation of homeostatic sleep pressure (***Heinrichs and Spitschan, 2025b***). For the revised main outcome, the constant-level model was favoured over the specified circadian model in all five conditions. Tonic minima also favoured the constant model in four conditions, whereas the L–M condition favoured the circadian model with an imprecisely estimated Bayes factor. Under the tested adaptation, contrast, temporal frequency, model, and priors, the principal result is therefore evidence for stable phasic pupil tracking rather than a general absence of circadian effects on photoreceptor sensitivity or pupil responses.

### Phasic response amplitudes and model fit across conditions

Fundamental response amplitudes differed across conditions. Broadband flux produced the largest mean fundamental amplitude, whereas melanopsin-directed modulation of ipRGCs (MEL) and nominal red-green chromatic cone-opponent modulation (L–M) produced the lowest. This pattern is compatible with known differences in temporal filtering across the pathways contributing to the pupil response at 0.4 Hz (***Spitschan et al., 2014; Joyce et al., 2015***), although response amplitude and model fit cannot distinguish photoreceptor sensitivity from downstream processing or residual measurement noise.

The comparatively low fit for L–M may partly reflect nonlinear opponent processing of L- and M-cone signals. In particular, cancellation at the fundamental and rectification could generate higher-order harmonics not fully captured by the bi-harmonic model; the present analysis does not directly test this mechanism.

The L–M condition produced the largest relative second-harmonic response and therefore the lowest *A*_*f*_ /*A*_2*f*_ ratio. Second harmonics have also been reported for pupil responses to stimuli involving L- and/or M-cones and attributed to nonlinear post-receptoral processing (***Spitschan et al., 2014; Barrionuevo and Cao, 2016***). The approximate blue/violet-yellow chromatic, S-cone directed modulation (S) and MEL conditions also contained small *A*_2*f*_ components, but the present design cannot determine whether these arose from pathway nonlinearities, residual photoreceptor contrast, or measurement noise. More generally, the L–M result shows that *A*_*f*_ alone does not capture all periodic structure in the response. The low *R*^2^ for L–M may therefore reflect both signal quality and mismatch between the fitted model and the response waveform.

### Model comparison favoured stable phasic tracking under the tested conditions

Across all five conditions, the Bayes factor favoured the constant-level model over the specified circadian model (Table 5). This is more informative than a failure to reject a null hypothesis, but it remains conditional on the model structures, data-dependent priors, and marginal-likelihood estimates used here. The low marginal *R*^2^ further indicates that the fixed circadian terms accounted for little variance beyond stable participant differences. The convergent direction across stimuli suggests that the finding was not confined to the weakest response condition, although numerical precision was limited for LMS and S.

Support for M0 was not confined to the lowest-amplitude responses: it also occurred for FLUX and LMS, which produced the largest mean fundamental amplitudes. A floor effect restricted to the weaker MEL and L–M responses is therefore an incomplete explanation for the common direction of model comparison, although the analysis does not establish the smallest circadian modulation that could have been resolved in each condition. Because the five comparisons used the same participants, apparatus, and background, they provide convergent measurements rather than independent replications.

#### Tonic minimum did not show the same pattern in every condition

For FLUX, LMS, MEL, and S, the tonic-minimum Bayes factor also favoured the constant-level model. The L–M condition was the exception (log_10_ BF_10_ = 0.67), and its fixed cosine slope was positive. Because the corresponding marginal-likelihood estimate was numerically imprecise and only seven participants contributed to this condition, the result requires replication. It should not be collapsed into the phasic null finding or dismissed solely because it differs from the other comparisons.

The contrast between the phasic and tonic comparisons also argues against treating pupil outcomes as interchangeable. Tonic minimum integrates the slowly varying response during the 5-s stimulus, whereas *A*_*f*_ isolates tracking at 0.4 Hz after removal of the low-pass component. If replicated, modulation confined to tonic minimum would implicate slower pupil dynamics rather than phasic tracking itself. In the present data, however, the numerical imprecision and small sample make this a hypothesis for replication rather than a demonstrated dissociation.

### Design differences may help explain divergence from earlier findings

The phasic findings differ from reports of time-of-day or diurnal variation in other pupil metrics. Differences in stimulus design, outcome definition, adaptation, and sleep–wake protocol may contribute to this divergence, but the present study cannot determine which difference is responsible.

#### Monochromatic vs. silent substitution stimuli

The earlier studies discussed here used narrow-band blue and red stimuli to weight ipRGC and cone responses. Such stimuli co-activate multiple photoreceptor classes because their spectral sensitivities overlap. By the principle of univariance, their responses cannot be uniquely attributed to one class (***Spitschan et al., 2017***). Silent-substitution stimuli are instead designed to modulate a targeted class or combination while nominally holding others constant, allowing a more directed comparison while remaining subject to observer variability and photoreceptor splatter.

Using a detrended outcome during prolonged wakefulness, ***Daguet et al. (2024***) reported circadian variation in post-illumination responses to red and blue light. Those wavelength-defined responses and the 0.4-Hz tracking measure used here probe different mixtures of pathways and temporal components. The results are therefore not direct replications and need not agree.

#### Sleep–wake protocol

Earlier studies obtained repeated measurements during prolonged wakefulness and used constant-routine controls or statistical detrending (***Daguet et al., 2019, 2024***). The present protocol instead interleaved sleep opportunities with wake episodes (***Heinrichs and Spitschan, 2025b***), limiting the systematic accumulation of sleep pressure. Because achieved sleep and residual alertness were not quantified in the materials available for this analysis, homeostatic and masking effects cannot be considered eliminated. The protocol difference may nevertheless help explain some divergence across studies.

Future studies should combine this schedule with objective sleep recording and contemporaneous alertness measures so that residual homeostatic and masking effects can be quantified rather than assumed to be absent.

### Pupil stability does not localise circadian gating

***Zele et al. (2011***) proposed that circadian modulation of ipRGC sensitivity could gate photic input to the SCN, and ***Daguet et al. (2024***) considered retinal and central mechanisms. Molecular work in mice supports gating in the retina and within the SCN (***Komal et al., 2025, 2026***). Circadian and pupillomotor functions also involve partly distinct ipRGC subpopulations and downstream targets (***Komal et al., 2026; Chen et al., 2011***). A pupil measurement therefore cannot be assumed to report gating in the retino-hypothalamic pathway.

The stable phasic response to the melanopsin-directed stimulus is compatible with gating else-where in the system, but it does not demonstrate where gating occurs. Likewise, local retinal clocks regulate photoreceptor and retinal-circuit physiology (***Green and Besharse, 2004; Tosini et al., 2008; Cao et al., 2021***), yet the present pupil measure cannot exclude those mechanisms. The result constrains one downstream functional readout under one set of stimulus conditions; it does not localise a gate to the retina, retino-hypothalamic tract, or SCN.

### Strengths

The design combines repeated sampling, photoreceptor-directed stimulation, and temporal de-composition of the pupil response. Eleven measurement blocks were scheduled across 40 h, and the six-block inclusion criterion imposed a minimum level of temporal coverage for each retained participant-by-condition series. The hierarchical models treated repeated block-level observations as nested within participants and used participant-specific melatonin-derived periods rather than imposing a common 24-h period. These features strengthen inference about within-participant trajectories under this protocol, although they do not compensate for the small number of participants.

#### Protocol and viewing control

The 40-hour ultra-short sleep–wake protocol distributed measurement episodes across circadian phase while limiting continuous wakefulness (***Heinrichs and Spitschan, 2025b***). Measurements were conducted in a light-tight enclosure under controlled spectral conditions, reducing variation in ambient light exposure during pupillometry.

#### Maxwellian view optical system

The Maxwellian-view system, as also used by ***Zele et al. (2011***), made retinal irradiance largely independent of pupil diameter when alignment was maintained. This reduces a potential feedback confound if baseline diameter changes with time of day or alertness.

#### Silent substitution and sinusoidal stimulus design

Monochromatic stimuli co-activate photoreceptor classes and complicate pathway attribution (***Daguet et al., 2024***). Silent substitution enabled photoreceptor-directed comparisons while nominally holding non-targeted excitations constant (***Spitschan and Woelders, 2018; Woelders et al., 2018***).

The common 0.4-Hz frequency was a compromise across pathways with different temporal filtering (***Joyce et al., 2015; Spitschan et al., 2014***). Analysing both phasic tracking and tonic constriction allowed slower and faster response components to be considered without equating either component with a single receptor class.

The 5-s stimulus contained two full cycles, permitting estimation of the response at the fundamental and second harmonic. With only two cycles, however, precision depends on signal quality and the adequacy of the fitted waveform.

### Limitations

The most consequential limitation is the small effective sample. Although 12 participants completed the protocol, quality control left eight participants (seven for L–M) and 74–83 retained block-level observations per condition. Repeated observations helped characterise each participant’s trajectory, and the hierarchical models represented participant-level differences rather than treating blocks as independent participants. Population-level inference nevertheless rests on only seven or eight individuals, so small or heterogeneous circadian effects may remain unresolved. The six-block threshold ensured a minimum level of temporal coverage, but exclusions have not been summarised by protocol time; time-dependent missingness therefore cannot be ruled out. The age range of 22.9–30.5 years also limits generalisation to older and clinical populations. A larger replication should retain dense within-participant sampling while increasing the number and diversity of participants.

#### Prototype-grade pupillometry device

The RetinaWISE device was a second-generation prototype (***Gibertoni et al., 2024***). Internal smoothing obscured blink-related artefacts, sampling was irregular, and a timing desynchronisation introduced an approximately 1-s offset between recording and stimulus onset. Sampling irregularity and signal quality were partly addressed by trial-wise resampling, two-stage artefact detection, completeness thresholds for the full trial and baseline, a 0.625-s maximum-gap criterion, and averaging at least three accepted trials per retained block. These safeguards regularised the time base and reduced contamination of the cosinor estimates by gross artefacts and long gaps. They cannot recover blink information obscured by device smoothing or resolve uncertainty associated with the timing offset; device-related uncertainty was therefore reduced, not eliminated. Independent hardware timestamps and access to unsmoothed traces would strengthen a future replication.

#### Limited stimulus contrast

The colour gamut of the LEDs restricted the achievable intensity range and thereby the maximum contrast attainable in both modulation directions. The 30% Weber contrast was the greatest common modulation depth across all five stimulus directions. Using one contrast improved comparability, and every condition evoked a measurable pupil response, so the stability finding was not obtained in the absence of a response. A common nominal contrast does not, however, produce equal response amplitude or precision across pathways. Measurements at several contrasts would help distinguish contrast-dependent sensitivity from circadian modulation.

#### Strong model assumptions

Model dependence was partly addressed by using participant-specific melatonin-derived periods and participant-level effects and by reporting predictive, residual, autocorrelation, convergence, and marginal-likelihood-precision diagnostics. These checks exposed residual-normality departures and numerically imprecise Bayes factors rather than concealing them. However, the circadian model represented only an approximately sinusoidal trajectory, candidate periods were estimated from about 40 h of data using a five-parameter bi-harmonic fit, uncertainty in the selected period was not propagated, and the priors were scaled using the observed outcome dispersion. The analysis therefore operationalised circadian variation as an approximately sinusoidal trajectory over nominal protocol time with participant-specific melatonin-derived periods; observations were not aligned to an independently estimated melatonin phase marker. Non-sinusoidal or phase-localised effects could have been missed, and the constant-model comparison does not test every systematic non-constant pattern. Joint modelling of melatonin and pupil responses, prior-sensitivity analysis, repeated marginal-likelihood estimation, and alternative temporal shapes would provide stronger robustness checks.

#### Revised and omitted pupil outcomes

The revised phasic outcome was directly tied to the 0.4-Hz modulation, and the decomposition considered its fundamental and second harmonic while analysing tonic constriction separately. This provides a defined stimulus-linked measure, but it does not substitute for the registered outcome, maximum constriction normalised to broadband flux. Explicit disclosure in Materials and methods and Appendix 1 distinguishes these revised analyses from confirmatory tests; it does not restore confirmatory status. The registered outcome was not part of the main analysis; it is analysed separately in Appendix 4, where it also favours the constant-level model in every condition and does not reproduce the L–M tonic result. The post-illumination pupil response and circadian changes in baseline diameter were not analysed. The post-illumination response is also the measure most consistently associated with diurnal variation in the cited literature (***Zele et al., 2011; Münch et al., 2012; Daguet et al., 2024***). Effects expressed in these outcomes or at another temporal frequency therefore remain possible. These and other recorded outcomes could be analysed from the existing recordings if they can be reconstructed reliably, with all timing and preprocessing decisions documented and their evidential status stated explicitly.

#### Single chromatic adaptation state

All conditions were measured against the same 60 cd/m^2^ white background after a 4-min adaptation, and the Maxwellian-view system made retinal irradiance largely independent of pupil diameter when alignment was maintained. These controls improved comparability across measurement blocks, but they limit inference to one adapting spectrum. Cone-opponent effects do not always appear against a neutral background and can emerge under coloured adapting light (***Spitschan et al., 2017***). In addition, the stimulus directions were optimised for a standard observer and the planned rod-escape control was not included, so individual photoreceptor splatter and rod intrusion were not empirically excluded. Observer-specific calibration, a rod control, and multiple adapting backgrounds would address these limitations in future work (***Spitschan and Woelders, 2018; Woelders et al., 2018***).

### Implications

The consistent direction of the phasic model comparison constrains a specific hypothesis: a large, approximately sinusoidal change in 0.4-Hz pupil tracking under the tested adaptation and contrast. Support for the constant model in both the larger- and smaller-amplitude conditions argues against interpreting the result solely as failure to evoke a measurable response. It does not establish that retinal sensitivity is invariant, explain earlier effects measured with other outcomes, or show that time of day is irrelevant to pupillometry.

The contrast between stable phasic tracking and the possible L–M tonic signal raises the more specific possibility that temporal variation depends on the response component rather than acting as a uniform change in pupil responsiveness. Because the L–M evidence was numerically imprecise and based on seven participants, this is a hypothesis for replication rather than a demonstrated dissociation.

Chromatic pupillometry is increasingly used to assess retinal and subcortical function, including in neurodegenerative disease (***Steiner and de Zeeuw, 2024***). Whether clinical measurements require time-of-day standardisation should be tested directly in larger and clinically representative samples using the outcomes of interest.

### Conclusion

Across all five photoreceptor-directed conditions, the constant-level model was favoured for phasic pupil tracking at 0.4 Hz. The agreement across stimulus directions, under controlled adaptation and a protocol designed to limit progressive sleep pressure, supports stable phasic tracking across the sampled protocol rather than a general circadian pattern in this response component. This finding defines a boundary condition for circadian effects on pupillometry; it does not establish the absence of circadian retinal regulation. Smaller, non-sinusoidal, tonic, post-stimulus, contrast-dependent, or adaptation-dependent effects remain possible. The imprecise L–M tonic result provides a focused hypothesis for preregistered replication in a larger sample using the protocol-specified pupil outcomes.

## Materials and methods

This experimental study investigated circadian modulation of photoreceptor-directed pupil responses. The protocol is described in detail in ***Heinrichs and Spitschan (2025b***). Data were collected between January and October 2024. The study was approved by the Ethics Committee of the Technical University of Munich (2023-369-S-SB). All participants provided written informed consent. All research was conducted in accordance with the Declaration of Helsinki.

### Participant sample

Twelve participants enrolled and completed the study. Four did not meet the data-availability criterion for the circadian analysis. The final sample therefore comprised eight participants (three reported as female), aged 22.9–30.5 years. Participant characteristics, including sleep parameters collected during recruitment, are listed in Table 7.

**Table 7.**
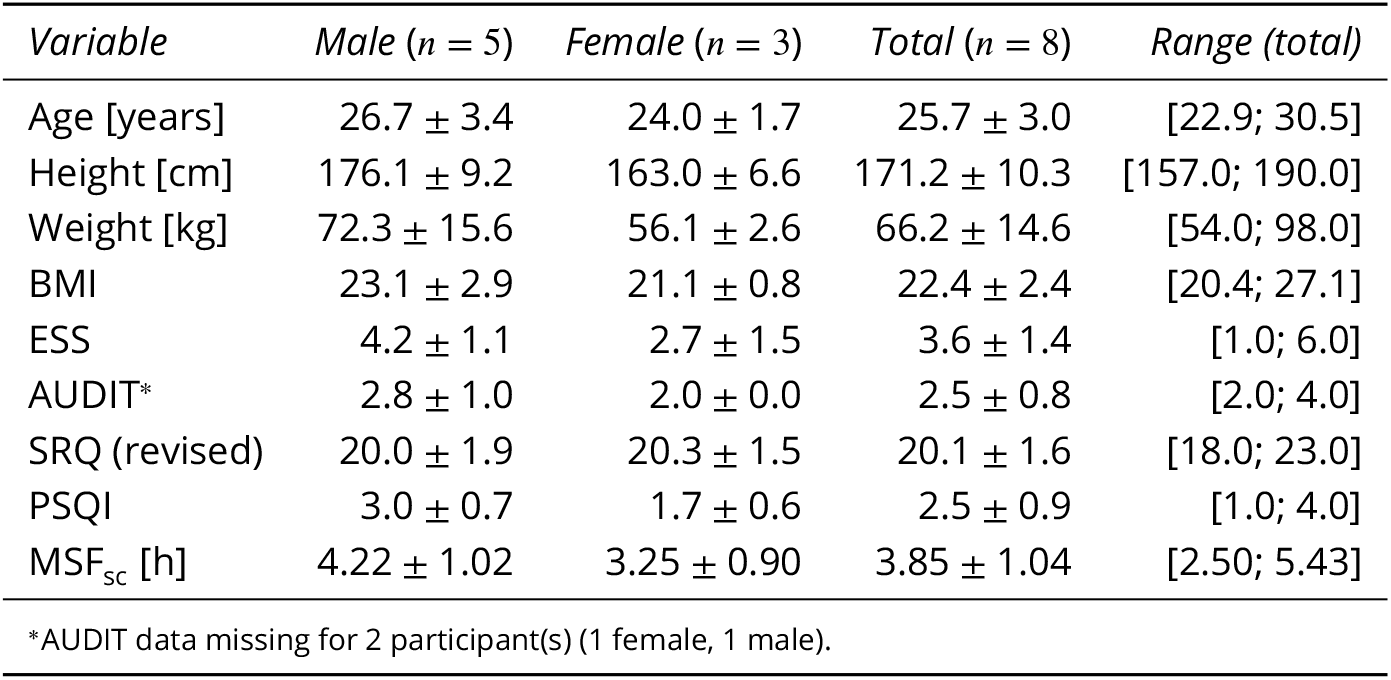
Demographic and questionnaire summary by sex. Values are mean ± SD; the final column gives the total-sample range as [min; max]. Age is given in decimal years. ESS = Epworth Sleepiness Scale; AUDIT = Alcohol Use Disorders Identification Test; SRQ = Sleep Regularity Questionnaire (revised); PSQI = Pittsburgh Sleep Quality Index; MSF_sc_ = sleep-corrected mid-sleep on free days (chronotype).

| *Variable* | *Male (n = 5)* | *Female (n = 3)* | *Total (n = 8)* | *Range (total)* |
| --- | --- | --- | --- | --- |
| Age [years] | 26.7 ± 3.4 | 24.0 ± 1.7 | 25.7 ± 3.0 | [22.9; 30.5] |
| Height [cm] | 176.1 ± 9.2 | 163.0 ± 6.6 | 171.2 ± 10.3 | [157.0; 190.0] |
| Weight [kg] | 72.3 ± 15.6 | 56.1 ± 2.6 | 66.2 ± 14.6 | [54.0; 98.0] |
| BMI | 23.1 ± 2.9 | 21.1 ± 0.8 | 22.4 ± 2.4 | [20.4; 27.1] |
| ESS | 4.2 ± 1.1 | 2.7 ± 1.5 | 3.6 ± 1.4 | [1.0; 6.0] |
| AUDIT* | 2.8 ± 1.0 | 2.0 ± 0.0 | 2.5 ± 0.8 | [2.0; 4.0] |
| SRQ (revised) | 20.0 ± 1.9 | 20.3 ± 1.5 | 20.1 ± 1.6 | [18.0; 23.0] |
| PSQI | 3.0 ± 0.7 | 1.7 ± 0.6 | 2.5 ± 0.9 | [1.0; 4.0] |
| $\text{MSF}_{\text{sc}}$ [h] | 4.22 ± 1.02 | 3.25 ± 0.90 | 3.85 ± 1.04 | [2.50; 5.43] |

\*AUDIT data missing for 2 participant(s) (1 female, 1 male).

### Study protocol

The study protocol consisted of two in-lab visits, extending across a total of three weeks. Both visits were preceded by a 1-week stabilisation period, in which participants adhered to a stable sleep-wake rhythm and stable meal times; the second visit was additionally followed by one more week of circadian stabilisation. The first in-lab visit consisted of an evening session and an adaptation night. Over the course of the evening, participants were introduced to and familiarised with the devices and experimental procedures, in order to facilitate timely completion during the data collection. Participants were here exposed to the light output of the experimental devices during the first evening session (“experimental light exposure evening” session, LE).

During the second in-lab visit, participants completed an evening in dim light, an adaptation night, and, after waking the following morning, a 40-hour ultra-short sleep–wake schedule designed to distribute sleep and wake across circadian phase. The evening session served to homogenise light history and did not involve visual stimuli (“dim-light evening” session, DL). The 40-hour session is hereafter referred to as the “forced desynchrony (FD)” session. It provided 11 scheduled wake/measurement blocks. The nominal schedule alternated 2 h 30 min of wakefulness with 1 h 15 min sleep opportunities (sleep–wake ratio 1:2; nominal cycle length 3 h 45 min).

Pupillometry data were collected on two occasions during each wake period to reduce participant burden from darkness and passive viewing.

#### Saliva collection

Saliva samples (>1 mL) were collected approximately every 45 minutes throughout the forced-desynchrony session, except during sleep periods, using Salivette collection devices (Sarstedt, Nürnberg, Germany). Participants chewed on a cotton swab for 5 minutes before placing it into the collection tube. Samples were centrifuged for 3 minutes at 3,000 rpm and stored at −20°C until analysis. Melatonin concentrations were determined by radioimmunoassay (RIA, RK-DSM2, NovoLytiX GmbH, Switzerland) with a quantification range of 0.5–50 pg/mL, a detection limit of 0.2 pg/mL, mean intra-assay precision of 7.9%, and mean inter-assay precision of 9.8%. Concentrations were analysed as reported by the laboratory: values below the detection limit or the lower limit of quantification, including values reported as 0 pg/mL, were neither substituted nor excluded. Salivary melatonin concentrations were used to estimate each participant’s circadian period as described under “Salivary melatonin fitting”.

#### Experimental setup

Pupillary light responses were measured using the second prototype of the RetinaWISE device (***Gib-ertoni et al., 2024***), a custom-built six-primary Maxwellian-view stimulation system that enables precise temporal control of the spectral composition of retinal stimulation. The device delivers spatially homogeneous retinal illumination, yet permits the presentation of custom stimulus geometries through insertion of optical field stops. Pupil tracking was performed by built-in infrared cameras that alternated between the two eyes. The experimental setup was placed within a commercially available light-tight enclosure, a tent originally designed for photographic darkroom applications, to minimise contamination by ambient light and ensure that participants were exposed only to the experimental light stimuli. During measurements, the participant was seated in front of the stimulation device, resting their head in front of the device on a chin rest and against a head rest. The experimenter performed the optical alignment procedure through a small opening at the back of the tent. The optical system was configured in Maxwellian view, allowing retinal irradiance to remain largely independent of changes in pupil diameter when the participant was correctly aligned with the optical axis. To minimise head movements and maintain alignment throughout the recordings, the participant’s head was fixed with an elastic headband attached to the head rest. Stimuli were presented monocularly to the dominant eye as an annulus with an inner diameter of 10° and an outer diameter of 30° of visual angle (5–15° eccentricity). Monocular stimulation was chosen because the interaction between circadian phase and binocular facilitation of the pupillary light response remains insufficiently characterised. Eye dominance was determined with the Miles test (***Miles, 1930***). Only the trace from the stimulated dominant eye was analysed; the non-dominant-eye trace was not used. Pupil diameter was quantified in millimetres using the proprietary pupil-tracking algorithm implemented in the software provided with the RetinaWISE device.

#### Experimental procedure

Each test began with a 4 min adaptation phase in which participants were exposed to the 60 cd/m^2^ background stimulus at levels typically considered to be rod-saturating (***Aguilar and Stiles, 1954; Adelson, 1982; Spitschan et al., 2017***). It was followed by 25 trials of 18 s each. Within each trial, participants were exposed to sinusoidal light modulations at 0.4 Hz. This temporal frequency was selected as a practical compromise at which each targeted pathway was expected to produce a measurable pupil response. Each stimulus epoch lasted 5 s and was followed by 13 s of interleaved white-light baseline stimulation (60 cd/m^2^), resulting in a total trial duration of 18 s. There were five stimulus conditions, targeting different photoreceptors and post-receptoral mechanisms: broadband light flux modulation of all cone classes and melanopsin (FLUX), L–M, achromatic luminance modulation, equally for all cones (LMS), MEL, and chromatic S. Each block was scheduled to contain two experimental runs, each beginning with one adaptation phase and containing five trials per condition. The order of conditions within each run of 25 trials (five per condition) was derived from an m-sequence to approximately counterbalance first-order carry-over effects between conditions (***Buračas and Boynton, 2002; Brooks, 2012***). The 24 transitions between successive trials covered most, but not all, of the 25 ordered pairs of conditions. To increase the number of trials available, each wake period was scheduled to include two experimental runs, separated by a short break. If a run had to be repeated, for example because the signal was lost partway through, the valid trials recorded before the interruption were retained. All accepted trials recorded within a participant’s wake period, or “experimental block”, were averaged and analysed together, irrespective of the run in which they were recorded, to achieve a higher trial count for the block averages. In 4 of the 83 analysed participant blocks, more trials were accepted than two runs contain, so the number of accepted trials per condition and block ranged from 3 to 16 (median 9).

#### Silent substitution stimuli

The five stimulus conditions were each designed to up- and down-modulate a photoreceptor class or defined combination while nominally holding the remaining photoreceptor excitations constant. These comprised a melanopsin-directed condition (MEL); conditions probing post-receptoral mechanisms underlying luminance (LMS) and chromatic vision (L–M and S); and a broadband condition that modulated all cone classes and melanopsin together at equal contrast. All stimuli were presented as sinusoidal modulations at 0.4 Hz, the common frequency selected to elicit measurable responses across the targeted pathways while allowing repeated presentation of every condition. Photoreceptor activation was modulated around the baseline level, with contrast peaks at 0.625 and 3.125 s and contrast troughs at 1.875 and 4.375 s relative to stimulus onset. A peak Weber contrast of 30% was chosen as the target modulation depth for each targeted photoreceptor class, representing the maximum contrast achievable across all five conditions. LMS modulated L, M and S cones in phase; FLUX modulated L, M and S cones and melanopsin in phase; L–M modulated L and M cones in counterphase, each at 30%; S and MEL modulated only the targeted class. The remaining classes were nominally held at 0% contrast. For each condition, we used a numerical optimisation procedure (PySilSub; ***Martin et al., 2023***) to determine the spectral composition that achieved this target contrast. The optimisation was based on the default action spectra of the 32-year-old standard observer for the cone fundamentals and melanopsin (***Commission Interna-tionale de l’Eclairage, 2018***); the resulting directions are therefore photoreceptor-directed rather than a demonstration of perfect isolation in every participant. Rods were not included in the optimisation, so the directions were not designed to silence rods; rod intrusion was not tested, and rod signals were assumed to be saturated by the 60 cd/m^2^ background. The background (baseline) spectrum was defined by setting each LED channel to half its maximum output. This provided sufficient dynamic range in both directions to accommodate the positive and negative excursions of the sinusoidal modulation across all five conditions within a single experimental run.

### Data preprocessing

Pupillometry data were preprocessed in a series of steps including resampling, artefact removal, and trial- and session-level quality control, using custom routines written in Python. As part of data preparation, parameters were obtained from the block-averaged pupil traces with a two-step parameter extraction pipeline.

#### Raw signal preprocessing and quality control

##### Device-level processing and timing

The pupillometry device performed internal preprocessing, including smoothing the raw pupil size signal and rudimentary blink detection. However, the internal smoothing algorithm obscured the characteristic rapid signal drops (100–400 ms) typically associated with blinks (***Milo et al., 2009***, BNID 100706), precluding reliable blink detection from the smoothed signal. We therefore implemented a custom artefact detection pipeline, described below.

A timing desynchronisation between device components resulted in an approximately 1 s delay in light stimulus output relative to the onset of recording, along with an unspecified delay during file saving at the end of each sequence. Consequently, the recorded sequences were longer than the 18 s specified in the protocol.

##### Resampling

Because the device did not operate at a fixed sampling rate, we first removed all samples with missing pupil size values, then computed the mean inter-sample interval per trial and derived a realised sampling frequency as 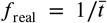. Data were resampled trial-wise to this frequency to regularise the time base while preserving the trial structure.

##### Artefact removal

Samples with pupil diameter values outside the physiological range of 1.5–9 mm were excluded (***Kret and Sjak-Shie, 2019***). Artefact detection then proceeded in two stages. First, constriction and dilation velocity were computed and subjected to outlier detection based on a rolling median absolute deviation (MAD) with a window size of 2 s (approximately 10 % of trial length) and a scaling factor of *n* = 1.5. Second, a size-based artefact detection step targeted residual spikes in the pupil size signal using the same rolling MAD approach with a more conservative scaling factor of *n* = 4.5. Both scaling factors were determined by visual inspection of representative traces. This approach relies on statistical outlier detection rather than explicit gap detection, given the limitations of the smoothed signal noted above.

##### Data completeness and trial rejection

Trials were evaluated for data completeness to avoid distortions in the trial-averaged pupil trace. Because the response was baseline-corrected, completeness was considered both across the trial and within the baseline. We inspected joint trial-acceptance matrices over candidate thresholds from 0 % to 100 %. A trial was retained if at least 75 % of the full trial and at least 40 % of the baseline interval ([−1, 0] s relative to stimulus onset) were complete. Trials failing either threshold were rejected. A secondary trial-rejection criterion addressed spectral integrity. Given the sinusoidal stimulus and response at 0.4 Hz, the maximum permitted gap was 0.625 s, one quarter of a 0.4-Hz cycle, to limit aliasing at the stimulus frequency and its second harmonic.

##### Block and participant rejection per condition

At the block level, a minimum of three accepted trials per condition and block was required; accepted trials were then averaged to yield a single pupil trace per participant, condition, and block. At the participant level, a participant was included in the circadian analysis of a given condition only if averaged traces for at least six blocks were available, ensuring sufficient temporal coverage for cosinor fitting of a near-24 h rhythm. Data quality was thus enforced at two levels, the block level and the participant level.

#### Two-step parameter extraction

The baseline-corrected (i.e. relative) trial-averaged pupil traces were processed per participant, chromatic condition, and block using a two-step decomposition pipeline separating tonic (sustained) from oscillatory (phasic) components.

The raw pupil response to each modulation conflates these two distinct temporal dynamics. Separating these components is informative beyond the raw response because photoreceptor pathways also differ in their temporal filtering properties (***Spitschan et al., 2014***). The sluggish, sustained signalling of ipRGCs is expected to weight the tonic component, and faster cone-driven contributions the phasic tracking of the 0.4 Hz modulation (***Barrionuevo et al., 2014***), adding a temporal axis of selectivity that complements the spectral isolation of silent substitution. This biases the two components toward, rather than cleanly separating, the underlying channels: melanopsin also contributes to phasic responses at these frequencies (***Barrionuevo et al., 2014***), so a circadian modulation of the phasic amplitude in the MEL condition would still be interpretable as an ipRGC effect.

##### Step 1: Tonic component extraction via low-pass filtering

To isolate the slow tonic component, a low-pass filter was applied to the trial-averaged pupil trace of each participant, condition and block. The filter was a zero-phase second-order Butterworth, implemented via forward-backward filtering with a cutoff of 0.2 Hz, yielding an effective fourth-order response. The filter was applied to the full averaged trace (−1 to 18 s relative to stimulus onset). Missing samples were linearly interpolated for filtering only and set back to missing in the tonic component and the residual, so they did not enter the tonic features or the cosinor fit. Three summary features were extracted from the tonic signal component within the 5-second stimulus-presentation phase, i.e. the [0, 5] s window from stimulus onset to offset, hereafter referred to as the stimulus period-of-interest (POI). The principal tonic feature was the tonic minimum, defined as the most negative deflection of the low-pass-filtered signal during the stimulus POI, reflecting the maximum depth of sustained constriction relative to baseline. Two further features characterised the tonic response: the latency of that minimum, measured in seconds from stimulus onset, and the tonic mean level, defined as the arithmetic mean of the filtered signal across the stimulus window. The residual was obtained by subtracting the tonic signal from the original trace, representing the fast, transient component which was processed downstream.

##### Step 2: Phasic component extraction via bi-harmonic cosinor fitting

The oscillatory residual was modelled within the same [0, 5] s POI window (stimulus onset to offset, with no post-stimulus extension) as a bi-harmonic cosinor. Throughout this work, cosinor models are written in the amplitude–phase form

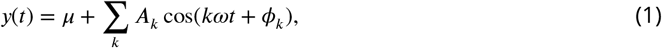

so that ϕ_*k*_ is the phase angle and the peak of harmonic *k* falls at *t* = −ϕ_*k*_/(*k*ω). The same convention is used for the pupil trace, salivary melatonin, and circadian (Eq. 5) models alike. For the bi-harmonic pupil and saliva models, the equivalent linear sine–cosine parameterisation was fitted by ordinary least squares:

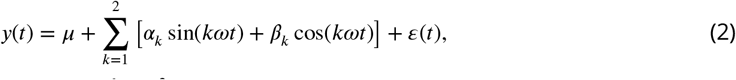

where μ is the mean level and ε(*t*) ∼ *N* (0, σ^2^) is the Gaussian residual error, assumed for the pupil, saliva, and circadian models alike. Eq. 2 describes a single fit and therefore carries no participant or block indices; the pupil and saliva models were each fitted independently to every unit of data, as specified below. The hierarchical circadian model introduced later indexes participants and blocks explicitly. For the pupil model, *y*(*t*) is the phasic pupil residual from Step 1 and *t* is time in seconds from stimulus onset, with *ω* = 2*πf* and *f* = 0.4 Hz the chromatic stimulus modulation frequency. Because the tonic component has already been removed, μ here is a residual offset close to zero rather than a mesor of the pupil signal. Subscripts denote the harmonic index: *k* = 1 for the fundamental and *k* = 2 for the second harmonic. Amplitude and phase were derived for each harmonic *k* as:

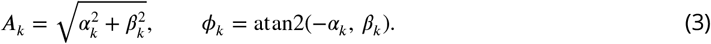

The two fitted harmonics are written *A*_*f*_, *ϕ*_*f*_ for the fundamental (*k* = 1) and *A*_2*f*_, *ϕ*_2*f*_ for the second harmonic (*k* = 2), with slopes α_*f*_, β_*f*_ and α_2*f*_, β_2*f*_ correspondingly. Naming the subscript after the stimulus frequency keeps these quantities distinct from the circadian amplitude *A*_circ_ (Eq. 6).

The main outcome reported here was the fundamental amplitude *A*_*f*_ (in mm), reflecting the magnitude of pupil oscillation at the 0.4-Hz stimulus frequency. The tonic minimum from Step 1 provided a complementary outcome. Additional extracted parameters were the second-harmonic amplitude *A*_2*f*_ at 0.8 Hz, the fundamental and second-harmonic phases *ϕ*_*f*_ and *ϕ*_2*f*_ (in radians), and the pre-stimulus baseline mean (the arithmetic mean of absolute pupil size across the pre-stimulus phase of each trial).

##### Aggregation and baseline correction

For each participant, condition and experimental block, pupil traces were averaged sample-wise across at least three accepted trials, ignoring missing samples. The averaged trace was then baseline-corrected to its own pre-stimulus mean over 1 second, and the resulting relative pupil size was used for further analysis. Figure 3 shows the relative pupil response across phases for all five conditions (FLUX, L–M, LMS, MEL, and S): the 1-second pre-stimulus phase, the stimulus presentation (grey shading), and the post-stimulus phase. For visualisation, traces in this figure were further averaged across blocks and participants.

Averaging required at least three valid trials to be available; however, small gaps of missing data could occur within a trial, so an individual point on the average trace might be the mean of three, two, or even a single trial. No further smoothing was applied before the decomposition described below.

### Statistical analysis

The statistical analysis proceeded in two stages: salivary melatonin concentrations were first modelled to obtain a participant-specific estimate of the circadian period, which then set the period of the pupil model. Two parameters from the decomposition were carried forward: the cosinor amplitude at the fundamental stimulus frequency, quantifying phasic tracking of the chromatic modulation, and the tonic response minimum. For each, we compared a specified approximately circadian sinusoidal model with a constant-level model.

#### Deviations from the published protocol

The analyses reported here differ from the published protocol (***Heinrichs and Spitschan, 2025b***) in the following respects:

- **Pupil outcomes**. The protocol specified maximum constriction during the 5-s stimulus, normalised to the broadband flux response, as the primary pupil outcome. The present report instead analyses the unnormalised 0.4-Hz phasic amplitude as its main outcome and tonic minimum as a complementary outcome.
- **Melatonin model**. The protocol specified a single-harmonic cosinor; the present report uses a bi-harmonic cosinor.
- **Candidate circadian periods**. The protocol specified a search from 20 to 28 h; the present analysis searches from 22 to 28 h.
- **Sequential stopping**. The protocol specified a sequential Bayes-factor stopping rule based on the melanopsin-directed response, with a maximum sample of 12 participants. Twelve participants completed the protocol, of whom eight contributed to the analyses after quality control. The sequential rule was not applied, because evaluating the evidence during data collection was not feasible alongside the demands of running the protocol; the sample was therefore fixed rather than sequentially stopped.
- **Rod-escape control**. The protocol planned a subsequent rod-escape control; that control is not included in the present report.

Accordingly, the phasic-amplitude and tonic-minimum analyses are described as revised analyses rather than confirmatory tests of the protocol’s primary pupil outcome. Appendix 1—table 1 provides the same comparison in tabular form and states the implications for interpretation. Where the timing or rationale for a change was not documented in the materials available for preparing this manuscript, none is inferred here.

#### Salivary melatonin fitting

The assumed circadian period for each participant was obtained by analysing salivary melatonin concentration. For each participant, salivary melatonin concentration *h*(*t*) was modelled using the bi-harmonic cosinor in Eq. 2, with *y*(*t*) = *h*(*t*) and *ω* = 2π/τ. Here, *t* is time in hours since protocol start, τ is the participant-specific melatonin-derived fixed period within a given fit, μ is the melatonin mesor, and ε(*t*) ∼ *N* (0, σ^2^) is the Gaussian residual error of Eq. 2. The model was fitted via ordinary least squares independently for each participant and each candidate period τ ∈ [22; 28] h in iterative steps of 1/60 h. The best-fitting period 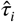 was selected per participant by minimising the Akaike information criterion (AIC). Amplitude and phase for each harmonic *k* were derived according to Eq. 3; the phase was expressed in radians.

#### Circadian mixed-effects model specification

The phasic analysis combined two per-participant quantities: the circadian period estimate (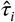, in hours) from the saliva cosinor model, which set the period duration, and the pupil cosinor amplitude at the stimulus fundamental frequency, which was the outcome. The same model comparison was then applied to tonic minimum.

##### Outcome selection

Prior to model fitting, each outcome distribution was inspected by condition using histograms and normal quantile–quantile plots. The Shapiro–Wilk statistic (***Shapiro and Wilk, 1965***), skewness, and excess kurtosis were computed as additional descriptors. We retained the original scale and used the Gaussian likelihood specified below (Eq. 5); departures from residual normality in selected conditions temper interpretation and are reported with the model diagnostics.

##### Model specification

For each of the five stimulus conditions, two hierarchical Bayesian models were specified. Throughout this section, *i* indexes participants and *b* experimental blocks. The null model *M*_0_ assumed a constant pupil amplitude across the participant’s circadian rhythm, with participant-specific mean levels (intercepts):

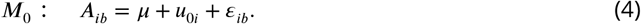

The circadian cosinor model *M*_1_ additionally included fixed and random sine and cosine slopes, assuming a circadian modulation of the outcome with a participant-specific period 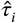 and participant-specific amplitude and phase:

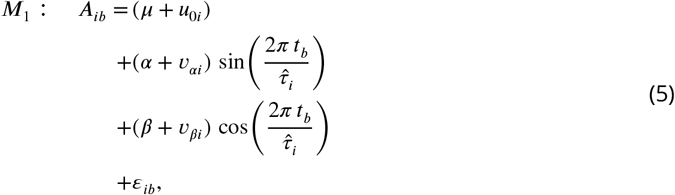

where *A*_*ib*_ is the outcome for participant *i* in block *b*, taken as the phasic fundamental amplitude *A*_*f*_ or the tonic minimum; *t*_*b*_ = *b*×3.75 is the nominal elapsed time in hours since protocol start; 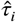 is the individual period estimate from the saliva application of the bi-harmonic cosinor model (Eq. 2), giving the participant-specific angular frequency 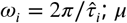 is the population intercept; *α* and *β* are the population-level sine and cosine slopes; *u*_0*i*_, *v*_*αi*_, *v*_*βi*_ are participant-specific random deviations; and *ε*_*ib*_ is the residual error. Random effects were modelled as independent Gaussian distributions, i.e. the random-effect covariance matrix was constrained to be diagonal, assuming no correlation between individual differences in intercept, sine slope, and cosine slope. This choice was motivated by the small sample size, which would not support reliable estimation of off-diagonal covariance terms. Unlike the bi-harmonic pupil and salivary melatonin models, the circadian model includes only the fundamental harmonic and is fitted hierarchically, with population-level effects and participant-specific random effects. The two circadian regressors are denoted 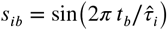 and 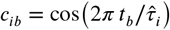_*i*_, on which α and β are the respective slopes. Models were compiled with `bambi:` *M*_0_ : *A* ∼ 1 + (1 ∣ participant) and *M*_1_ : *A* ∼ 1 + *s* + *c* + (1 + *s* + *c* ∣ participant).

##### Priors

Priors were weakly informative and scaled by the standard deviation of the outcome within each condition, SD(*y*). The intercept had a Normal prior centred at the empirical mean with SD = 1.5 × SD(*y*), and the sine and cosine slopes α and β had Normal priors centred at zero with SD = 0.7×SD(*y*). The standard deviations of the participant-specific deviations had Half-Student-*t* priors (ν = 4) with scale SD(*y*) for the intercept and 0.7 × SD(*y*) for the sine and cosine slopes. The residual standard deviation had a Half-Student-*t* prior (ν = 4) with scale SD(*y*), the `bambi` default. Models were built with `bambi` 0.17.2 and PyMC 5.28.1. Because these scales use the observed outcome dispersion, the Bayes factors are conditional on this data-dependent prior specification and should be interpreted accordingly.

##### Inference and model comparison

Both models were fitted using Sequential Monte Carlo (SMC) with 10000 draws, which yields posterior samples and an estimate of the log marginal likelihood log *Z* required for Bayes factor computation. Model evidence for the circadian model compared with the constant model was quantified as the Bayes factor: 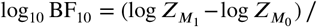; positive values favour the circadian model M1, and negative values favour the constant-level model M0. Monte Carlo precision was assessed using the ratio of the SMC log-marginal-likelihood standard deviation *SD*_*SMC*_ to the absolute difference between M0 and M1 (SD/ |Δ log *Z*|). We used 0.10 as a pragmatic internal threshold; values above it indicate that the numerical Bayes-factor estimate is imprecise, not that the corresponding biological effect is absent. Convergence was assessed via R-hat 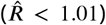 for all parameters of both M0 and M1. Model fit was summarised using Bayesian *R*^2^ (***Gelman et al., 2019***), computed separately for conditional (fixed + random effects) and marginal (fixed effects only) predictions.

##### Circadian parameter extraction

For each condition the following quantities were derived from the posterior distributions: fixed-effect estimates for the intercept μ and the sine and cosine slopes α and β, and random-effect standard deviations for all three terms to quantify between-participant variability (posterior mean and 95 % highest-density interval, HDI). Bayesian *R*^2^ is summarised as the posterior median and a 95 % equal-tailed interval (ETI; 2.5th and 97.5th percentiles) because *R*^2^ is bounded on [0, 1]. Since HDI and ETI are both 95 % credible intervals, either is referred to as the 95 % CI; the fixed-effect posteriors were unimodal and approximately symmetric on visual inspection of the model diagnostics, so the two interval types nearly coincide for the reported estimates. Finally, the population-level circadian amplitude (in mm) and phase (in radians) were derived from the fixed-effect posterior samples, following the same amplitude–phase convention as in the pupil and saliva cosinor models (Eq. 3):

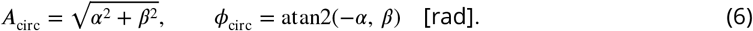

## Data availability

De-identified raw and processed pupillometry and melatonin data, inclusion masks, analysis and figure code, and environment metadata will be deposited in Edmond before submission. Persistent identifiers, licences, and an archived code release will be added here when available.

## Use of generative AI

OpenAI Codex was used for language editing, structural editing, and manuscript-format checks. It was not used to generate or analyse study data. The authors reviewed the suggested changes and remain responsible for the manuscript’s content.

## Author contributions

**H.S.H**.: Conceptualization, Data Curation, Formal Analysis, Investigation, Methodology, Project Administration, Software, Validation, Visualization, Writing – Original Draft Preparation, Writing – Review & Editing. **D.G**.: Data Curation, Formal Analysis (Preprocessing). **M.S**.: Conceptualization, Funding Acquisition, Methodology, Project Administration, Resources, Supervision, Writing – Review & Editing.

## Acknowledgements

We thank Jannik Romanowski and Johannes Boldt for adapting the experimental setup and enclosure tent and for refining the chin rest; Martial Geiser for support with optical engineering and calibration; and Katharina Stein and Constanza Bravo for piloting the protocol and informing refinement of the procedure.

## Appendix 1

### Analysis scope and deviations from the published protocol

The published protocol for this study (***Heinrichs and Spitschan***, ***2025b***) and the analyses reported in the article differ in several material respects. Appendix 1—table 1 records those differences so that the evidential status of the present results is explicit. It does not assign a rationale or timing to changes that are not documented in the materials accompanying this manuscript.

#### Rationale for the revised pupil outcome

The following rationale is provided to aid interpretation. It was not documented in the protocol record and does not restore confirmatory status.

The stimulus was a continuous 0.4-Hz sinusoid, designed so that the pupil would track the modulation throughout the stimulus epoch. The expected response is therefore a sustained oscillation, which is more appropriately quantified by a cosinor fit and its amplitude at the stimulus frequency than by maximum constriction. Maximum constriction captures only the single deepest excursion and is better suited to a brief light pulse that evokes a transient response (***McAdams et al., 2018***). The revised decomposition also separates the slow, sustained constriction (tonic) from the oscillation at the stimulus frequency superimposed on it (phasic), which need not vary in the same way across circadian phase.

This reasoning is borne out in the data. Appendix 4 derives the registered outcome and shows that it is dominated by the sustained component rather than by oscillatory tracking: the L–M condition yields the largest maximum constriction of any condition while having one of the smallest phasic amplitudes, so the two outcomes order the conditions differently. Maximum constriction is therefore not a less precise measure of the same quantity; it measures something else.

The protocol also specified dividing each condition-specific response by the broadband FLUX response, with the aim of removing circadian variation common to the photoreceptor pathways and isolating variation specific to the targeted photoreceptor. This is a sound approach in principle, but its interpretation here has two limitations. First, the broadband FLUX response is itself one of the five conditions whose variation across circadian phase is of interest, so that variation must be characterised rather than assumed away through its use as a denominator. Second, a flat ratio may indicate either that neither response varies or that both vary together, and distinguishing these cases is precisely what a circadian test is intended to do. If circadian variation is observed in both the FLUX condition and another condition, their relative responses may then be explored in a secondary analysis. That ratio analysis is reported in Appendix 4; it favours the constant-level model in every condition, as the phasic amplitude does. The tonic minimum agrees in four of five conditions; in L–M it favours the circadian model, with a numerically imprecise Bayes factor (see the main article).

### Scope of the Bayesian comparison

For each stimulus condition, M0 contains a population intercept and participant-specific intercepts. M1 adds population and participant-specific sine and cosine terms at a period set by each participant’s melatonin fit. The Bayes factor therefore compares these two complete hierarchical specifications; it is not a model-free test of whether any temporal variation exists. The priors are scaled using the observed outcome dispersion, so the Bayes factors are also conditional on this data-dependent choice.

Time is represented by nominal block time (*t*_*b*_ = *b* × 3.75 h), and the melatonin estimate sets the period but does not align observations to an independently estimated melatonin phase marker. Uncertainty in the estimated period is not propagated into the pupil model. Consequently, the analysis is most directly interpreted as a comparison between constant and approximately sinusoidal trajectories across nominal protocol time.

The derived circadian amplitude 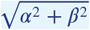 is non-negative by construction, and its phase is weakly identified when both slopes are near zero. Sine and cosine slopes, the complete model comparison, and their numerical precision therefore take precedence over derived amplitude and phase summaries.

## Appendix 2

### Complete model output for phasic pupil tracking

This appendix reports the full posterior parameter estimates and numerical diagnostic summary for the revised phasic-amplitude outcome analysed in the article. It adds no outcome, model comparison, or hypothesis test beyond those reported in the main text.

**Appendix 2—table 1.** Parameter estimates for circadian cosinor models (Amplitude (fundamental)) across conditions. Values are posterior means with 95% credible intervals, except *R*^2^, reported as posterior median with a 95% equal-tailed interval. M1 = cosinor model; M0 = null (intercept-only) model.

| Condition | Model parameter | M1 (cosinor) |  | M0 (null) |  |
| --- | --- | --- | --- | --- | --- |
|  |  | Estimate | 95% CI | Estimate | 95% CI |
| FLUX | *Fixed effects* |  |  |  |  |
|  | Intercept | **0.255** | **[0.188; 0.319]** | **0.255** | **[0.182; 0.326]** |
| | $\alpha$ (sin) | 0.013 | [-0.010; 0.036] | | |
| | $\beta$ (cos) | -0.010 | [-0.034; 0.015] | | |
|  | *Random effects* |  |  |  |  |
|  | SD(Intercept) | **0.104** | **[0.070; 0.141]** | **0.114** | **[0.073; 0.158]** |
| | SD($\alpha$) | **0.016** | **[0.000; 0.038]** | | |
| | SD($\beta$) | **0.023** | **[0.001; 0.047]** | | |
|  | *Model fit* |  |  |  |  |
| | Conditional $R^2$ | **0.786** | **[0.700; 0.845]** | **0.769** | **[0.680; 0.830]** |
| | Marginal $R^2$ | **0.053** | **[0.003; 0.196]** | | |
| L-M | *Fixed effects* |  |  |  |  |
|  | Intercept | **0.117** | **[0.089; 0.145]** | **0.117** | **[0.087; 0.146]** |
| | $\alpha$ (sin) | 0.001 | [-0.017; 0.019] | | |
| | $\beta$ (cos) | -0.014 | [-0.029; 0.002] | | |
|  | *Random effects* |  |  |  |  |
|  | SD(Intercept) | **0.038** | **[0.019; 0.059]** | **0.039** | **[0.019; 0.062]** |
| | SD($\alpha$) | **0.011** | **[0.000; 0.027]** | | |
| | SD($\beta$) | **0.009** | **[0.000; 0.022]** | | |
|  | *Model fit* |  |  |  |  |
| | Conditional $R^2$ | **0.413** | **[0.241; 0.560]** | **0.363** | **[0.185; 0.519]** |
| | Marginal $R^2$ | **0.070** | **[0.004; 0.218]** | | |
| LMS | *Fixed effects* |  |  |  |  |
|  | Intercept | **0.223** | **[0.164; 0.282]** | **0.224** | **[0.158; 0.287]** |
| | $\alpha$ (sin) | -0.001 | [-0.023; 0.021] | | |
| | $\beta$ (cos) | **-0.031** | **[-0.058; -0.005]** | | |
|  | *Random effects* |  |  |  |  |
|  | SD(Intercept) | **0.092** | **[0.060; 0.126]** | **0.101** | **[0.062; 0.146]** |
| | SD($\alpha$) | **0.013** | **[0.000; 0.032]** | | |
| | SD($\beta$) | **0.027** | **[0.001; 0.052]** | | |
|  | *Model fit* |  |  |  |  |
| | Conditional $R^2$ | **0.753** | **[0.650; 0.822]** | **0.703** | **[0.591; 0.781]** |
| | Marginal $R^2$ | **0.143** | **[0.014; 0.367]** | | |
| MEL | *Fixed effects* |  |  |  |  |
|  | Intercept | **0.095** | **[0.076; 0.114]** | **0.095** | **[0.074; 0.115]** |
| | $\alpha$ (sin) | -0.004 | [-0.019; 0.011] | | |
| | $\beta$ (cos) | -0.005 | [-0.017; 0.007] | | |
|  | *Random effects* |  |  |  |  |
|  | SD(Intercept) | **0.027** | **[0.013; 0.042]** | **0.028** | **[0.013; 0.045]** |
| | SD($\alpha$) | **0.013** | **[0.000; 0.028]** | | |
| | SD($\beta$) | **0.007** | **[0.000; 0.017]** | | |
|  | *Model fit* |  |  |  |  |
| | Conditional $R^2$ | **0.344** | **[0.177; 0.503]** | **0.295** | **[0.126; 0.454]** |
| | Marginal $R^2$ | **0.037** | **[0.001; 0.157]** | | |
| S | *Fixed effects* |  |  |  |  |
|  | Intercept | **0.190** | **[0.145; 0.236]** | **0.189** | **[0.140; 0.240]** |
| | $\alpha$ (sin) | -0.015 | [-0.032; 0.002] | | |
| | $\beta$ (cos) | **-0.020** | **[-0.035; -0.004]** | | |
|  | *Random effects* |  |  |  |  |
|  | SD(Intercept) | **0.072** | **[0.047; 0.097]** | **0.078** | **[0.048; 0.109]** |
| | SD($\alpha$) | **0.012** | **[0.000; 0.028]** | | |
| | SD($\beta$) | **0.010** | **[0.000; 0.025]** | | |
|  | *Model fit* |  |  |  |  |
| | Conditional $R^2$ | **0.766** | **[0.675; 0.831]** | **0.721** | **[0.615; 0.794]** |
| | Marginal $R^2$ | **0.168** | **[0.034; 0.343]** | | |

**Appendix 2—table 2.** Model diagnostics for amplitude (fundamental) per chromatic condition.

| Diagnostic | FLUX | LMS | L-M | MEL | S |
| --- | --- | --- | --- | --- | --- |
| *Model comparison (SMC)* |  |  |  |  |  |
| $\overline{\log Z}$ M0 | 94.20 | 90.03 | 113.23 | 140.54 | 115.53 |
| $SD_{\log Z}$ M0 | 0.42 | 0.30 | 0.10 | 0.17 | 0.25 |
| $\overline{\log Z}$ M1 | 86.47 | 86.44 | 109.05 | 135.46 | 113.00 |
| $SD_{\log Z}$ M1 | 0.29 | 0.24 | 0.14 | 0.05 | 0.16 |
| $\log_{10} BF_{10}$ | -3.36 | -1.56 | -1.81 | -2.21 | -1.10 |
| $SD/\vert \Delta \log Z\vert $ | 0.067 | **0.108** | 0.042 | 0.036 | **0.116** |
| *Convergence* |  |  |  |  |  |
| $\hat{R}_{\max}$ M0 | 1.003 | 1.001 | 1.000 | 1.002 | 1.001 |
| $\hat{R}_{\max}$ M1 | 1.001 | 1.001 | 1.000 | 1.000 | 1.001 |
| *Posterior predictive* |  |  |  |  |  |
| $p_B(\bar{y})$ M0 | 0.497 | 0.496 | 0.501 | 0.502 | 0.503 |
| $p_B(SD)$ M0 | 0.482 | 0.489 | 0.505 | 0.503 | 0.489 |
| $p_B(\bar{y})$ M1 | 0.601 | 0.588 | 0.552 | 0.489 | 0.450 |
| $p_B(SD)$ M1 | 0.544 | 0.554 | 0.617 | 0.621 | 0.547 |
| *Residuals* |  |  |  |  |  |
| Res. mean M0 | 0.000 | 0.000 | 0.000 | 0.000 | 0.000 |
| Res. SD M0 | 0.058 | 0.059 | 0.042 | 0.034 | 0.043 |
| Shapiro $W$ M0 | 0.987 | 0.962 | 0.971 | 0.990 | 0.976 |
| Shapiro $p$ M0 | 0.554 | 0.017 | 0.091 | 0.804 | 0.129 |
| Res. mean M1 | -0.003 | -0.002 | -0.001 | 0.000 | 0.001 |
| Res. SD M1 | 0.052 | 0.050 | 0.040 | 0.032 | 0.038 |
| Shapiro $W$ M1 | 0.984 | 0.964 | 0.967 | 0.986 | 0.968 |
| Shapiro $p$ M1 | 0.407 | 0.025 | 0.050 | 0.514 | 0.042 |
| *Autocorrelation* |  |  |  |  |  |
| ACF lag-1 mean M0 | 0.007 | 0.014 | -0.082 | -0.212 | 0.041 |
| ACF lag-1 SD M0 | 0.152 | 0.186 | 0.188 | 0.352 | 0.242 |
| ACF lag-1 mean M1 | -0.088 | -0.113 | -0.149 | -0.233 | -0.134 |
| ACF lag-1 SD M1 | 0.110 | 0.209 | 0.162 | 0.327 | 0.272 |

*Note.* M0 is the null model, M1 the circadian cosinor model; quantities are reported for both wherever they are defined for each model separately. *Model comparison:* $\overline{\log Z}$ is the mean SMC log marginal likelihood; $SD/|\Delta \log Z|$ is the pooled SD of $\log Z$ across the independent SMC populations divided by the absolute $\log Z$ difference, and values $\geq 0.1$ (bold) mark a numerically imprecise Bayes factor. *Convergence:* $\hat{R}_{\max}$ is the maximum $\hat{R}$ across all parameters, computed across independent SMC populations; it detects disagreement between populations, not within-run mixing, and values $> 1.01$ are shown in bold. *Posterior predictive:* $p_B$ is the Bayesian $p$ -value for the posterior predictive mean and SD; values near 0.5 indicate good fit. *Residuals:* mean and SD of pooled posterior-mean residuals; Shapiro-Wilk $W$ and $p$ test residual normality. *Autocorrelation:* mean and SD across participants of the within-participant residual ACF at a lag of one block (3.75 h); pairs spanning an excluded block are not used. This is descriptive: with roughly ten blocks per participant the lag-1 ACF of uncorrelated residuals is expected near $-1/n$ rather than 0, and a mean near 0 can arise from participant ACFs of opposite sign (see SD).

## Appendix 3

### Complete model output for tonic constriction

This appendix reports the full posterior parameter estimates and numerical diagnostic summary for the revised tonic-minimum outcome analysed in the article. It adds no outcome, model comparison, or hypothesis test beyond those reported in the main text.

**Appendix 3—table 1.** Parameter estimates for circadian cosinor models (Tonic minimum) across conditions. Values are posterior means with 95% credible intervals, except *R*^2^, reported as posterior median with a 95% equal-tailed interval. M1 = cosinor model; M0 = null (intercept-only) model.

| Condition | Model parameter | M1 (cosinor) |  | M0 (null) |  |
| --- | --- | --- | --- | --- | --- |
|  |  | Estimate | 95% CI | Estimate | 95% CI |
| FLUX | *Fixed effects* |  |  |  |  |
|  | Intercept | **-0.447** | **[-0.590; -0.309]** | **-0.445** | **[-0.599; -0.296]** |
| | $\alpha$ (sin) | -0.065 | [-0.136; 0.004] | | |
| | $\beta$ (cos) | 0.016 | [-0.043; 0.074] | | |
|  | *Random effects* |  |  |  |  |
|  | SD(Intercept) | **0.217** | **[0.129; 0.304]** | **0.228** | **[0.134; 0.332]** |
| | SD($\alpha$) | **0.054** | **[0.001; 0.122]** | | |
| | SD($\beta$) | **0.034** | **[0.001; 0.085]** | | |
|  | *Model fit* |  |  |  |  |
| | Conditional $R^2$ | **0.621** | **[0.488; 0.721]** | **0.587** | **[0.446; 0.690]** |
| | Marginal $R^2$ | **0.073** | **[0.005; 0.222]** | | |
| L-M | *Fixed effects* |  |  |  |  |
|  | Intercept | **-0.920** | **[-1.148; -0.692]** | **-0.924** | **[-1.184; -0.682]** |
| | $\alpha$ (sin) | 0.041 | [-0.054; 0.134] | | |
| | $\beta$ (cos) | **0.167** | **[0.086; 0.252]** | | |
|  | *Random effects* |  |  |  |  |
|  | SD(Intercept) | **0.332** | **[0.203; 0.478]** | **0.351** | **[0.194; 0.518]** |
| | SD($\alpha$) | **0.055** | **[0.001; 0.140]** | | |
| | SD($\beta$) | **0.051** | **[0.001; 0.127]** | | |
|  | *Model fit* |  |  |  |  |
| | Conditional $R^2$ | **0.699** | **[0.579; 0.782]** | **0.594** | **[0.443; 0.702]** |
| | Marginal $R^2$ | **0.248** | **[0.081; 0.428]** | | |
| LMS | *Fixed effects* |  |  |  |  |
|  | Intercept | **-0.531** | **[-0.716; -0.337]** | **-0.534** | **[-0.746; -0.321]** |
| | $\alpha$ (sin) | 0.034 | [-0.040; 0.108] | | |
| | $\beta$ (cos) | 0.029 | [-0.040; 0.094] | | |
|  | *Random effects* |  |  |  |  |
|  | SD(Intercept) | **0.299** | **[0.197; 0.409]** | **0.323** | **[0.196; 0.454]** |
| | SD($\alpha$) | **0.050** | **[0.001; 0.120]** | | |
| | SD($\beta$) | **0.041** | **[0.001; 0.102]** | | |
|  | *Model fit* |  |  |  |  |
| | Conditional $R^2$ | **0.743** | **[0.642; 0.813]** | **0.739** | **[0.639; 0.809]** |
| | Marginal $R^2$ | **0.046** | **[0.002; 0.180]** | | |
| MEL | *Fixed effects* |  |  |  |  |
|  | Intercept | **-0.445** | **[-0.623; -0.265]** | **-0.442** | **[-0.623; -0.228]** |
| | $\alpha$ (sin) | 0.023 | [-0.079; 0.130] | | |
| | $\beta$ (cos) | 0.023 | [-0.062; 0.106] | | |
|  | *Random effects* |  |  |  |  |
|  | SD(Intercept) | **0.270** | **[0.161; 0.391]** | **0.290** | **[0.163; 0.423]** |
| | SD($\alpha$) | **0.097** | **[0.002; 0.199]** | | |
| | SD($\beta$) | **0.055** | **[0.001; 0.133]** | | |
|  | *Model fit* |  |  |  |  |
| | Conditional $R^2$ | **0.575** | **[0.425; 0.687]** | **0.546** | **[0.392; 0.662]** |
| | Marginal $R^2$ | **0.032** | **[0.001; 0.154]** | | |
| S | *Fixed effects* |  |  |  |  |
|  | Intercept | **-0.599** | **[-0.781; -0.409]** | **-0.601** | **[-0.800; -0.405]** |
| | $\alpha$ (sin) | 0.024 | [-0.064; 0.109] | | |
| | $\beta$ (cos) | 0.061 | [-0.018; 0.144] | | |
|  | *Random effects* |  |  |  |  |
|  | SD(Intercept) | **0.287** | **[0.175; 0.407]** | **0.298** | **[0.175; 0.425]** |
| | SD($\alpha$) | **0.056** | **[0.001; 0.136]** | | |
| | SD($\beta$) | **0.053** | **[0.001; 0.125]** | | |
|  | *Model fit* |  |  |  |  |
| | Conditional $R^2$ | **0.623** | **[0.488; 0.721]** | **0.600** | **[0.461; 0.703]** |
| | Marginal $R^2$ | **0.059** | **[0.003; 0.204]** | | |

**Appendix 3—table 2.**
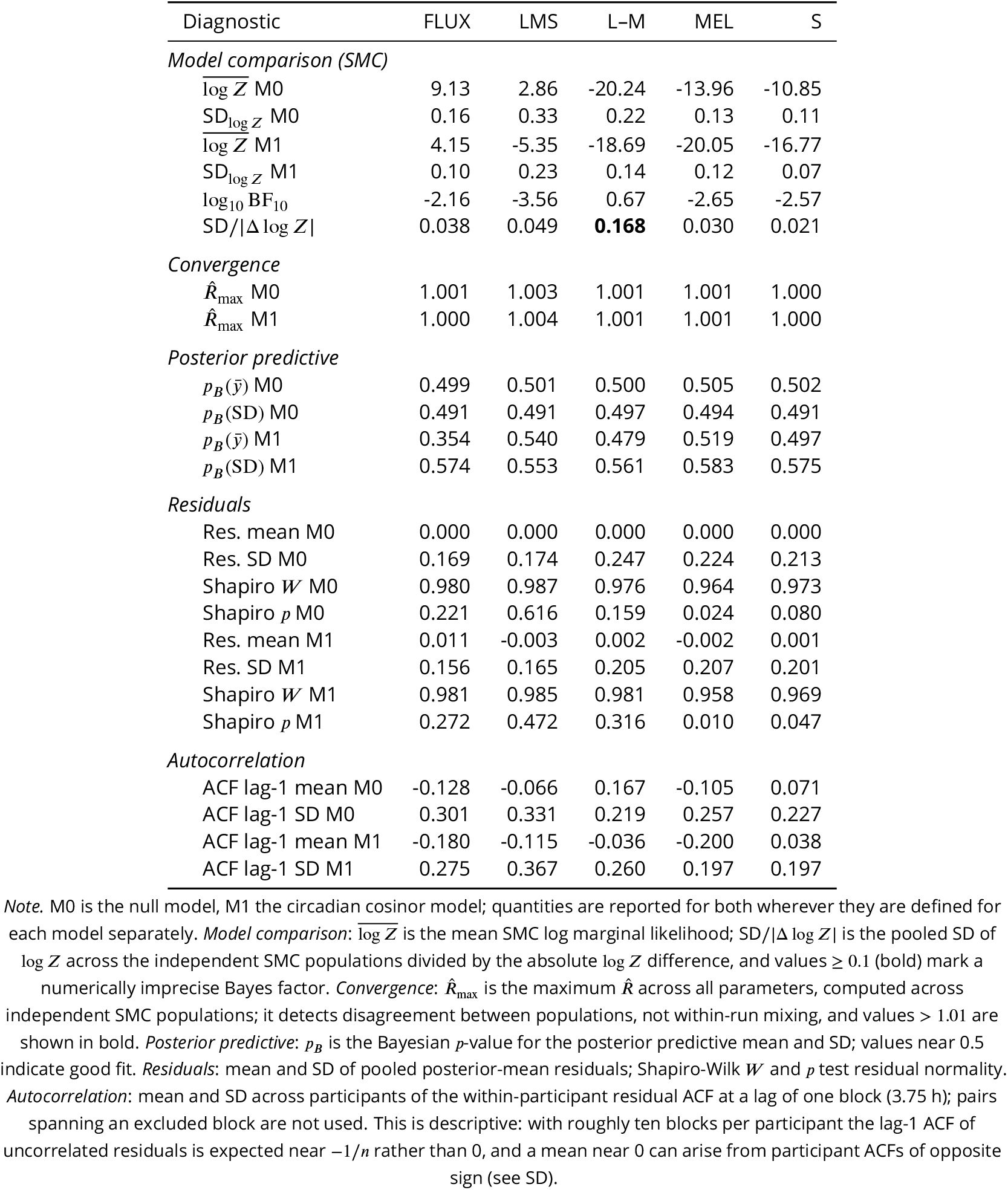
Model diagnostics for tonic minimum per chromatic condition.

## Appendix 4

### The registered pupil outcome

Appendix 1—table 1 records that the published protocol (***Heinrichs and Spitschan, 2025b***) specified a different primary pupil outcome from the one analysed in the main article. That outcome is the maximum pupil-constriction amplitude for each condition normalised to the corresponding broadband FLUX response, which the protocol denotes *R* and terms the relative pupil constriction:

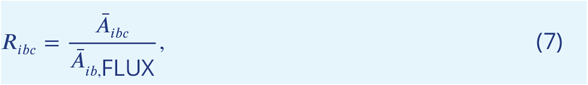

where *Ā*_*bc*_ is the mean maximum pupil-constriction amplitude for participant *i*, measurement block *b* and condition *c*. This appendix analyses that outcome directly and characterises what it measures in these data.

#### Derivation

The derivation starts from the same cleaned traces and the same quality control as the analyses in the main article. It differs in the outcome computed from them and in the order of baseline correction: here each trial is corrected to its own baseline before averaging, whereas the main analysis corrects the trial-averaged trace. Maximum constriction was obtained separately for each accepted trial as the difference between the pre-stimulus mean over 1 second and the smallest diameter recorded during the 5-s stimulus, so that constriction is positive. Trial amplitudes were then averaged over all accepted trials in a retained block, pooling all runs recorded within it. Each non-FLUX block amplitude was divided by the FLUX amplitude from the same participant and the same block, and the requirement of at least six blocks per participant and condition was applied after that pairing. The FLUX condition is not itself modelled, since *Ā*_FLUX_/*Ā*_FLUX_ = 1 by construction.

FLUX was recorded in every one of the 95 retained blocks that hold any data, so every non-FLUX block had a same-block FLUX denominator and pairing removed none. The six-block requirement then removed 7–15 blocks per condition (fewest for S, most for L–M), leaving 8 participants (7 for L–M) and 74–81 block-level observations per condition: the same blocks as in the main-article phasic analysis of these four conditions. No block-level amplitude was at or below zero, and the smallest FLUX denominator was 0.363 mm, so no observation required special handling.

#### What this outcome measures

The registered outcome is not a less precise version of the phasic amplitude analysed in the main article. It measures something different, and Appendix 4—figure 1 shows what.

Each trial comprises a large sustained constriction out of the white inter-trial background with the 0.4-Hz modulation superimposed on it. A maximum taken over the whole stimulus reads the single deepest sample and is therefore dominated by the sustained component. The consequence is visible in the condition ordering: the L–M condition produces the largest maximum constriction of any condition (1.239 mm, against 0.806 mm for MEL and 0.996 mm for FLUX), yet in the phasic analysis reported in the main article it has the second-smallest fitted amplitude at the stimulus frequency (0.12 mm, above only MEL at 0.10 mm). The two outcomes therefore order the conditions differently, which would not be expected if they measured the same quantity with differing precision.

Maximum constriction does track the stimulus: the minima cluster descriptively at the troughs of the pupil response to the 0.4-Hz modulation. The clustering differs by condition, however, and is weakest for L–M, so the endpoint is not a clean measure of oscillatory depth. Taken together, these observations are the reason the main article decomposes the averaged trace into tonic and phasic components rather than analysing maximum constriction.

#### Order of operations

The protocol specifies that the maximum is taken on each trial and the trial amplitudes are then averaged. Reversing the two steps – averaging the trials into one trace and taking a single maximum from it – changes the block-level amplitude by only 0.060–0.146 mm, and both routes yield values near 1 mm, so the choice is not visible in the result. Both quantities were therefore computed for every retained block and are reported in Appendix 4—figure 2.

On average, the per-trial route gave a deeper endpoint by 4.865–18.134% depending on condition. Averaged within participants, the standard deviation of *R* across blocks was 0.373–0.525 for the registered ordering and 0.412–0.710 for the alternative, and the two orderings gave values correlated at *r* = 0.830–0.917.

#### Circadian models

The outcome *R* was entered into the same constant-level model M0 and cosinor model M1 used for the revised outcomes in the main article, fitted separately for each non-FLUX condition with participant-specific periods from the salivary melatonin cosinor and the same sequential Monte Carlo settings. Full parameter estimates appear in Appendix 4—table 2, numerical diagnostics in Appendix 4—table 3, and posterior fits against the observed series in Appendix 4—figure 5.

Evidence favoured the constant-level model in every condition, with log_10_ BF_10_ ranging from −2.09 (LMS) to −0.32 (L–M). The numerical precision ratio for L–M was 0.15, above the pragmatic 0.10 threshold, so the magnitude of this Bayes factor is imprecise. The comparison against the two revised outcomes is shown in Appendix 4—figure 4. Every ratio and phasic-amplitude comparison therefore favours M0, so the phasic conclusion of the main article does not depend on which of these two outcomes is analysed; the tonic minimum in L–M is the one exception across the three outcomes and is discussed below.

One difference is worth recording. The tonic minimum is the only outcome that favours a circadian effect in the L–M condition, and the registered outcome does not reproduce it. Maximum constriction is dominated by the sustained component, but *R* takes it from unfiltered single trials and divides it by the same-block FLUX response, so it is not a direct replication of the tonic minimum. The discordance nevertheless suggests that the tonic result should be treated as provisional rather than corroborated, particularly because the precision ratios of both L–M Bayes factors exceed the 0.10 threshold (0.17 for the tonic minimum, 0.15 for *R*).

#### Sensitivity analyses and limitations

The sensitivity analyses reported here, all specified before fitting, were a log transform of *R*, the averaged-trace ordering, the period range of 20–28 h given in the protocol, and leave-one-participant-out refits. The averaged-trace ordering was compared only descriptively (Order of operations, above).

##### Outcome transformation

The raw ratio is right-skewed and residual normality is rejected in every condition, so the Gaussian likelihood cannot be assumed adequate. Under a log transform every Bayes factor kept its direction, with broadly similar magnitudes (Appendix 4—table 1). Residual normality was no longer rejected for L–M and S but was still rejected for LMS and MEL, so the transform does not by itself establish that a Gaussian likelihood is adequate.

**Appendix 1—table 1.** Comparison of the published protocol with the analyses reported here. These differences mean that the 0.4-Hz phasic-amplitude and tonic-minimum analyses should not be interpreted as confirmatory tests of the protocol’s primary pupil outcome.

| Aspect | Published protocol | Present article | Consequence for interpretation |
| --- | --- | --- | --- |
| Pupil outcome | Maximum constriction during the 5-s stimulus, normalised to the broadband FLUX response | Unnormalised 0.4-Hz phasic amplitude and tonic minimum | Revised outcomes; neither is the protocol's primary outcome. The registered outcome is analysed in Appendix 4; like the phasic amplitude, it favours the constant-level model in every condition |
| Melatonin model | Single-harmonic cosinor | Bi-harmonic cosinor | Additional fitted harmonic |
| Candidate period | 20–28 h | 22–28 h | Narrower search range. Widening it to 20–28 h selects the same period for every participant and leaves the results unchanged (Appendix 4) |
| Stopping | Sequential Bayes-factor rule based on the MEL response, with a maximum sample of 12 | The article reports 12 completers and eight participants after quality control; the sequential rule was not applied, because evaluating the evidence during data collection was not feasible alongside the demands of running the protocol | Fixed sample; recruitment did not depend on interim evidence |
| Rod control | Rod-escape control planned after the main data collection | Not included in this report | Photoreceptor directions remain nominal; rod intrusion is not empirically excluded here |

##### Period-search range

This check retains all participants and changes only the period-search range; the two-harmonic melatonin model is retained.

Widening the period search from 22–28 h to the registered 20–28 h changed no result at all. Every participant’s best-fitting period already lay between 23.1 and 25.2 h, so the wider range selected the same period in every case and the Bayes factors are identical to three decimal places. The narrowed range recorded in Appendix 1—table 1 therefore had no bearing on these data.

##### Participant influence

The leave-one-participant-out (LOPO) check instead retains the period estimates from the usual 22–28 h search and omits one participant per refit of the raw-ratio model. It is separate from the period-range check above.

The L–M condition alone warrants a further check, because its evidence lies close to the point of indifference (log_10_ BF_10_ = −0.32). Refitting with each participant held out in turn moves log_10_ BF_10_ from values favouring M0 to a value slightly favouring M1; omitting participant 204 alone reverses the sign, so a single participant determines which model is favoured for *R*. No conclusion about circadian modulation of L–M should therefore be drawn from the ratio outcome. Participant influence was not assessed for the other three conditions or for the phasic-amplitude and tonic-minimum models.

##### Limitations

Several limitations follow from the definition in Equation 7 rather than from the data. A change in *R* may arise from the numerator, the denominator, or both, and changes shared between a condition and FLUX cancel. The four condition ratios share the same FLUX observations and are therefore not four independent tests. Inference rests on seven or eight participants.

Finally, restoring the registered outcome does not make this a confirmatory analysis. The sequential stopping rule cannot be reconstructed retrospectively, the priors used here were not specified in the protocol, and the single-harmonic melatonin model recorded in Appendix 1—table 1 was not restored; only the period range was. These analyses were carried out after the results of the main analysis were known.

**Appendix 4—table 1.** Sensitivity of the FLUX-normalised maximum constriction (*R*) to a log transform. log_10_ BF_10_ compares the circadian model M1 with the constant-level model M0; negative values favour M0. Shapiro–Wilk *p* refers to the M1 residuals.

| *Condition* | $\log_{10} \text{BF}_{10}$ | | Shapiro–Wilk $p$ | |
| --- | --- | --- | --- | --- |
| | $R$ | $\log R$ | $R$ | $\log R$ |
| LMS | −2.09 | −2.14 | 0.00 | 0.01 |
| L–M | −0.32 | −0.55 | 0.04 | 0.13 |
| MEL | −1.38 | −1.31 | 0.01 | 0.00 |
| S | −1.58 | −1.31 | 0.00 | 0.98 |

**Appendix 4—table 2.**
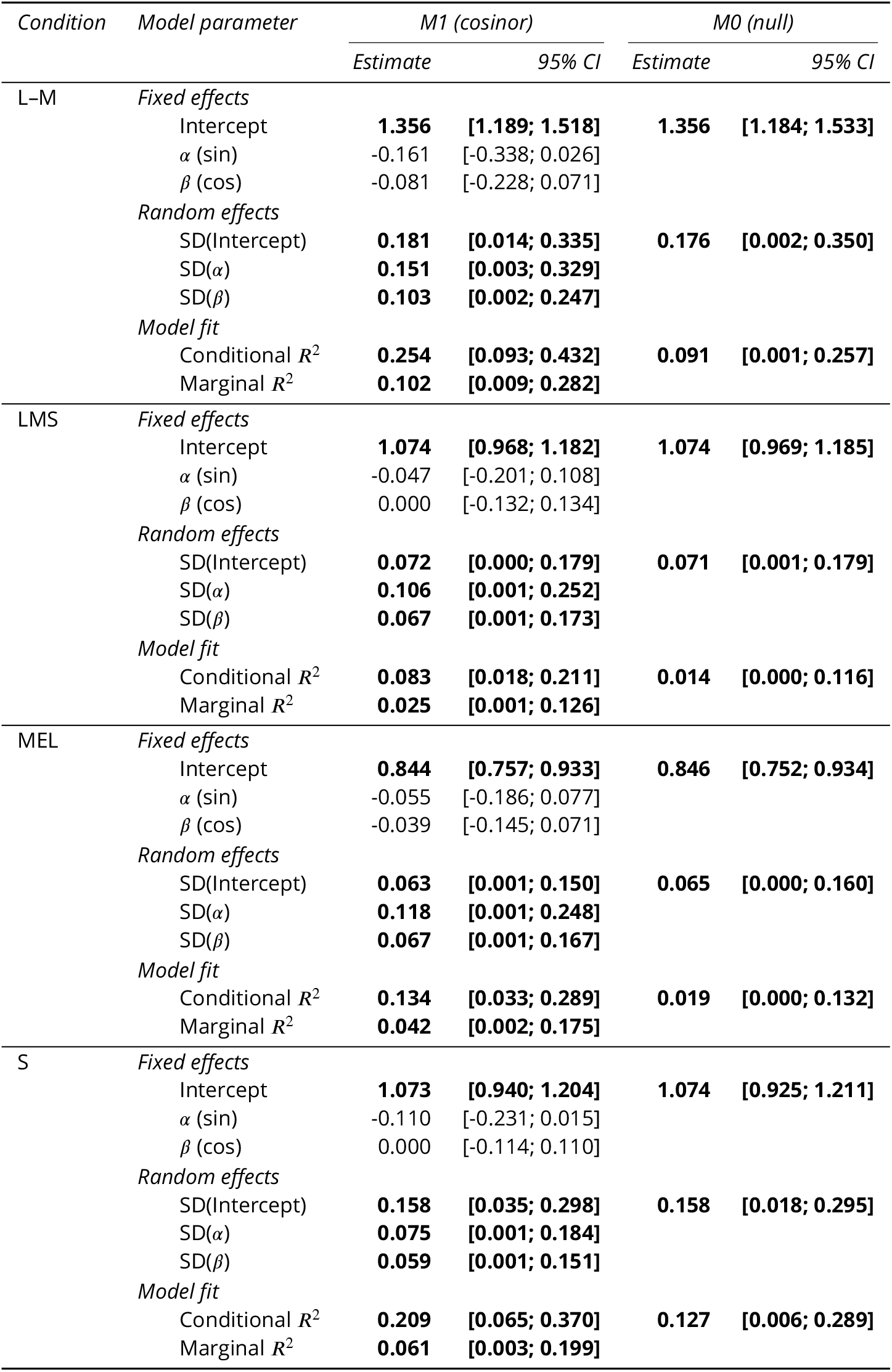
Parameter estimates for circadian cosinor models (Protocol ratio (condition / Flux)) across conditions. Values are posterior means with 95% credible intervals, except *R*^2^, reported as posterior median with a 95% equal-tailed interval. M1 = cosinor model; M0 = null (intercept-only) model.

**Appendix 4—table 3.** Model diagnostics for protocol ratio (condition / flux) per chromatic condition.

| Diagnostic | LMS | L-M | MEL | S |
| --- | --- | --- | --- | --- |
| *Model comparison (SMC)* |  |  |  |  |
| $\log \bar{Z}$ M0 | -45.17 | -49.89 | -28.18 | -34.95 |
| $SD_{\log Z}$ M0 | 0.01 | 0.04 | 0.04 | 0.06 |
| $\log \bar{Z}$ M1 | -49.98 | -50.63 | -31.36 | -38.59 |
| $SD_{\log Z}$ M1 | 0.02 | 0.10 | 0.07 | 0.08 |
| $\log_{10} BF_{10}$ | -2.09 | -0.32 | -1.38 | -1.58 |
| $SD/\vert \Delta \log Z\vert $ | 0.004 | **0.152** | 0.025 | 0.026 |
| *Convergence* |  |  |  |  |
| $\hat{R}_{\max}$ M0 | 1.000 | 1.002 | 1.000 | 1.002 |
| $\hat{R}_{\max}$ M1 | 1.000 | 1.001 | 1.001 | 1.000 |
| *Posterior predictive* |  |  |  |  |
| $p_B(\bar{y})$ M0 | 0.503 | 0.503 | 0.502 | 0.508 |
| $p_B(SD)$ M0 | 0.538 | 0.529 | 0.535 | 0.520 |
| $p_B(\bar{y})$ M1 | 0.458 | 0.379 | 0.463 | 0.372 |
| $p_B(SD)$ M1 | 0.679 | 0.639 | 0.667 | 0.653 |
| *Residuals* |  |  |  |  |
| Res. mean M0 | 0.000 | 0.000 | 0.000 | -0.001 |
| Res. SD M0 | 0.386 | 0.413 | 0.311 | 0.321 |
| Shapiro $W$ M0 | 0.901 | 0.953 | 0.951 | 0.950 |
| Shapiro $p$ M0 | 0.000 | 0.007 | 0.004 | 0.003 |
| Res. mean M1 | 0.007 | 0.021 | 0.005 | 0.018 |
| Res. SD M1 | 0.376 | 0.362 | 0.289 | 0.305 |
| Shapiro $W$ M1 | 0.900 | 0.965 | 0.953 | 0.946 |
| Shapiro $p$ M1 | 0.000 | 0.039 | 0.005 | 0.002 |
| *Autocorrelation* |  |  |  |  |
| ACF lag-1 mean M0 | -0.223 | 0.021 | 0.045 | -0.113 |
| ACF lag-1 SD M0 | 0.233 | 0.223 | 0.199 | 0.319 |
| ACF lag-1 mean M1 | -0.279 | -0.090 | -0.049 | -0.142 |
| ACF lag-1 SD M1 | 0.243 | 0.260 | 0.216 | 0.336 |

**Appendix 4—figure 1.**
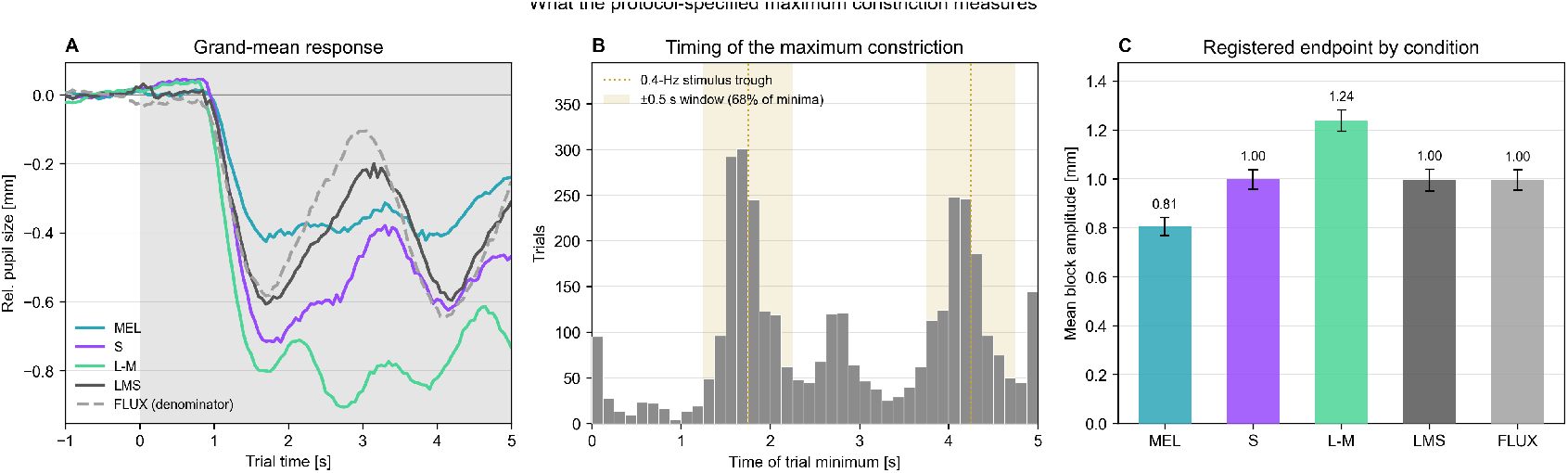
Maximum constriction is dominated by sustained constriction. All panels use the analysis sample of the main article: retained blocks from participant-condition series with at least six blocks (8 participants; 7 for L–M). (**A**) Grand-mean relative pupil size, each trial corrected to its own pre-stimulus mean over 1 second, across the 0–5 s window in which the maximum is sought. Shading marks the stimulus. FLUX, the denominator of Equation 7, is drawn dashed. The L–M condition constricts deepest and recovers most slowly while oscillating least. (**B**) Distribution of the time at which the minimum occurs, one row per condition. Dotted lines mark troughs of the pupil response, set at the first trough of the grand-mean trace in panel A (about 1.75 s) and one stimulus period later (4.25 s); they differ from the stimulus contrast troughs at 1.875 and 4.375 s. Shaded bands mark the 0.5-s window on either side of each; together the windows cover 40% of the stimulus window. The percentage in each row is the share of that condition’s trial minima within these windows. Because the troughs were read from the same trials, these shares are descriptive; 40% is the share expected if minima were spread evenly across the window. The share ranges from 46% for L–M to 85% for FLUX (68% of all 3562 trial minima), so the endpoint tracks the stimulus more closely in some conditions than in others. (**C**) Mean amplitude by condition. Blocks are first averaged within each participant; bars show the mean of these participant means, dots the participant means, and error bars the standard error across participants. The ordering differs from that of the phasic amplitudes reported in the main article, most markedly for L–M.

**Appendix 4—figure 2.**
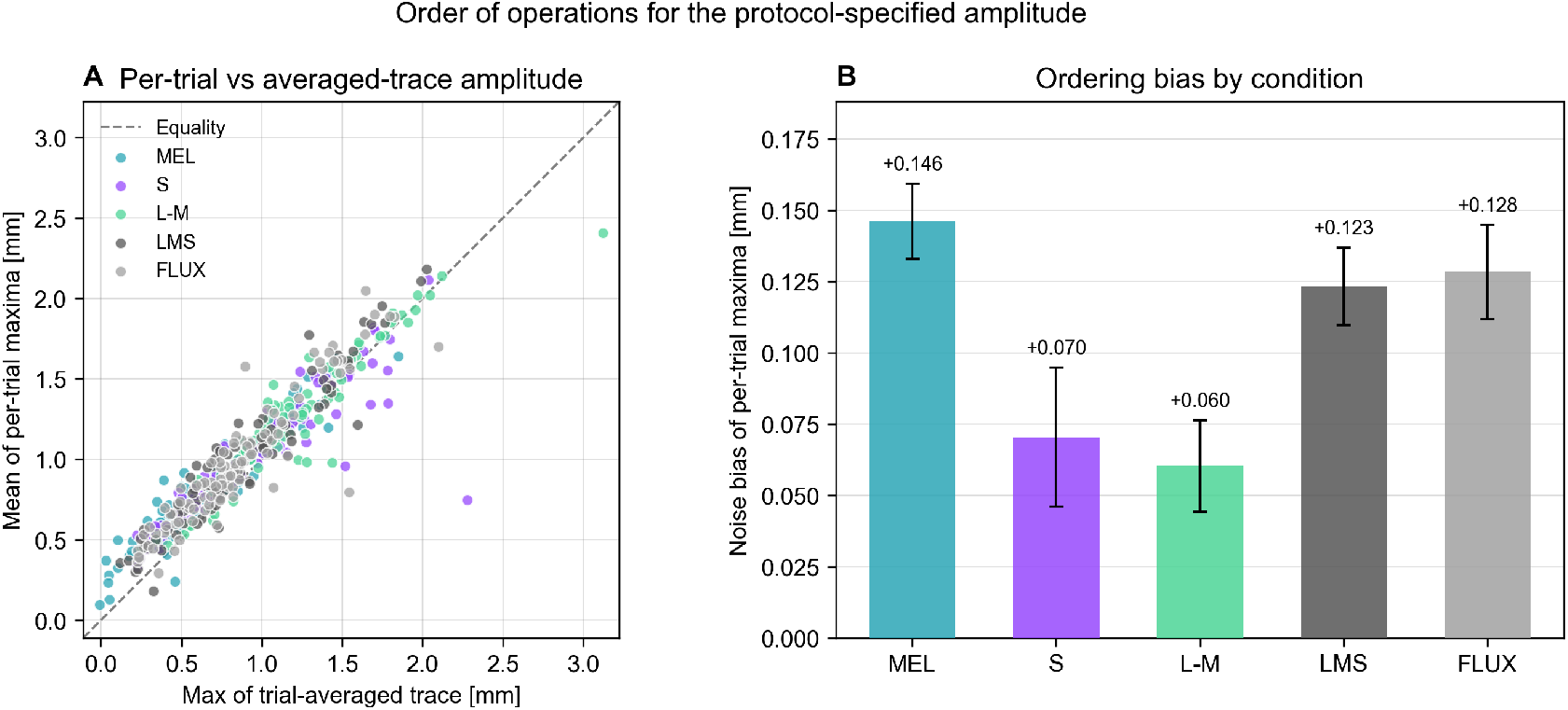
The order of operations is not visible in the result but is not neutral. Both panels use the same analysis sample as Appendix 4—figure 1. (**A**) Block-level amplitude obtained by averaging per-trial maxima, against the maximum of the trial-averaged trace, shown separately for MEL, S, L–M, LMS and FLUX. Each point represents one participant and block within a condition. All panels share the same axis limits; dashed lines mark equality. (**B**) Mean displacement between the two, by condition. Blocks are first averaged within each participant; coloured points show the mean of these participant means with the standard error across participants, and grey dots the participant means. On average the per-trial maximum is deeper in every condition, although not for every participant, and the mean difference varies 2.423-fold across conditions.

**Appendix 4—figure 3.**
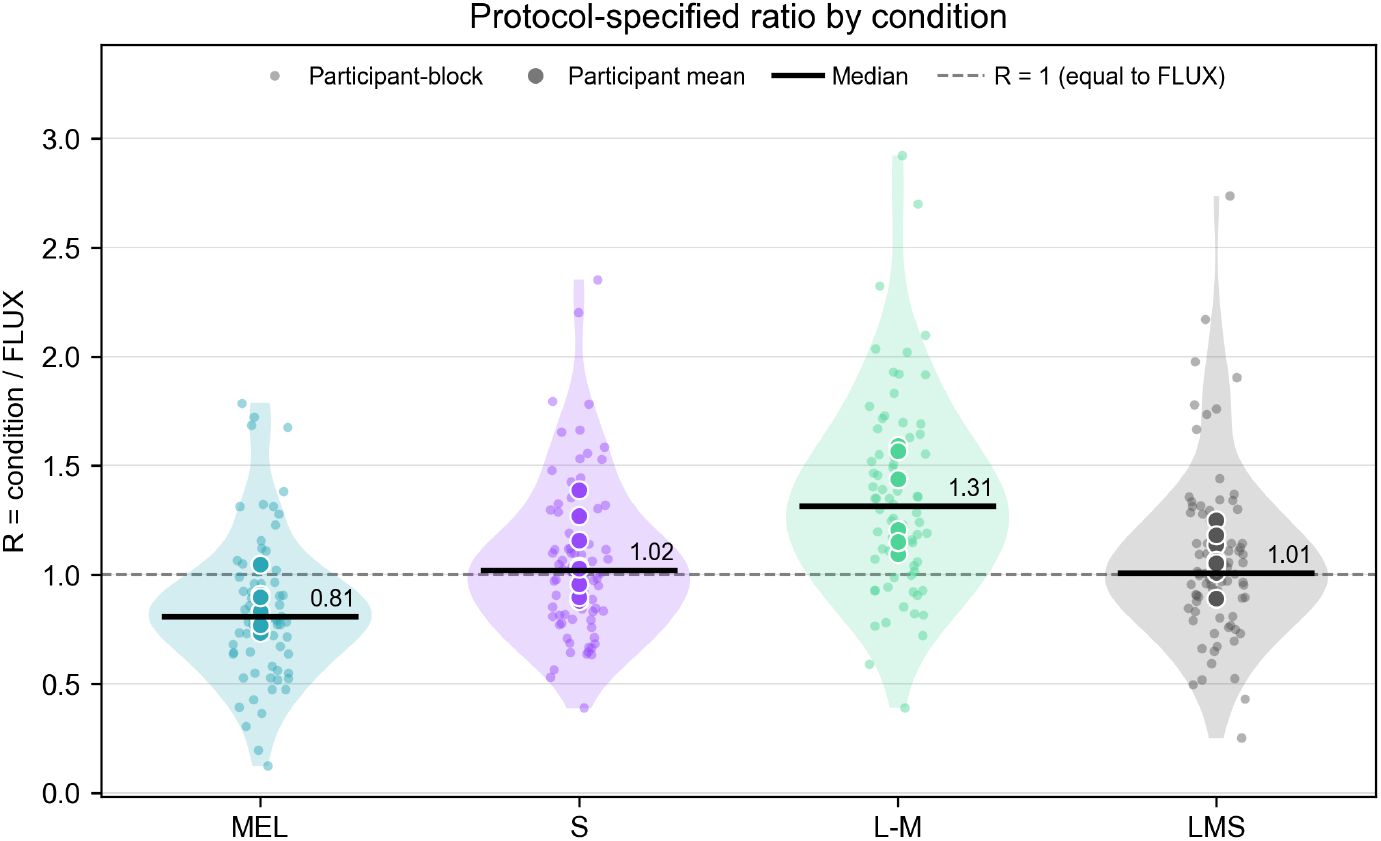
Distribution of FLUX-normalised maximum constriction (*R*). Half-violins show the distribution of retained block-level observations; small circles show individual blocks, diamonds participant means, and short horizontal lines the condition medians across blocks. The dashed line marks *R* = 1, where a condition evokes the same maximum constriction as FLUX in the same block.

**Appendix 4—figure 4.**
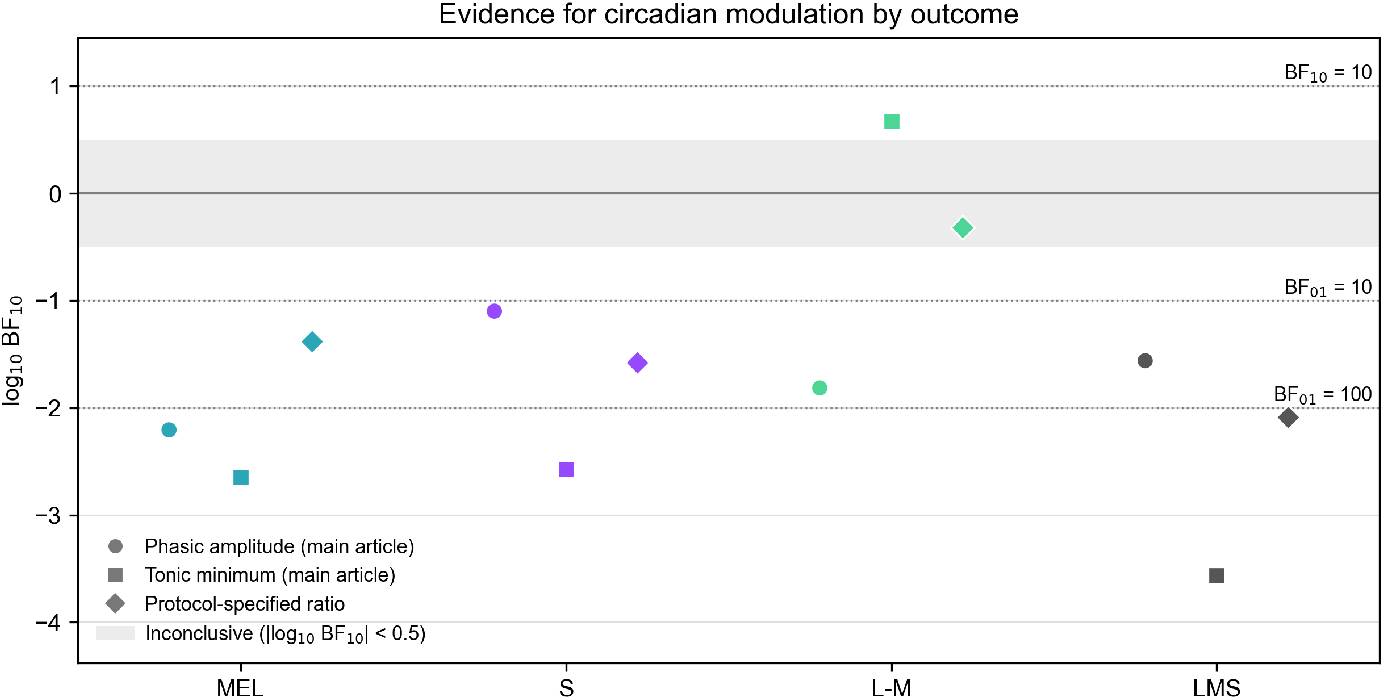
Evidence for circadian modulation under three pupil outcomes. Each row is one condition; marker shape denotes the outcome, and the line spans the three estimates. Negative values favour the constant-level model M0. Whiskers show ±1 SD of the log marginal-likelihood difference across SMC chains, which reflects Monte Carlo variability only; open markers flag estimates whose SD/|Δ log *Z*| exceeds the pragmatic 0.10 threshold. Evidence favours the constant-level model M0 for FLUX-normalised maximum constriction in every condition, as it does for the phasic amplitude. The axis is log_10_ BF_10_ and therefore unitless; the underlying amplitudes are in different units and are not plotted on a common scale.

**Appendix 4—figure 5.**
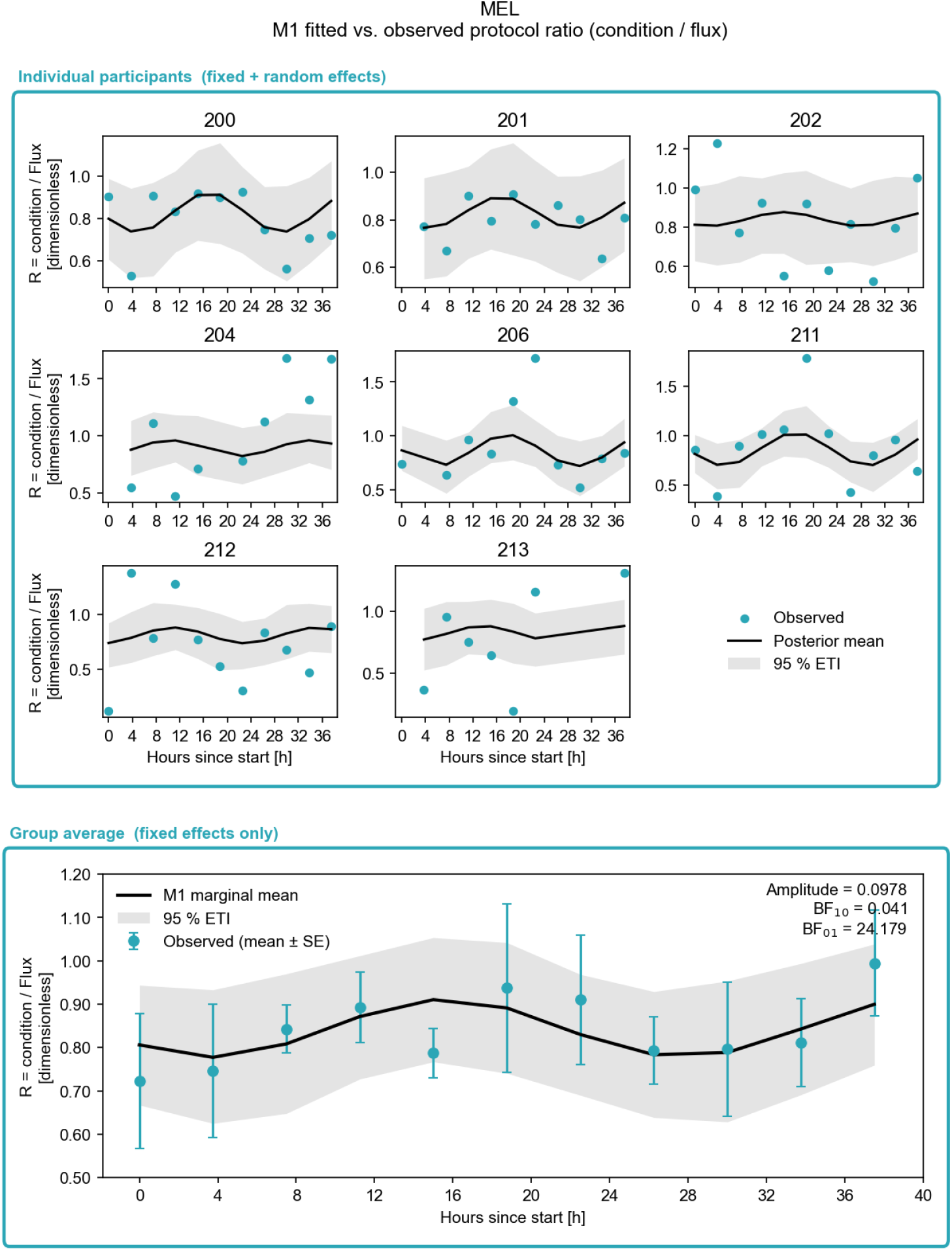
Cosinor model fitted to FLUX-normalised maximum constriction (*R*), MEL condition. Upper panels show each participant’s observed ratios with the conditional posterior mean and 95% equal-tailed interval; the lower panel shows block means with standard errors against the marginal posterior. Remaining conditions are available in the accompanying analysis outputs.

**4—figure supplement 1.**
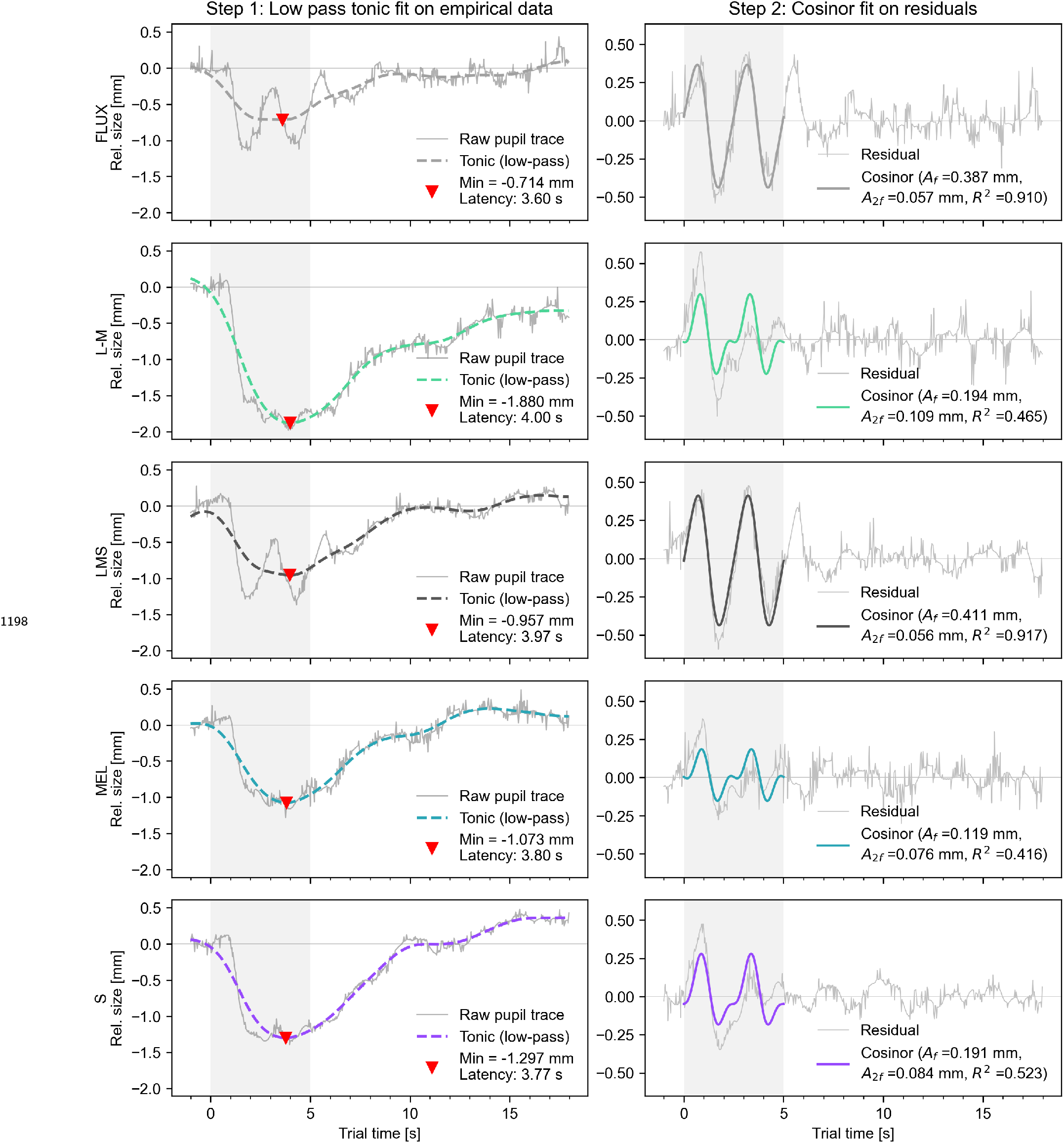
Two-step decomposition for participant 202. Each row represents one stimulus condition. Left panels show the empirical baseline-corrected pupil trace and zero-phase low-pass tonic component; right panels show the residual and fitted bi-harmonic phasic component. The grey interval marks the 5-s stimulus.

**4—figure supplement 2.**
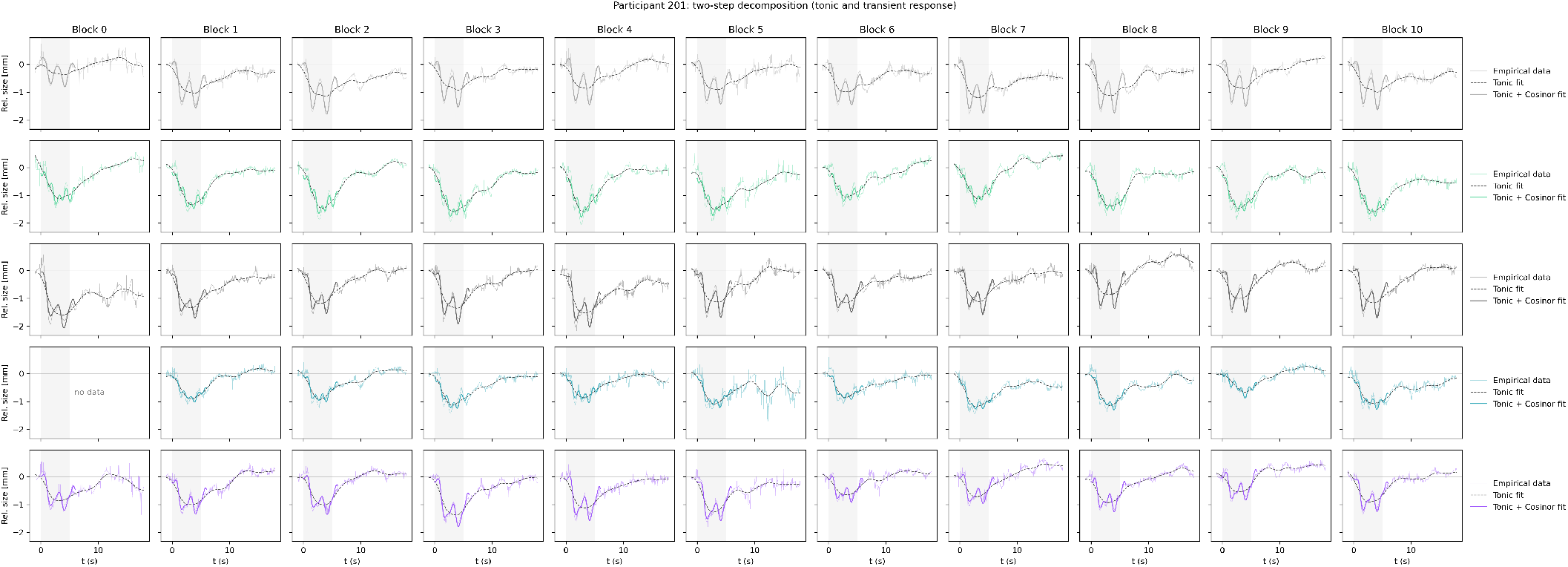
Block-level pupil-response traces for participant 201. Panels show retained trial-averaged traces by stimulus condition and protocol block. The grey interval marks the 5-s stimulus.

**4—figure supplement 3.**
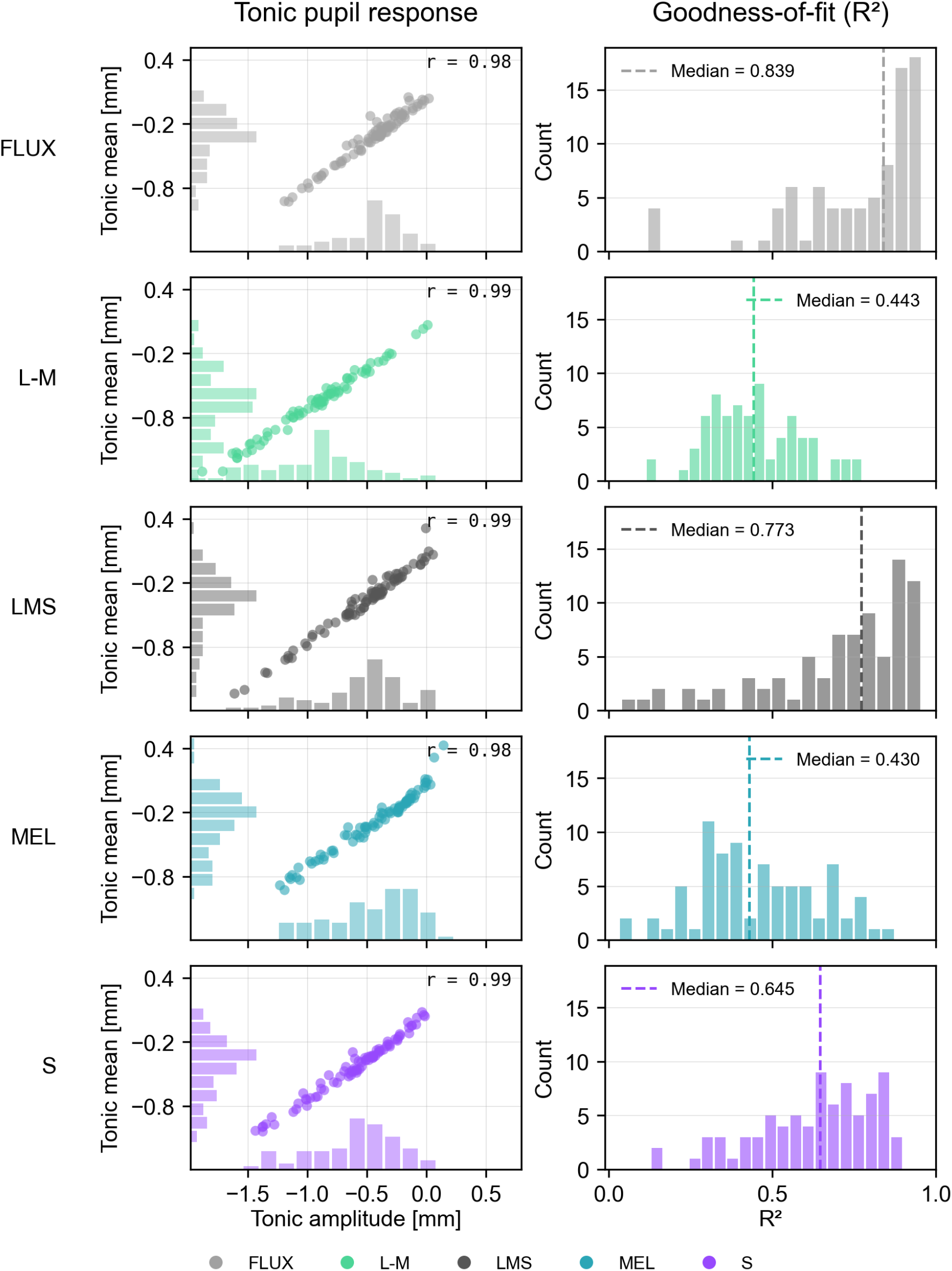
Rows show the five stimulus conditions, colour-coded as in the legend below. Left column: relationship between tonic mean and tonic minimum from the low-pass tonic step. Points are retained participant-by-condition-by-block traces; Pearson correlations are descriptive and do not account for repeated observations nested within participants. Marginal histograms show the corresponding univariate distributions. Right column: distribution of adjusted *R*^2^ for the bi-harmonic phasic fits from the cosinor step. Histograms pool retained participant-by-condition-by-block fits within each stimulus condition; dashed lines mark condition medians. Retained trace counts were 83 for FLUX (eight participants), 74 for L–M (seven), 80 for LMS (eight), 80 for MEL (eight), and 81 for S (eight).

**4—figure supplement 4.**
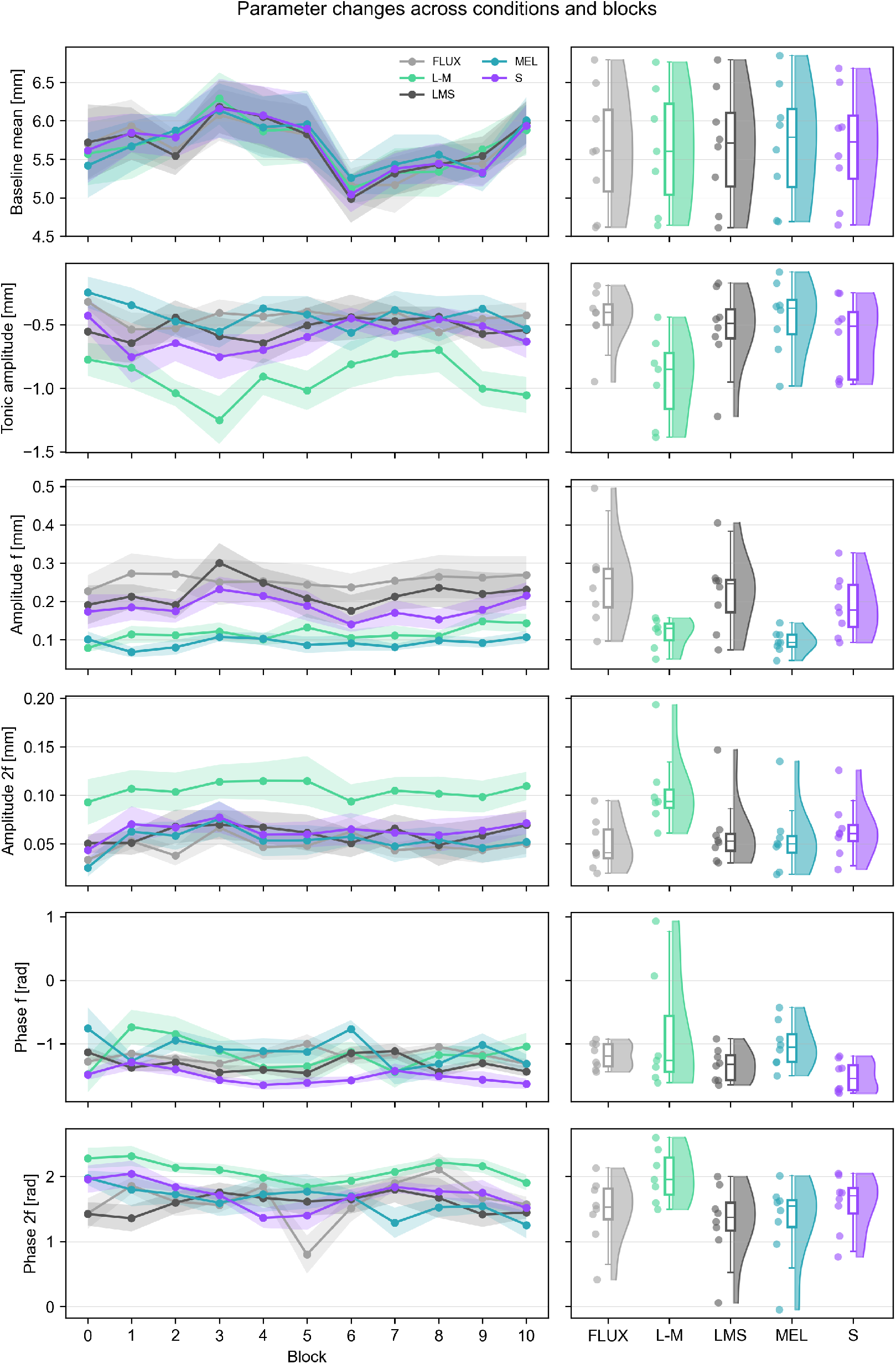
Extracted pupil parameters across protocol blocks. Rows show baseline pupil diameter, tonic minimum, fundamental and second-harmonic phasic amplitudes, and the corresponding phases; columns separate temporal trajectories from participant-level distributions by stimulus condition. Lines and points with bands in the left panels show participant means and standard errors. Phase summaries use circular statistics.

**4—figure supplement 5.**
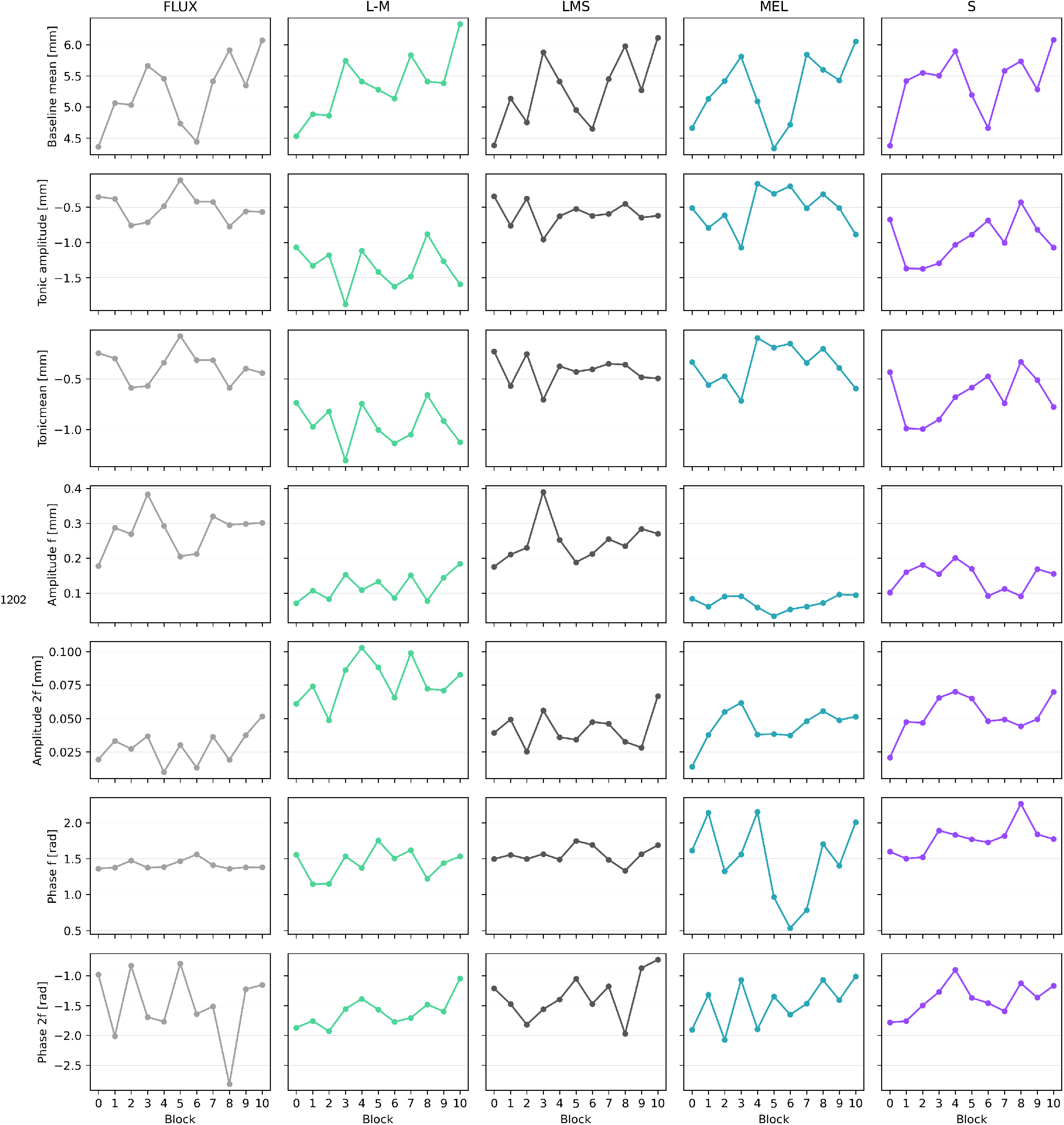
Extracted pupil parameters across protocol blocks for participant 202. Rows show baseline pupil diameter, tonic minimum, fundamental and second-harmonic phasic amplitudes, and the corresponding phases; columns show the five stimulus conditions.

**5—figure supplement 1.**
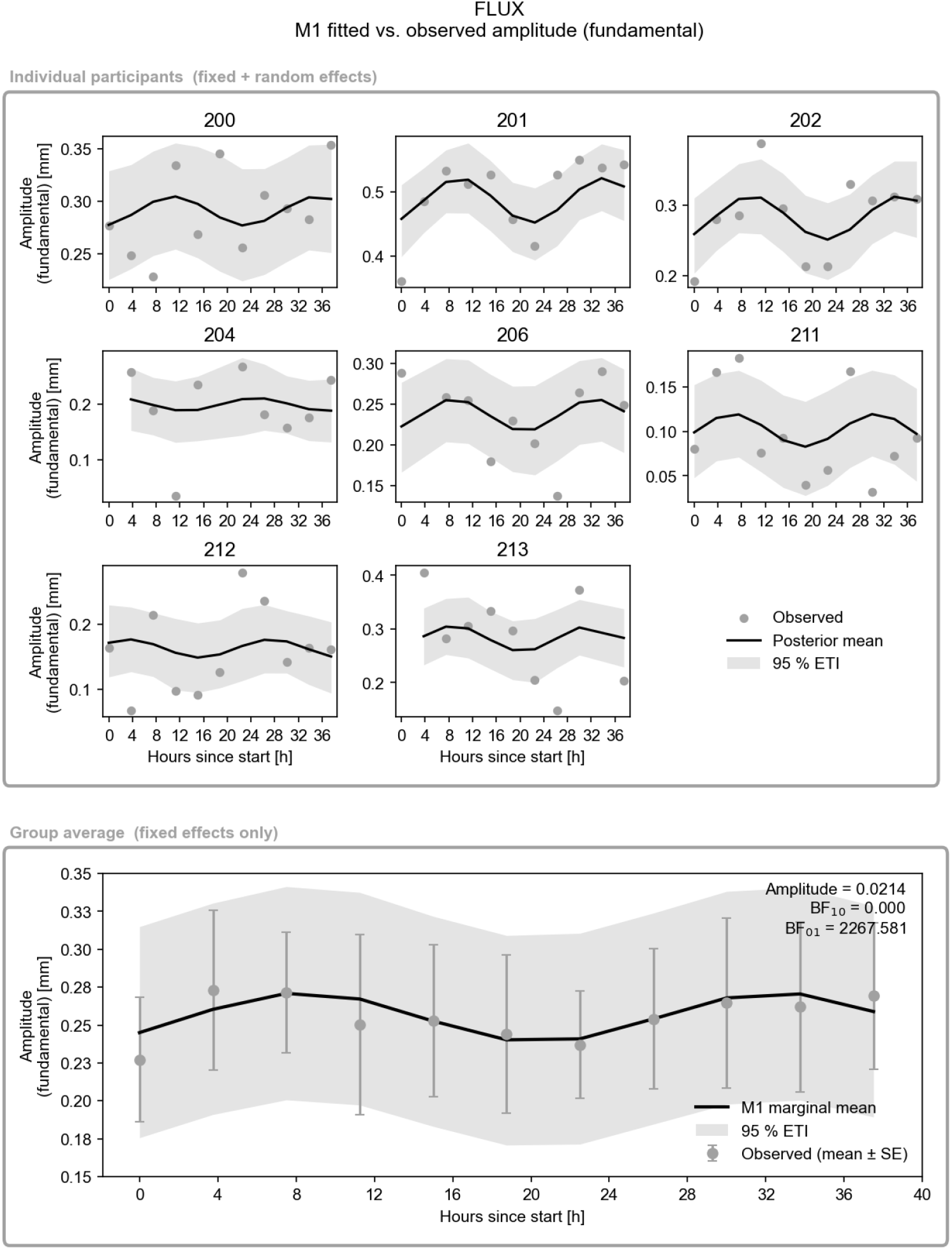
Participant-level M1 fits for phasic amplitude in the broadband FLUX condition (83 retained block-level estimates from eight participants). In the participant panels, points are observed estimates, black lines are conditional posterior means, and grey ribbons are 95% equal-tailed intervals. In the group panel, points and error bars are block means and standard errors, the black line is the fixed-effect posterior mean, and the grey ribbon is its 95% equal-tailed interval.

**5—figure supplement 2.**
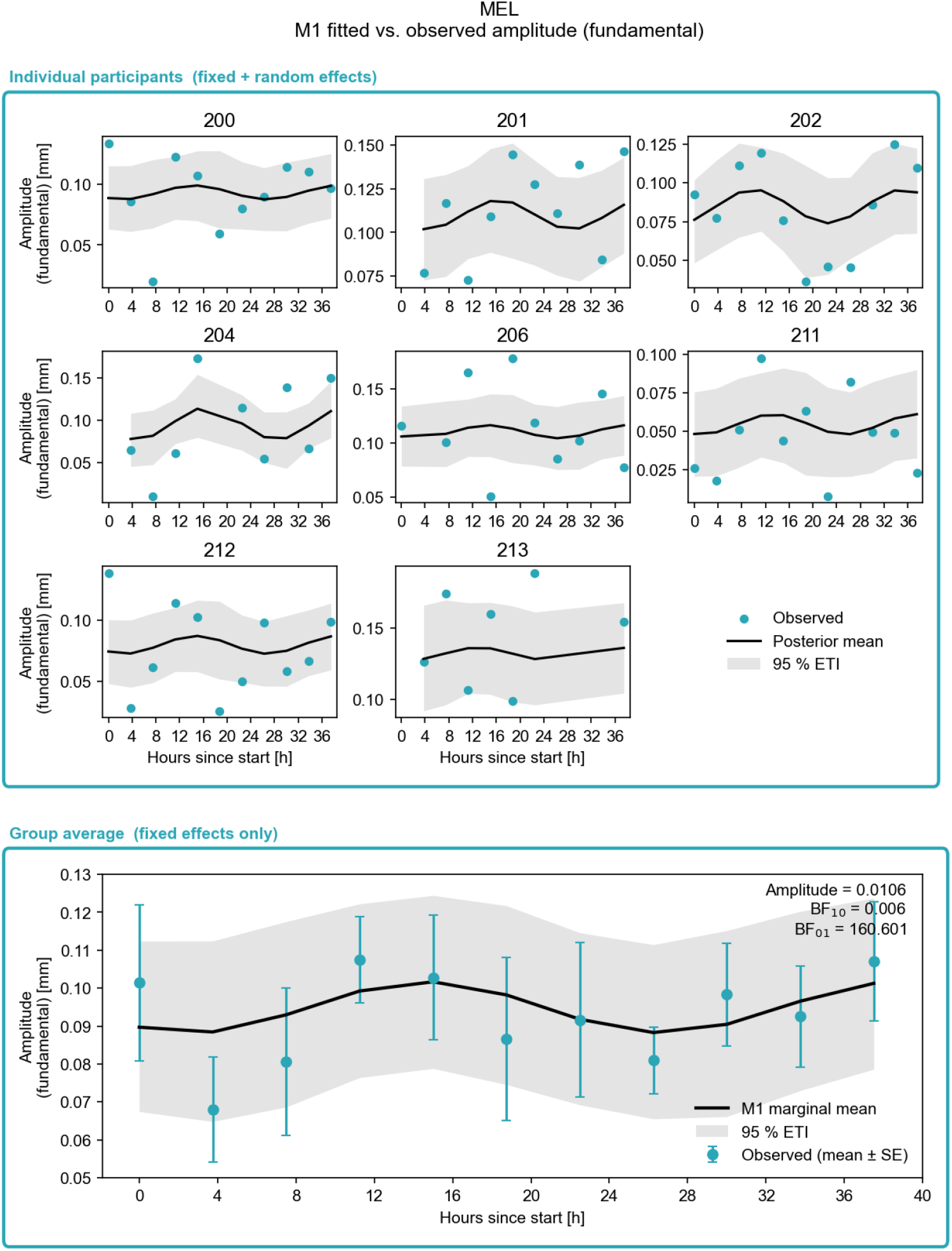
Participant-level M1 fits for phasic amplitude in the MEL condition (80 retained block-level estimates from eight participants). In the participant panels, points are observed estimates, black lines are conditional posterior means, and grey ribbons are 95% equal-tailed intervals. In the group panel, points and error bars are block means and standard errors, the black line is the fixed-effect posterior mean, and the grey ribbon is its 95% equal-tailed interval.

**5—figure supplement 3.**
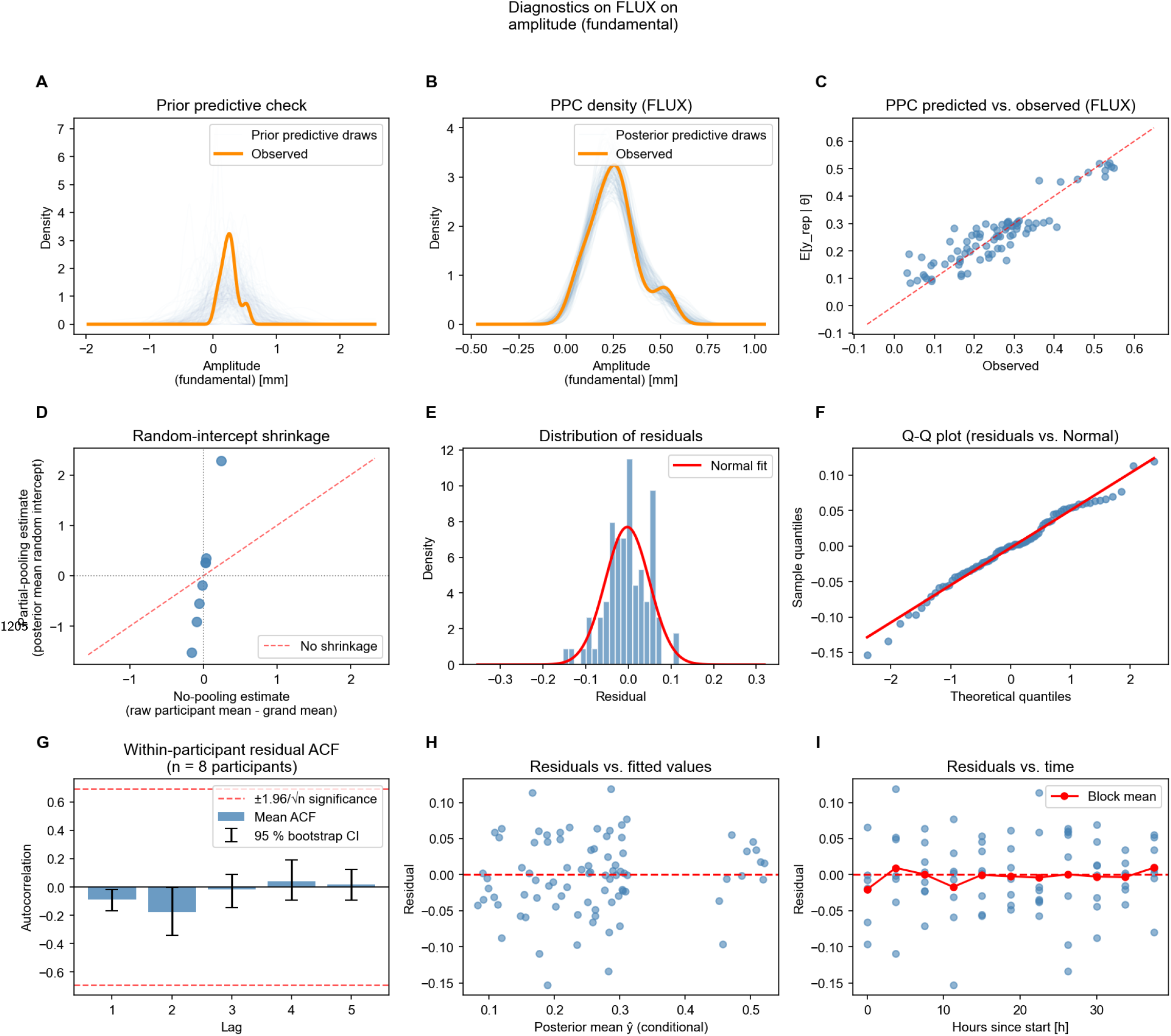
Phasic-amplitude diagnostics for the broadband FLUX condition (83 retained block-level estimates from eight participants). **A**, prior predictive simulations; **B**, posterior predictive distribution; **C**, predicted versus observed values; **D**, participant-level random-intercept shrinkage; **E**, residual distribution; **F**, normal quantile–quantile plot of residuals; **G**, within-participant residual autocorrelation; **H**, residuals versus fitted values; and **I**, residuals versus nominal protocol time.

**5—figure supplement 4.**
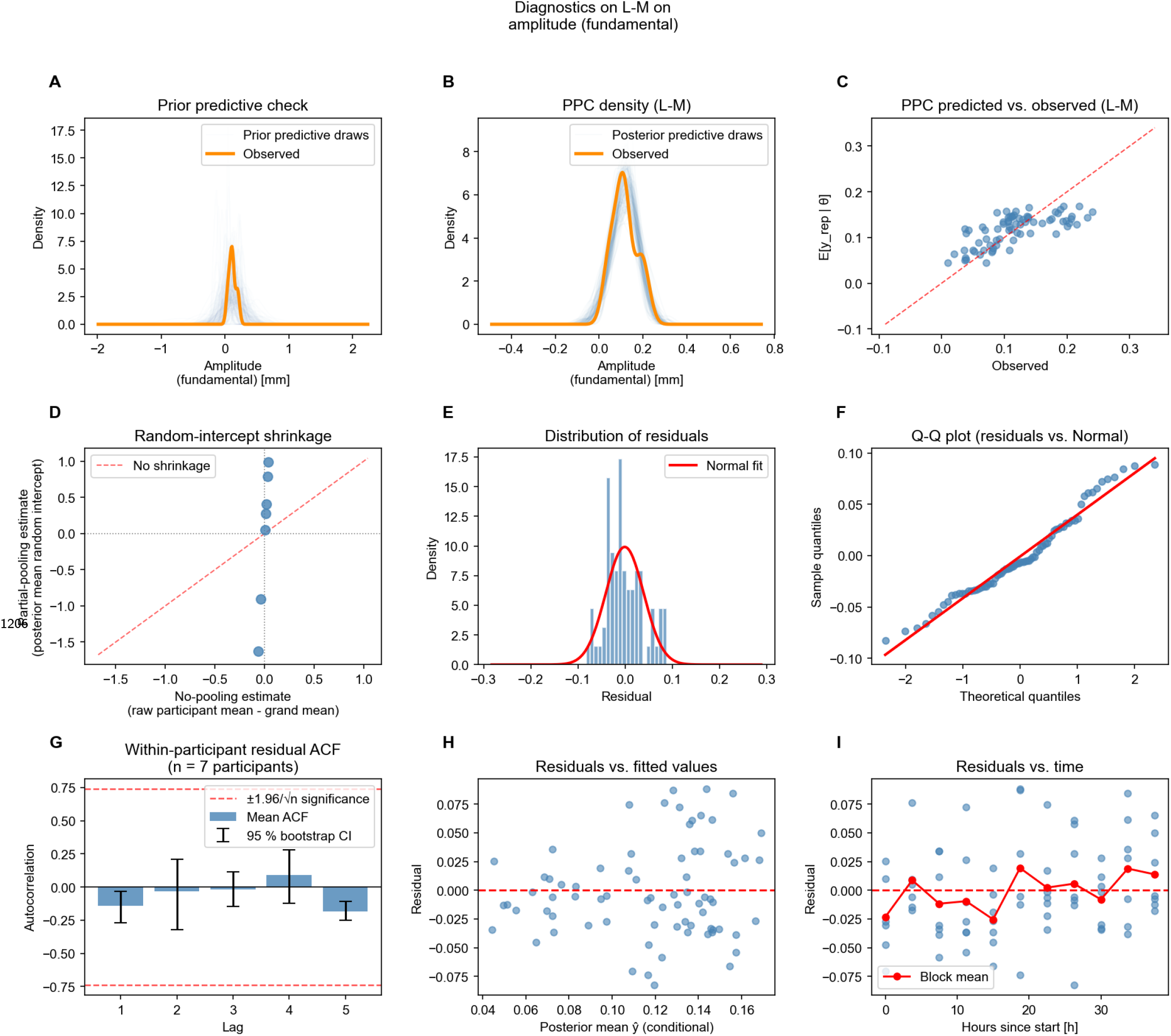
Phasic-amplitude diagnostics for the L–M condition (74 retained block-level estimates from seven participants). **A**, prior predictive simulations; **B**, posterior predictive distribution; **C**, predicted versus observed values; **D**, participant-level random-intercept shrinkage; **E**, residual distribution; **F**, normal quantile–quantile plot of residuals; **G**, within-participant residual autocorrelation; **H**, residuals versus fitted values; and **I**, residuals versus nominal protocol time.

**5—figure supplement 5.**
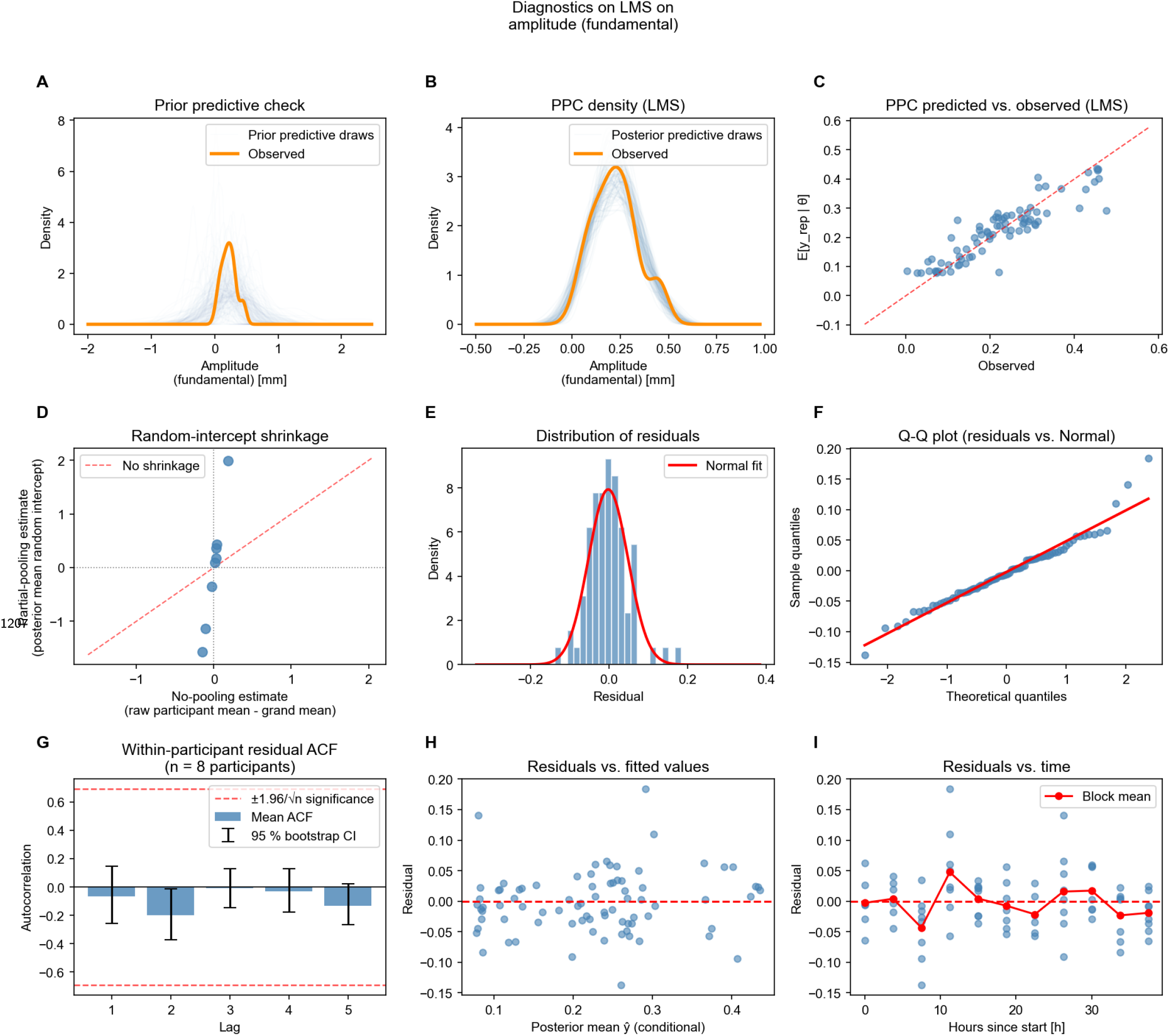
Phasic-amplitude diagnostics for the LMS condition (80 retained block-level estimates from eight participants). **A**, prior predictive simulations; **B**, posterior predictive distribution; **C**, predicted versus observed values; **D**, participant-level random-intercept shrinkage; **E**, residual distribution; **F**, normal quantile–quantile plot of residuals; **G**, within-participant residual autocorrelation; **H**, residuals versus fitted values; and **I**, residuals versus nominal protocol time.

**5—figure supplement 6.**
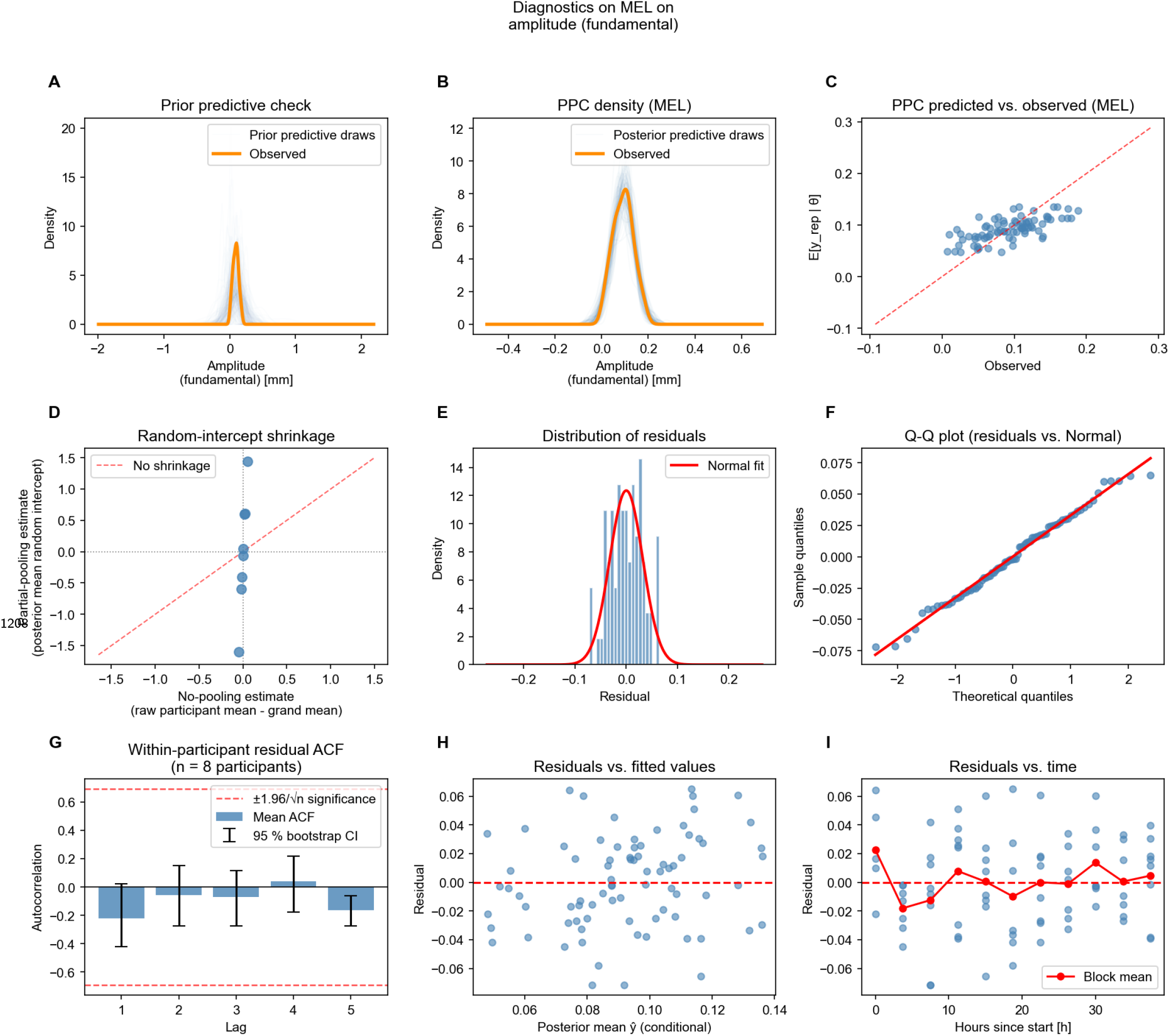
Phasic-amplitude diagnostics for the MEL condition (80 retained block-level estimates from eight participants). **A**, prior predictive simulations; **B**, posterior predictive distribution; **C**, predicted versus observed values; **D**, participant-level random-intercept shrinkage; **E**, residual distribution; **F**, normal quantile–quantile plot of residuals; **G**, within-participant residual autocorrelation; **H**, residuals versus fitted values; and **I**, residuals versus nominal protocol time.

**5—figure supplement 7.**
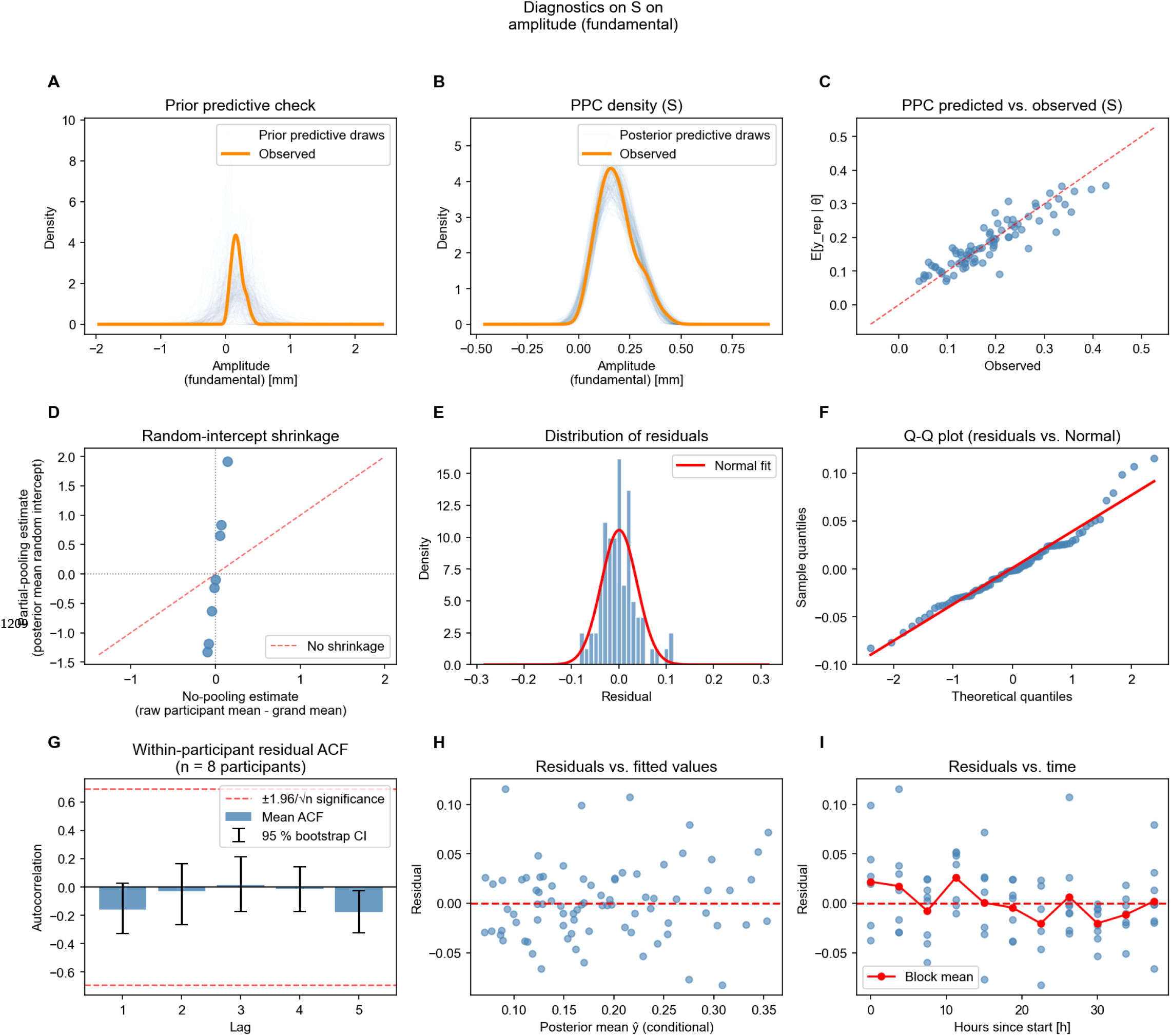
Phasic-amplitude diagnostics for the S condition (81 retained block-level estimates from eight participants). **A**, prior predictive simulations; **B**, posterior predictive distribution; **C**, predicted versus observed values; **D**, participant-level random-intercept shrinkage; **E**, residual distribution; **F**, normal quantile–quantile plot of residuals; **G**, within-participant residual autocorrelation; **H**, residuals versus fitted values; and **I**, residuals versus nominal protocol time.

**6—figure supplement 1.**
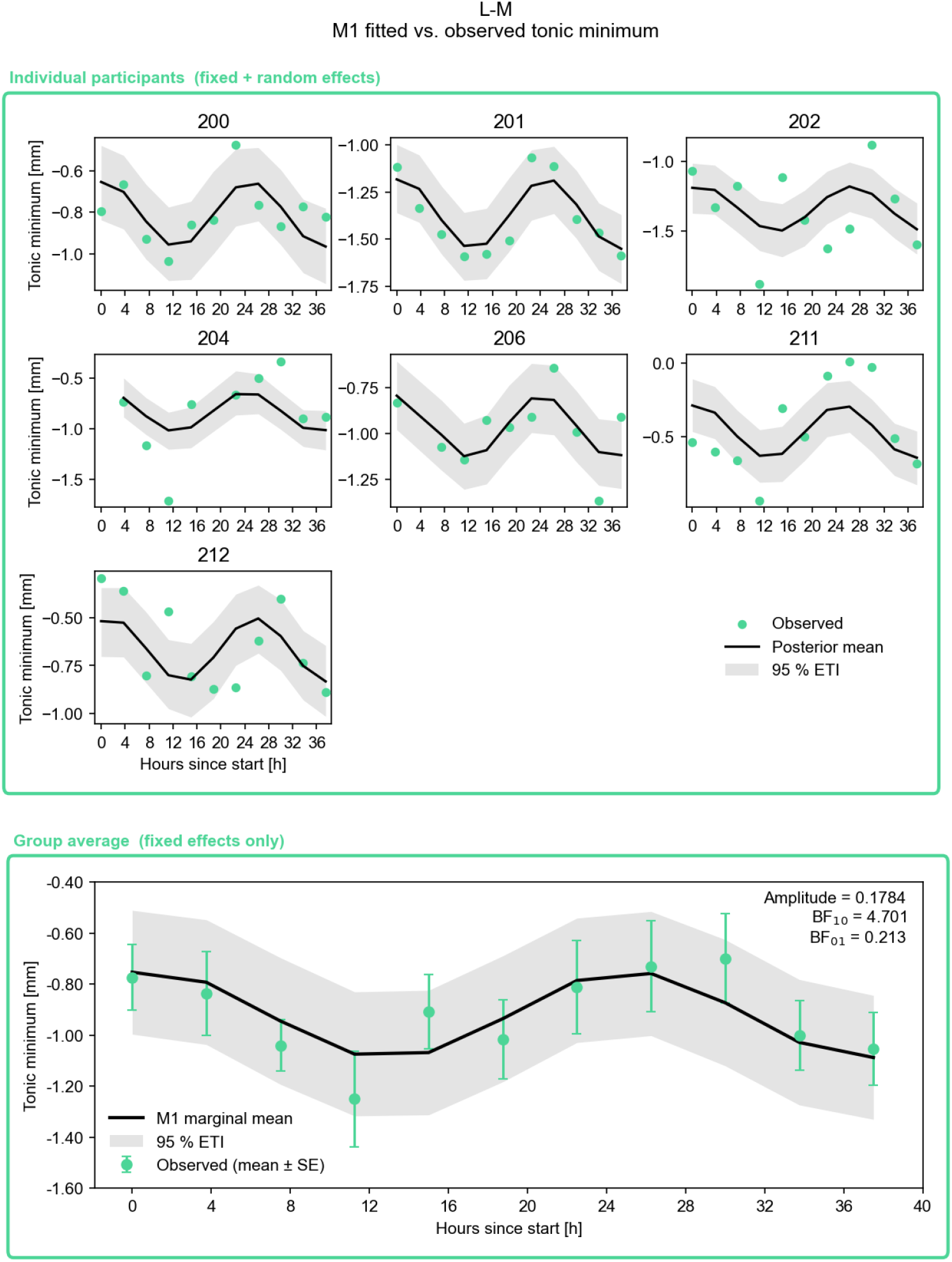
Participant-level M1 fits for tonic minimum in the L–M condition (74 retained block-level estimates from seven participants). In the participant panels, points are observed estimates, black lines are conditional posterior means, and grey ribbons are 95% equal-tailed intervals. In the group panel, points and error bars are block means and standard errors, the black line is the fixed-effect posterior mean, and the grey ribbon is its 95% equal-tailed interval.

**6—figure supplement 2.**
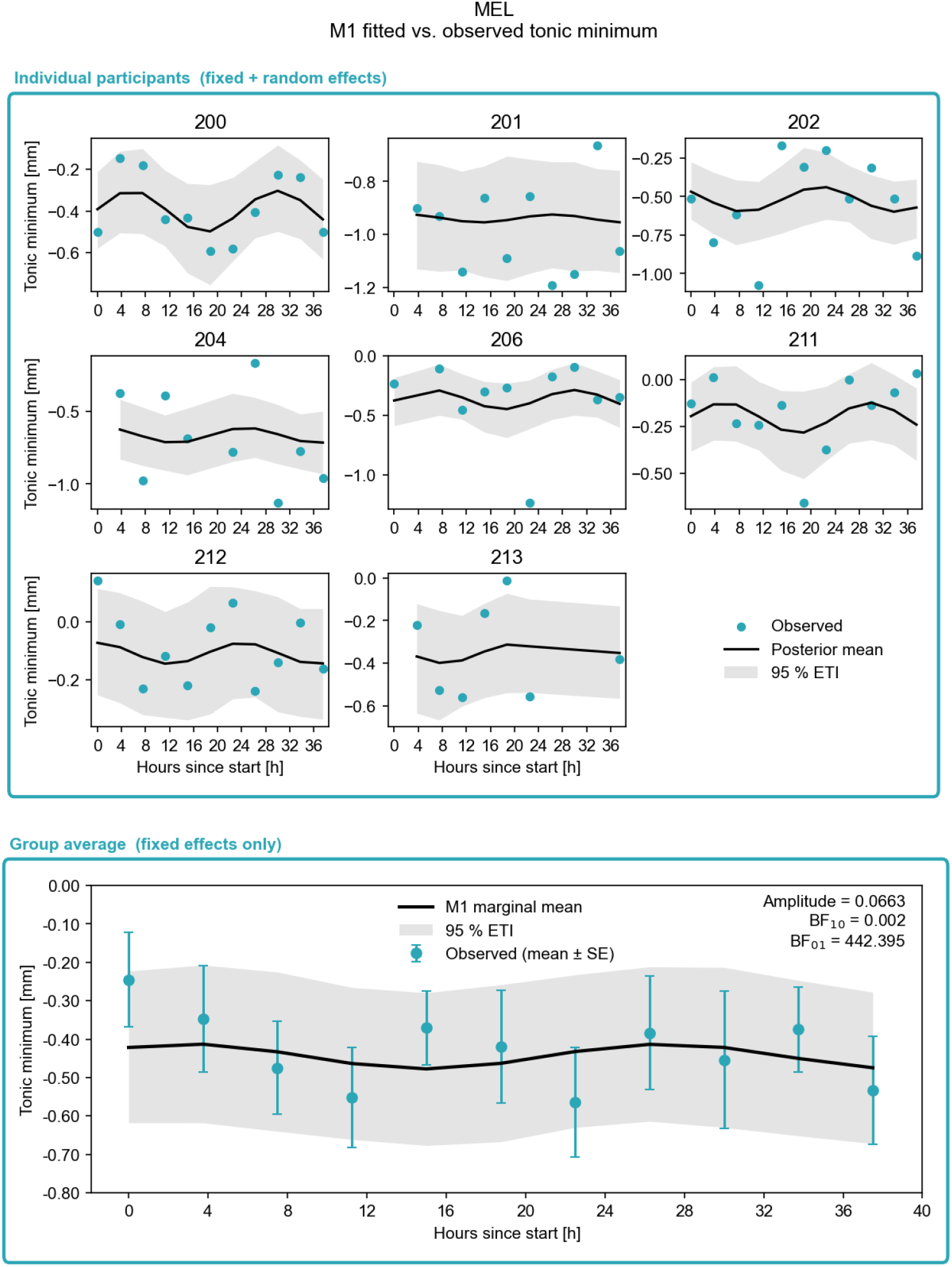
Participant-level M1 fits for tonic minimum in the MEL condition (80 retained block-level estimates from eight participants). In the participant panels, points are observed estimates, black lines are conditional posterior means, and grey ribbons are 95% equal-tailed intervals. In the group panel, points and error bars are block means and standard errors, the black line is the fixed-effect posterior mean, and the grey ribbon is its 95% equal-tailed interval.

**6—figure supplement 3.**
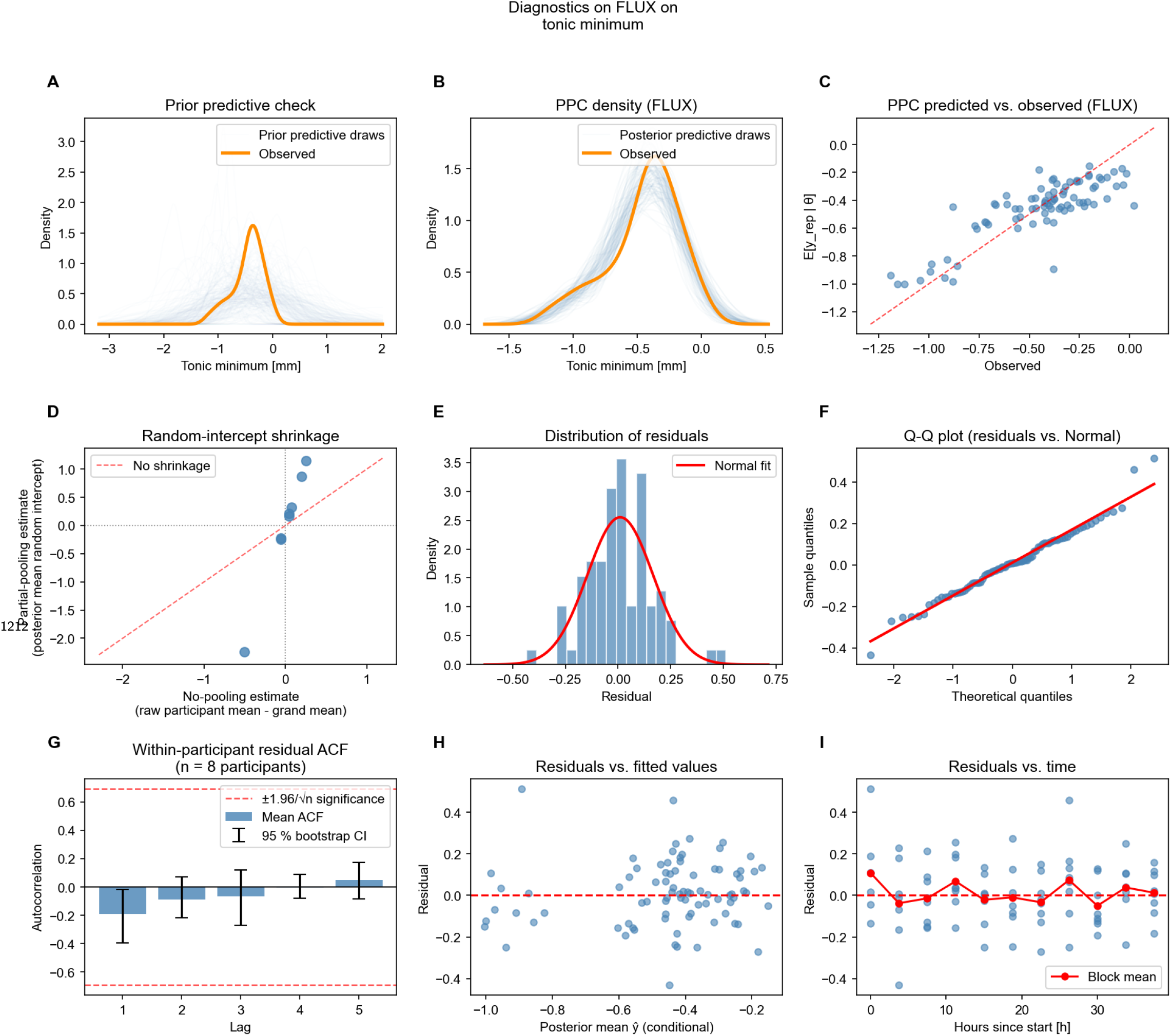
Tonic-minimum diagnostics for the broadband FLUX condition (83 retained block-level estimates from eight participants). **A**, prior predictive simulations; **B**, posterior predictive distribution; **C**, predicted versus observed values; **D**, participant-level random-intercept shrinkage; **E**, residual distribution; **F**, normal quantile–quantile plot of residuals; **G**, within-participant residual autocorrelation; **H**, residuals versus fitted values; and **I**, residuals versus nominal protocol time.

**6—figure supplement 4.**
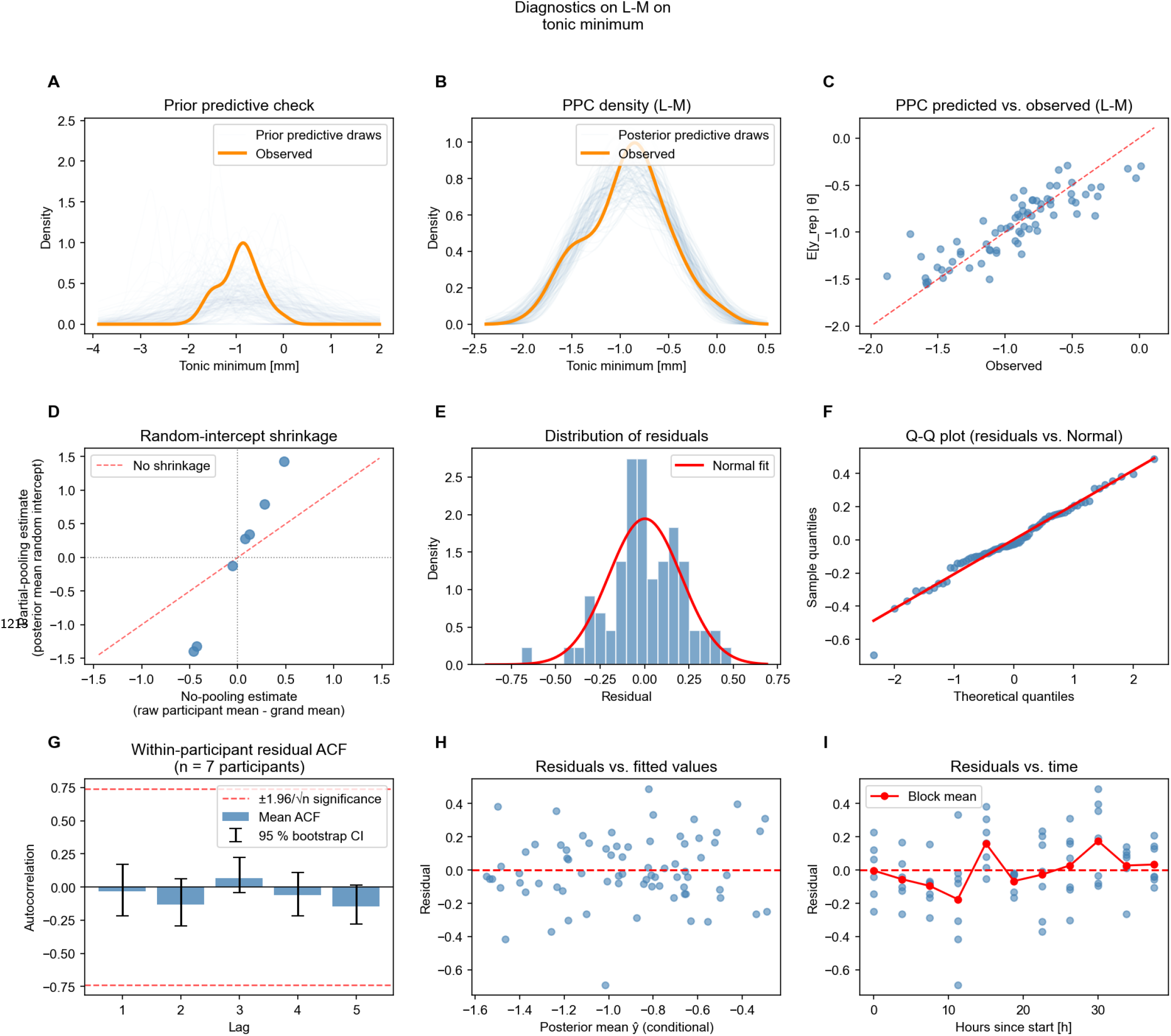
Tonic-minimum diagnostics for the L–M condition (74 retained block-level estimates from seven participants). **A**, prior predictive simulations; **B**, posterior predictive distribution; **C**, predicted versus observed values; **D**, participant-level random-intercept shrinkage; **E**, residual distribution; **F**, normal quantile–quantile plot of residuals; **G**, within-participant residual autocorrelation; **H**, residuals versus fitted values; and **I**, residuals versus nominal protocol time.

**6—figure supplement 5.**
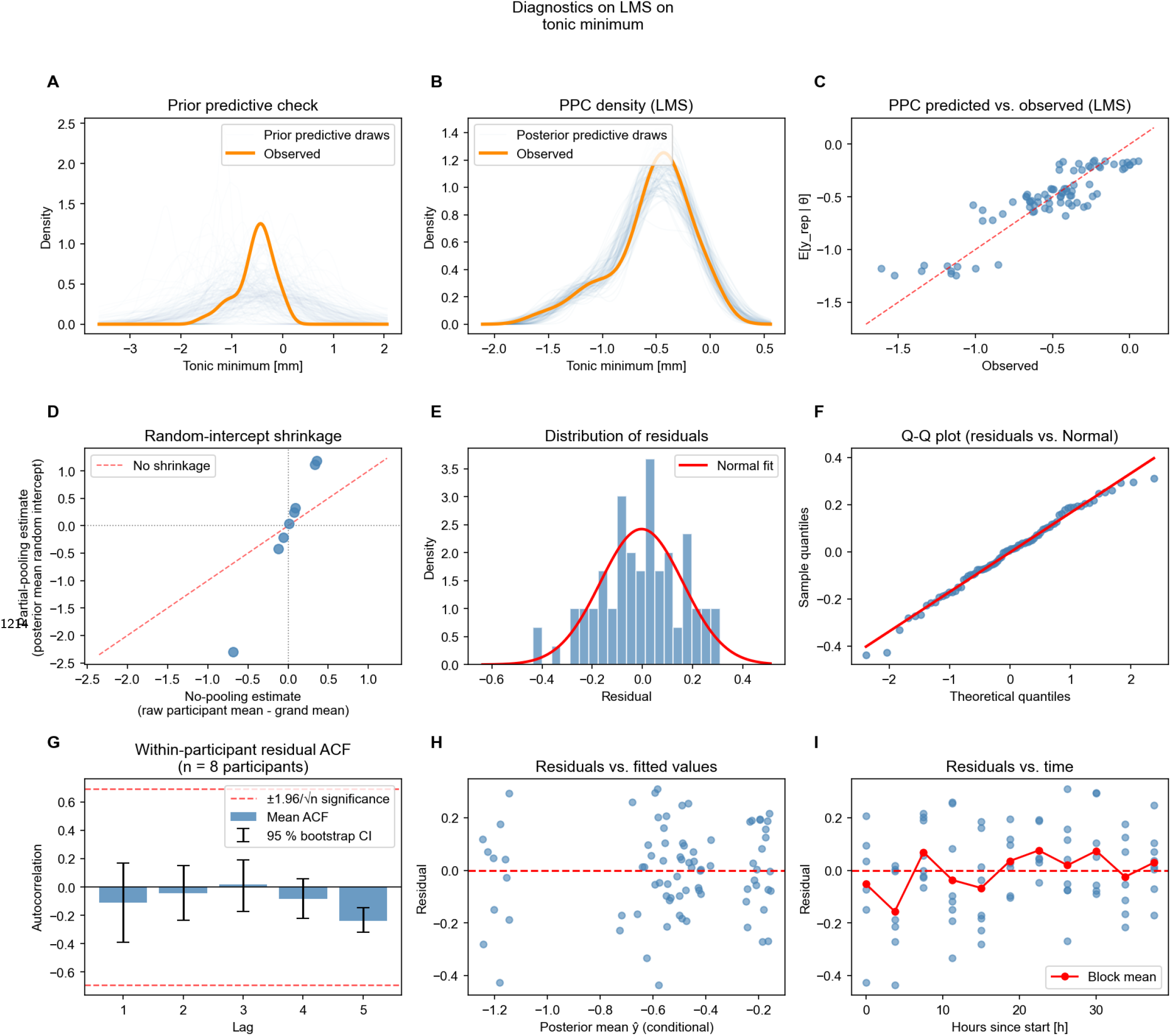
Tonic-minimum diagnostics for the LMS condition (80 retained block-level estimates from eight participants). **A**, prior predictive simulations; **B**, posterior predictive distribution; **C**, predicted versus observed values; **D**, participant-level random-intercept shrinkage; **E**, residual distribution; **F**, normal quantile–quantile plot of residuals; **G**, within-participant residual autocorrelation; **H**, residuals versus fitted values; and **I**, residuals versus nominal protocol time.

**6—figure supplement 6.**
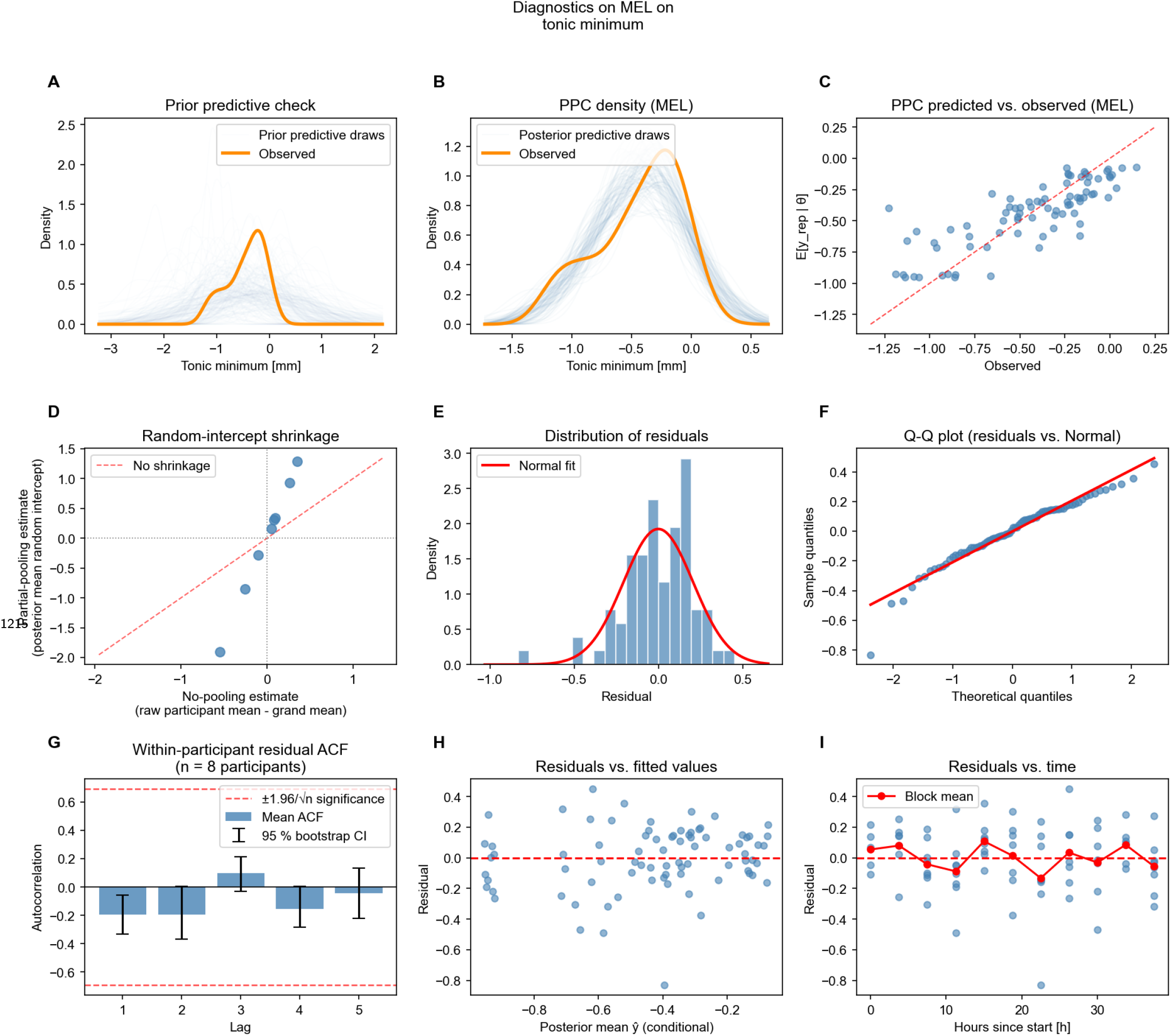
Tonic-minimum diagnostics for the MEL condition (80 retained block-level estimates from eight participants). **A**, prior predictive simulations; **B**, posterior predictive distribution; **C**, predicted versus observed values; **D**, participant-level random-intercept shrinkage; **E**, residual distribution; **F**, normal quantile–quantile plot of residuals; **G**, within-participant residual autocorrelation; **H**, residuals versus fitted values; and **I**, residuals versus nominal protocol time.

**6—figure supplement 7.**
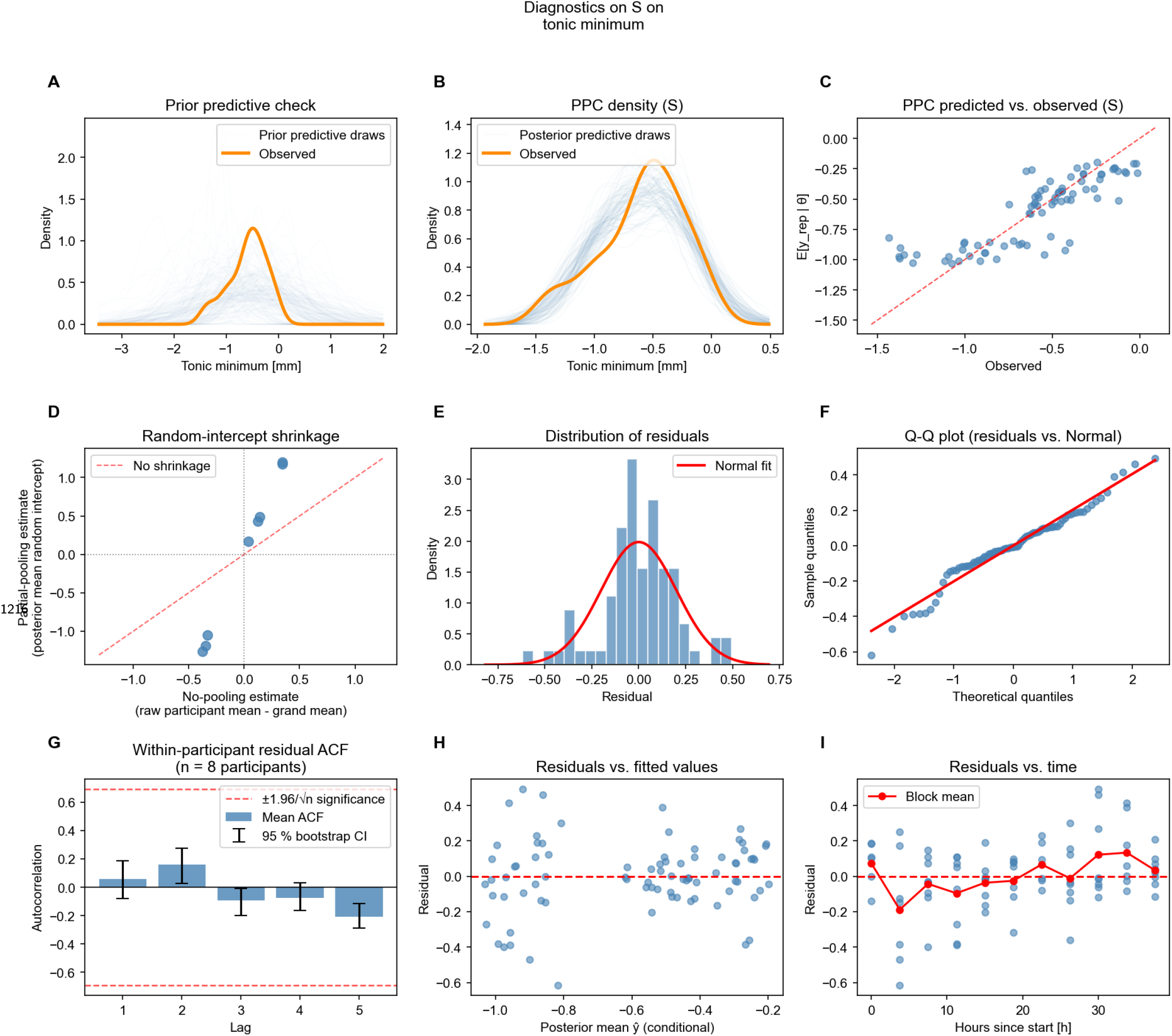
Tonic-minimum diagnostics for the S condition (81 retained block-level estimates from eight participants). **A**, prior predictive simulations; **B**, posterior predictive distribution; **C**, predicted versus observed values; **D**, participant-level random-intercept shrinkage; **E**, residual distribution; **F**, normal quantile–quantile plot of residuals; **G**, within-participant residual autocorrelation; **H**, residuals versus fitted values; and **I**, residuals versus nominal protocol time.

